# Spatiomolecular mapping reveals anatomical organization of heterogeneous cell types in the human nucleus accumbens

**DOI:** 10.1101/2025.09.10.675374

**Authors:** Prashanthi Ravichandran, Svitlana V. Bach, Robert A. Phillips, Madeline R. Valentine, Nicholas J. Eagles, Yufeng Du, Ishbel del Rosario, Ryan A. Miller, Heena R. Divecha, Madhavi Tippani, Kelsey D. Montgomery, Joel E. Kleinman, Shizhong Han, Stephanie C. Page, Thomas M. Hyde, Leonardo Collado-Torres, Alexis Battle, Keri Martinowich, Stephanie C. Hicks, Kristen R. Maynard

## Abstract

The nucleus accumbens (NAc) is a key component of the mesolimbic dopamine system that critically regulates many behaviors related to reward and motivation. The NAc is implicated in several neuropsychiatric disorders, including major depressive disorder, schizophrenia, and substance use disorders. Rodent studies have identified spatial organization of heterogeneous medium spiny neuron (MSN) subtypes across the NAc core and shell, but the extent to which this cellular diversity and spatial organization is conserved in the human brain remains unclear. Here, we generated a spatiomolecular atlas of NAc cell types and spatial domains by integrating spatial transcriptomics and single-nucleus RNA sequencing data from postmortem NAc tissue from 10 neurotypical adult donors. We identified 20 transcriptionally unique cell populations and 8 spatial domains, including specialized D1 islands composed of distinct dopamine receptor 1 (DRD1) MSN subtypes, which were enriched for *OPRM1*. In contrast to a discrete core vs. shell division, we observed continuous spatial gradients of gene expression across MSN domains, suggesting a more complex organization of DRD1 and DRD2 MSNs. Cross-species comparisons demonstrated conservation of MSN subtypes and spatial features between human, rodent, and nonhuman primate NAc. Genetic enrichment analysis with stratified linkage disequilibrium score regression revealed specific spatial domains associated with risk for psychiatric and addiction-related traits. To investigate this further, we spatially mapped ligand-receptor interactions involving neuropsychiatric risk genes. Finally, we leveraged existing rodent NAc data to identify drug-responsive transcriptional programs and predict their spatial distribution in the human NAc. Collectively, we provide a spatiomolecular framework for understanding the human NAc and its relevance to neuropsychiatric disease.

## 1 Introduction

The nucleus accumbens (NAc) is in the ventral part of the striatum and plays a central role in motivation, reward processing, and goal-directed behavior^1–3^. NAc dysfunction has been associated with several neuropsychiatric disorders, including schizophrenia (SCZ), depression, anxiety, and substance use disorders^4–8^. The NAc is a key node of the mesolimbic reward pathway, integrating glutamatergic input from cortical, thalamic, and limbic structures as well as dopaminergic input from the ventral tegmental area (VTA)^9,10^. GABAergic medium spiny neurons (MSNs) constitute the main output neurons of the NAc, and are divided into two major types: DRD1 MSNs expressing dopamine receptor D1 (*DRD1*) and DRD2 MSNs expressing *DRD2*, each having unique connectivity patterns with downstream subcortical targets^11–13^. Classically, DRD1 MSNs form the direct pathway, while DRD2 MSNs form the indirect pathway enabling activity in these circuits to promote opposite effects on cortical regions^11^. While MSNs account for over 90% of neurons in the NAc, several other GABAergic or neuropeptide-expressing populations play a role in modulating MSN function, including neurons expressing acetylcholine, parvalbumin, and somatostatin^14–16^.

Mounting evidence suggests that NAc cell types show extensive molecular, morphological, and functional diversity enabling complex circuit-specific behaviors^1,17^. Single-cell and single-nucleus RNA-sequencing (snRNA-seq) studies have identified molecularly distinct subpopulations of DRD1 and DRD2 MSNs in rodents^18–23^, non-human primates (NHPs)^24^, and in humans^25^. These MSN subpopulations show unique transcriptional responsiveness to drugs of abuse, including cocaine and morphine^20,26–28^, and differential enrichment of genes associated with risk for neuropsychiatric disorders and substance use phenotypes^25^. Spatial transcriptomic studies in rodents have further characterized the anatomical diversity of transcriptionally-distinct MSN subtypes across the anterior-posterior axis of the NAc^23^, and provided evidence for a spatiomolecular code across the striatum beyond cell type organization^29^. While the molecular anatomy of the NAc has been studied in great detail in rodents, it is less understood whether the same spatiomolecular architecture and cell type organization exists in the human NAc. For example, it is unclear whether the human NAc exhibits a spatially and cellularly distinct core and shell region as is described in the rodent and NHP NAc^23,30–32^.

One prominent anatomical feature of the human NAc is the presence of interface islands located along the ventral and medial border of the NAc^33,34^. These islands have also been characterized in rodents and NHPs and consist exclusively of DRD1 MSN subtypes, a subset of which show high expression of the μ-opioid receptor (*OPRM1*)^24,35^. The spatial location of these D1 islands overlaps with hotspots in the NAc shell that mediate the effects of opioids and hedonic reactions to palatable rewards^36^. While putative D1 island MSN subtypes have been identified in the human brain using snRNA-seq^25^, the cellular composition and spatiomolecular identity of interface islands has not yet been directly studied in the human NAc.

We generated an integrated single-nucleus and spatial transcriptomic atlas of the human NAc to better understand the anatomical organization of heterogeneous MSN cell types. We identified multiple transcriptionally-distinct populations of DRD1 and DRD2 MSNs that were organized in continuous gradients across the medial-lateral axis of the NAc. Consistent with previous work in rodents and NHPs, we identified *OPRM1*-expressing D1 interface islands along the medial border of the NAc, which were composed of two specific DRD1 MSN subpopulations. Using snRNA-seq data, we identified ligand-receptor pairs associated with substance dependence and neuropsychiatric disorders such as depression, anxiety, and SCZ. We then spatially localized these interacting cell types to discrete locations throughout the NAc. Finally, using non-negative matrix factorization (NMF) and topic modeling, we identified gene expression patterns associated with morphine and cocaine intake in rodents and demonstrated distinct spatial patterns of transcriptional response. We created interactive web resources for the scientific community to explore our integrated dataset and further advance understanding of reward circuitry in neuropsychiatric disorders and addiction.

## 2 Results

### 2.1 Cellular and spatiomolecular architecture of the human NAc

We defined the spatiomolecular landscape of the nucleus accumbens (NAc) in the adult human brain, by generating paired spatially-resolved (SRT) and single nucleus (snRNA-seq) transcriptomics data with the 10x Genomics Visium and Chromium platforms, from adjacent tissue sections of 10 adult neurotypical brain donors (6 male, 4 female) (**Fig. 1A, Table S1, Methods**). While we aimed to sample the NAc at the midpoint along the anterior-posterior (A-P) axis, anatomical alignment revealed that the sections were obtained from the middle (intermediate) as well as slightly anterior and posterior positions (**Fig. S1**). By documenting the A-P location of each section, we were able to gain insights into the molecular composition of the NAc along this neuroanatomical axis (**Fig. S2, Fig. S3, Fig. S4**). To profile the entire mediolateral and dorsoventral span of the NAc, we scored tissue blocks and profiled tissue subsections across multiple capture arrays (**Fig. 1A**). We used 2-5 capture arrays per donor (n = 38 capture areas total) and reconstructed the entire structure of the NAc by aligning capture arrays using shared histological landmarks found in overlapping regions^37^ (**Fig. 1A, Fig. S5, Fig. S6, Methods**). For each donor, we obtained nuclear sorted (PI+) and neuron-enriched (PI+NeuN+) samples from adjacent tissue sections (n=20) to increase the capture of heterogeneous MSNs and rare neuron populations.

**Fig. 1.**
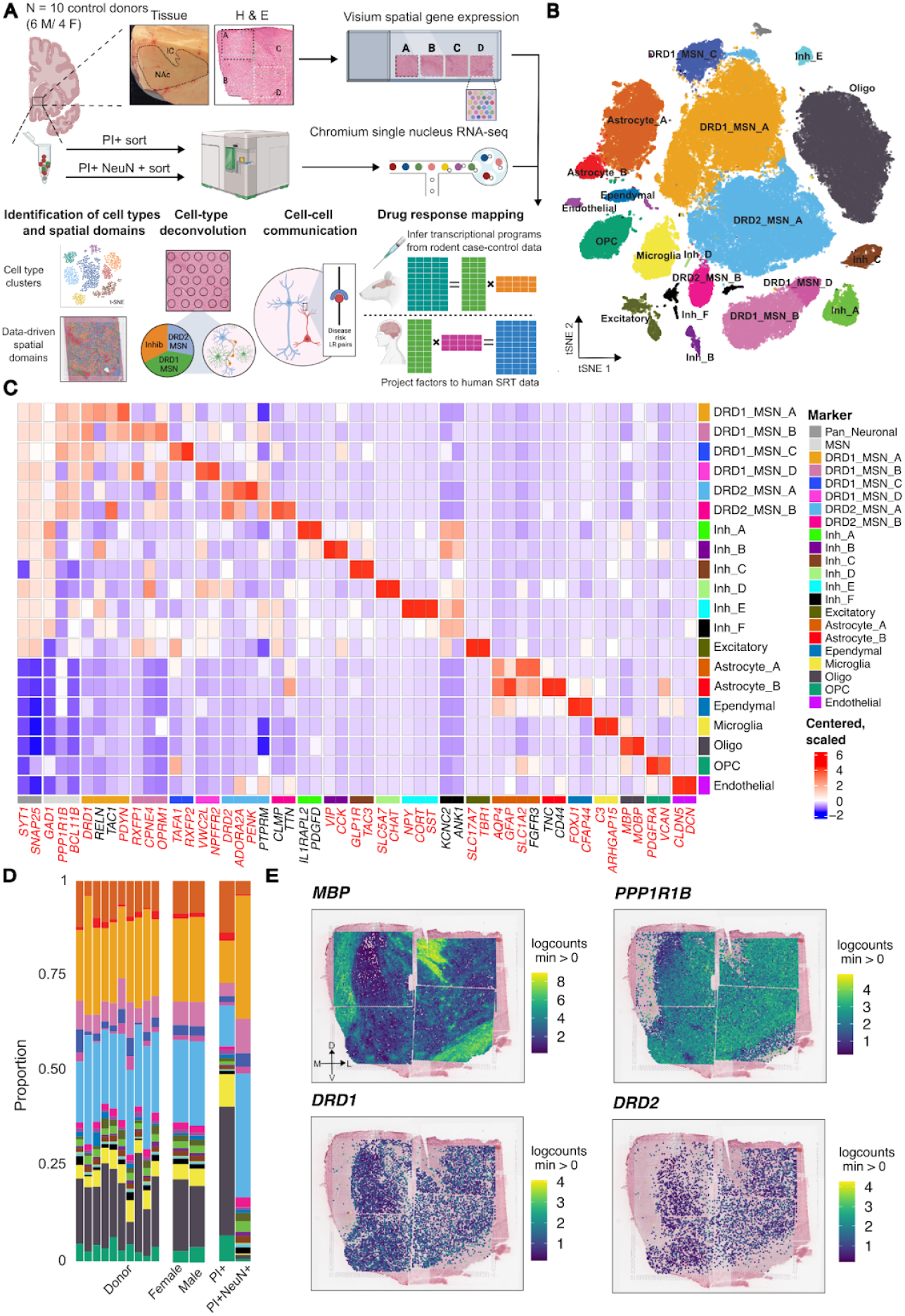
Cellular diversity and reconstruction of spatial gene expression in the human nucleus accumbens (NAc) (**A**) Tissue blocks containing the NAc were dissected from 10 neurotypical adult donors (6 male, 4 female). Paired single-nucleus RNA sequencing (snRNA-seq) and spatially-resolved transcriptomics (SRT) data were generated from adjacent tissue sections using 10x Genomics Chromium and Visium platforms. To capture the complete NAc, tissue blocks were scored to align with the width of the Visium capture array and spatial profiling was performed across 2–5 capture arrays per donor (n = 38 total). snRNA-seq was performed on the same tissue blocks on PI sorted (PI+) and neuron-enriched (PI+ NeuN+) nuclei. Downstream analyses included (i) Identification of transcriptionally distinct cell clusters from snRNA-seq, (ii) Mapping spatial domains in SRT data, (iii) Integration of snRNA-seq and SRT data via cell-type deconvolution to resolve spatially-localized populations, (iv) Inference of disease relevant ligand-receptor (LR)–based cell–cell communication networks, and (v) Cross-species drug-response mapping by integrating transcriptional programs derived from rodent datasets with human SRT data. (**B**) *t-*distributed stochastic embedding (t-SNE) summarizing 20 transcriptionally distinct cell populations, including subtypes of DRD1 and DRD2 medium spiny neurons (MSNs), inhibitory neurons, and non-neuronal glial cell types. (**C**) Heatmap of centered and scaled, cluster-averaged expression values for canonical and cluster-derived marker genes, computed from log-normalized counts (logcounts). (**D**) Differences in cell type proportions across donors, sexes, and sorting paradigms, showing that NeuN+ enrichment substantially increases the detection of neuronal populations compared to PI+ samples. (**E**) Spatial gene expression of *MBP* (oligodendrocyte marker), *PPP1R1B* (MSN marker), *DRD1*, and *DRD2* which define MSN subtypes demonstrating clear boundaries between white matter and grey matter, and confirming the presence of both DRD1 and DRD2 MSNs across the NAc.

Following standard pre-processing and quality control (QC) workflows for snRNA-seq data (**Fig. S7, Methods**), we retained 103,785 high-quality nuclei. Using unsupervised clustering and annotations based on established marker genes, we identified 20 transcriptionally distinct cell types including 13 neuronal and 7 non-neuronal populations (**Fig. 1B-C**). We observed similar proportions of cell types across donors and sex with more neuronal nuclei captured in NeuN+ sorted samples as expected (**Fig. 1D**). Neuronal clusters exhibited high expression of expected marker genes, with a large proportion of neurons expressing either *DRD1* or *DRD2*. Non-neuronal clusters included astrocytes, oligodendrocytes, oligodendrocyte precursor cells (OPCs), ependymal cells, endothelial cells, and microglia (**Fig. S8**).

Consistent with estimates that GABAergic MSNs constitute up to 95% of neurons in the NAc^23–26^, almost all neuronal populations exhibited high expression of *GAD1* (**Fig. S8**) and *PPP1R1B*, which encodes the canonical MSN marker, DARPP-32 (**Fig. S8**). In addition to *PPP1R1B*, expression of *BCL11B*^38^, *PDE10A*^39^, *and FOXP2*^20,40^ *was used to classify neurons as MSNs (**Fig. S8**). MSNs were further classified based on expression of DRD1* or *DRD2* (**Fig. S8**), which showed broad expression across the extent of the NAc (**Fig. 1E**). We identified four populations of *DRD1*-expressing MSNs (DRD1_MSN_A - DRD1_MSN_D) and two populations of *DRD2*-expressing MSNs (DRD2_MSN_A, DRD2_MSN_B), each with distinct transcriptional profiles (**Table S2**). DRD1_MSN_A and DRD2_MSN_A expressed *CALB1*, a marker of core-enriched MSNs in the rat NAc^30^. Cross-species comparison of cell type-specific gene signatures revealed that these subtypes corresponded to the D1 and D2 Matrix populations in non-human primate (NHP)^24^ (**Fig. S9, Fig. S10**). In contrast, DRD2_MSN_B was associated with the NHP D2 striosome cell type and expressed *CLMP* and *TTN*. DRD1_MSN_C showed high *RXFP2* expression and corresponded to a previously identified population of *RXFP2*+ DRD1-MSNs in humans^25^. Additionally, this population was most transcriptionally similar to D1 Shell/OT and the D1 Islands of Calleja (ICj) cell types in NHP^24^. Finally, we identified two populations, DRD1_MSN_B and DRD1_MSN_D, which expressed *RXFP1* and *CPNE4* and were closely associated with Grm8 MSNs in the rodent^26^ and D1 neurochemically unique domains in the accumbens and Pt (NUDAP) in NHP^24^. Interestingly, these cell types in the rodent and NHP localized to D1 islands, which have a distinct spatial topography, suggesting that this compartmentalized MSN population is a conserved feature of striatal organization across species. We observed high expression of the corticotrophin releasing hormone receptor 2 (*CRHR2)* in DRD1_MSN_B (**Fig. S8**), which is consistent with our previous findings showing *CRHR2* expression in *RXFP1*+ DRD1 MSN populations^25^. DRD1_MSN_D showed greater expression of *CRHR1* compared to *CRHR2*.

Beyond MSNs, we also identified several diverse populations of neurons expressing *GAD1*, including a population of *CHAT+* cholinergic neurons (Inh_D; **Fig. 1C**), which play a critical role in regulating drug-induced behavioral adaptations^41–44^. Additionally, we observed *KIT*+ parvalbumin neurons^45^ (Inh_A), *SST*+/*CORT*+ neurons (Inh_E)^46^, and *VIP+* neurons (Inh_B)^47^ (**Fig. 1C**). *SST*+/*CORT*+ neurons showed high expression of *CRHR2* (**Fig. S8**). In addition to these well characterized neuron populations, we identified a previously described population^25,48^ of *GLP1R+/TAC3+* GABAergic neurons (Inh_C; **Fig. 1C**). Finally, we found a distinct subpopulation of inhibitory neurons, Inh_F, which expressed *KCNC2* and *ANK1*, but did not express canonical marker genes corresponding to previously described inhibitory populations in the NAc.

We next investigated whether transcriptionally distinct NAc cell types were enriched for the heritability of complex traits using stratified LD score regression (s-LDSC)^49^ (**Methods**). We evaluated 20 complex traits spanning psychiatric disorders, substance use, and cognitive performance. As expected, we observed enrichment of Alzheimer’s Disease heritability in microglia (**Fig. S11, Table S3**). In addition, we found significant associations (FDR < 0.1) between drinks per week and DRD1_MSN_A and DRD2_MSN_A, and between depression and the Inh_F and Inh_B neuron populations (**Fig. S11**). These results extend previous findings that broadly implicate MSNs in alcohol-related traits^50^ by demonstrating that heritability is enriched in specific MSN cell types and inhibitory neuron populations.

### 2.2 Identification of molecularly-defined spatial domains in the human NAc

To map spatial gene expression across the NAc, we aligned Visium capture areas within each donor to reconstruct contiguous tissue sections^37^ (**Fig. S5, Fig. S6, Methods**). After QC (**Fig. S12, Fig. S13, Fig. S14 Fig. S15**), we retained 176,013 high-quality spots across 10 donors, with a median of 18,536 spots per donor. Controlling for broad differences between white and gray matter (**Fig. S16**), we identified 2,000 spatially variable genes (**Table S4**)^51^ as input for spatial clustering using *PRECAST*^*52*^. We evaluated potential clustering resolutions from *k* = 3 to 15 and selected *k* = 10 based on the Bayesian Information Criterion (BIC)^53^ (**Fig. S17**) and interpretability of differentially expressed genes (DEGs) across clusters. We annotated these clusters based on canonical marker genes^23–25,31,33,54^. To improve biological interpretability, we merged clusters 6 and 10 into a shared endothelial and ependymal domain based on expression of vascular and choroid plexus markers, including *CLDN5* and *TTR* (**Table S5**). We also merged clusters 2 and 9 into a white matter (WM) domain based on expression of oligodendrocyte and myelination-associated genes such as *MBP, MOBP*, and *PLP1* (**Table S5**). This yielded eight spatial domains (SpDs): MSN_1, MSN_2, MSN_3, Excitatory, Inhibitory, D1 islands, Endothelial/Ependymal, and White matter (WM) (**Fig. 2A**).

**Fig. 2.**
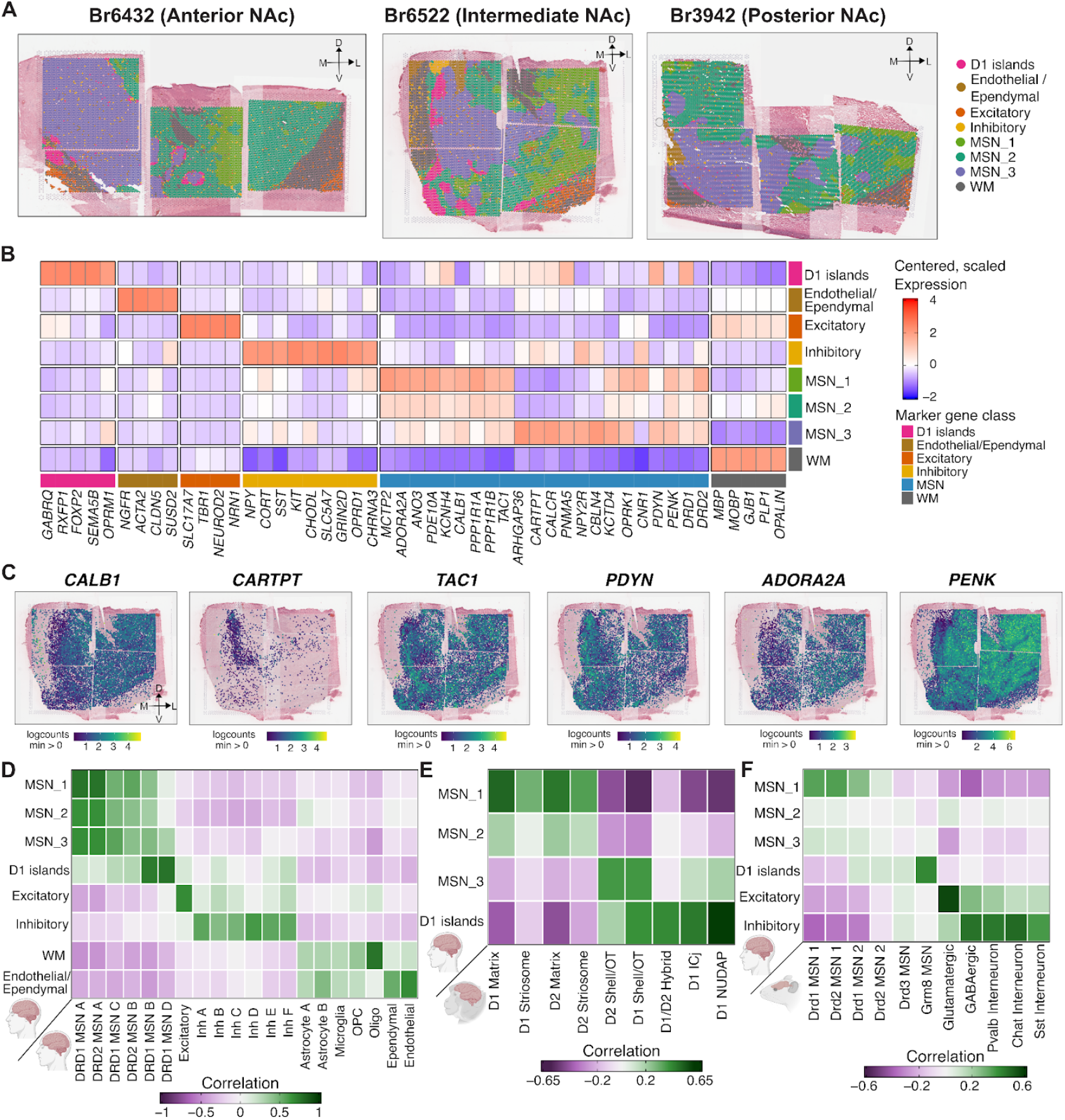
Transcriptionally distinct spatial domains (SpDs) in the human NAc. (**A**) Representative spatial domains (SpDs) across three representative donors spanning anterior (Br6432), intermediate (Br6522), and posterior (Br3942) positions along the anterior-posterior (A-P) axis. Eight spatial domains were identified with varying prevalence across the A-P axis (e.g., D1 islands are prominent in anterior and intermediate NAc, but nearly absent in posterior NAc). (**B**) Scaled and centered expression heatmap of selected marker genes across the 8 spatial domains. For each domain, log-normalized gene expression was averaged across all spots within that domain, followed by z-score transformation. (**C**) Spatial localization of select marker genes illustrating domain-specific and medial-lateral differences in gene expression. *ADORA2A* and *PDE10A* are expressed in the lateral NAc and MSN_1, while *TAC3, ARHGAP36*, and *CARTPT* show higher expression in the medial NAc and MSN_3. (**D**) Correlation of *t*-statistics derived from DEG analysis of SpDs and paired human snRNA-seq data. D1 islands are transcriptionally similar to DRD1_MSN_B and DRD1_MSN_D populations, while MSN_1 SpD is most transcriptionally similar to DRD1_MSN_A and DRD2_MSN_A. (**E**) Correlation of *t*-statistics derived from DEG analysis of human SpDs and NHP MSN subtypes ^24^. D1 islands map to specialized populations including D1 ICj and D1 NUDAP, MSN_1 SpD and MSN_2 SpD resemble core-biased NHP MSNs, and MSN_3 SpD maps to shell-associated NHP MSNs. (F) Correlation of *t*-statistics derived from DEG analysis of human SpDs and rodent MSN subtypes ^26^. D1 islands map to Grm8 MSNs in the rodent and MSN_1 is enriched for DRD1_MSN_1 and DRD2_MSN_1.

These SpDs were consistently identified across all samples, although their prevalence varied across anterior, intermediate, and posterior positions (**Fig. 2A**), reflecting known anatomical and functional heterogeneity along the A-P axis^23^. For example, D1 islands were found in anterior (Br6432) and intermediate (Br6522) positions and nearly absent in posterior (Br3942) positions. Further, D1 islands showed a distinct transcriptional signature, including expression of *TRHDE, CPNE4, FOXP2, GABRQ, DRD1*, and *RXFP1* (**Fig. 2B**). Consistent with these findings, previous studies in NHPs demonstrated that *RXFP1* and *CPNE4* selectively label distinct, DRD1-exclusive cell islands in the ventral striatum^24^. Further, concordant with the description of D1 islands in rodents^35^ and NHP^24^, we observed high levels of *OPRM1* and low levels of *OPRK1* in human D1 islands (**Fig. 2B**).

Beyond D1 islands, we identified three other transcriptionally distinct MSN SpDs: MSN_1, MSN_2, and MSN_3. Compared to the sharp spatial boundaries of the D1 islands, these MSN SpDs exhibited gradient-like transitions. Across multiple random initializations, *PRECAST* generated variable boundaries between these SpDs (**Fig. S18**), and the DEGs differentiating SpDs MSN_1-3 displayed smaller log_2_-fold changes (logFC) (**Table S6**). Nonetheless, we were still able to detect SpD-specific transcriptional profiles. Previous mouse^23^ and NHP^24^ studies characterized heterogeneity within the NAc along two intersecting axes: a spatial axis separating the lateral core from the medial shell and a molecular axis distinguishing the direct (DRD1 MSNs) from the indirect (DRD2 MSNs) pathways. To interpret the spatial and functional architecture of the transcriptional domains identified in our dataset, we examined how MSN_1-3 SpDs map onto these canonical subdivisions. MSN_1 SpD was enriched (logFC > 1, FDR < 0.05) for *SCN4B, ADORA2A, CALB1, PTPN7*, and *PDE10A* (**Fig. 2B, Table S6**). In the rodent NAc, *CALB1*, is a known marker of core-enriched MSNs^30^ and *ADORA2A* expression varies between the dorso-lateral and ventro-medial core and is enriched in DRD2 MSNs^23^. In contrast, MSN_3 SpD was enriched for *CARTPT, CALCR, PYDC1, PNMA5, NPY2R* and *ARHGAP36* (**Fig. 2C, Table S6**). Specifically, *CARTPT*+ neurons in the medial shell of the mouse NAc play a role in behavior aversion^55^, while *NPY2R* and *CALCR* are enriched in DRD2 MSNs preferentially found in the rodent ventromedial NAc core and shell^23^. MSN_2 was most transcriptionally similar to MSN_1, sharing markers such as *PTPN7, MME*, and *TPBG* (**Table S6**), but also exhibited features intermediate between MSN_1 and MSN_3, such as expression of canonical rodent core and shell marker genes. To determine how these MSN SpDs relate to direct and indirect pathways, we examined expression of canonical pathway marker genes^23^. The direct pathway includes markers such as *DRD1* and *PDYN*, which exhibited moderately higher expression in the medial NAc (MSN_3) (**Fig. 2B,C**). In contrast, indirect pathway markers such as *ADORA2A* and *PENK* exhibited moderately higher expression in the lateral NAc (MSN_1). However, we note that while these differences show some similarities to classical distinctions between direct vs. indirect pathways and core vs. shell identities, they do not support a clear one-to-one mapping between these organizational frameworks and the identified MSN domains.

The non-MSN SpDs were consistent with expected NAc cell types and surrounding anatomical structures. The Inhibitory SpD expressed *GNRH1, SLC5A7, NTRK1, NPY, CORT, SST, KIT, CHODL, NXPH2*, and *GAD1* (**Table S6**), representing a transcriptionally diverse set of inhibitory neurons. *GNRH1* has been reported in a rare class of striatal GABAergic neurons^56^, while *SST* and *CORT* are classic markers of somatostatin-expressing neurons involved in neuromodulatory signaling^46^. Finally, *SLC5A7* is a marker for cholinergic neurons which play a prominent role in modulating the response of MSNs to drugs such as cocaine^41,42^. The Endothelial/Ependymal SpD expressed markers such as *NGFR, CLDN5*, and *SUSD2*, consistent with vascular and choroid plexus cell types. Likewise, the WM SpD expressed canonical oligodendrocyte and myelin-related genes, including *MBP, MOBP*, and *PLP1*. Finally, consistent with prior NAc snRNA-seq studies in human brain^25^, we detected an Excitatory SpD expressing glutamatergic neuronal marker genes, including *SLC17A7, TBR1, NEUROD2*, and *NEUROD6* (**Table S6**). This Excitatory SpD could be detected in adjacent structures outside the boundaries of the NAc, as well as in rare dispersed spots in MSN SpDs. Single molecule fluorescence in situ hybridization (smFISH) for *SLC17A7* verified that no excitatory neuron cell bodies were localized within the NAc (**Fig. S19**). We hypothesized that dispersed *SLC17A7*+ NAc spots could reflect expression in axon terminals from glutamatergic afferents from cortical regions, but we were unable to detect co-localization of *SLC17A7*+ with axonal markers using smFISH/immunofluorescence.

Next, to relate SpDs to transcriptionally-defined MSN subtypes, we performed spatial registration^57,58^ by computing the correlation between enrichment statistics from our SpDs to matched cell types using snRNA-seq data (**Fig. 2D**). DRD1_MSN_A and DRD2_MSN_A subtypes were more strongly associated with MSN_1 SpD. In contrast, D1 islands shared gene expression features with DRD1_MSN_B and DRD1_MSN_D. We extended this analysis to NHP striatal cell types (**Fig. 2E**)^24^, and found that MSN_1 and MSN_2 SpDs were most similar to core-associated subtypes, including D1 Matrix, D2 Matrix, and Striosome MSNs. In contrast, MSN_3 SpD was more similar to shell-biased NHP subtypes, including D1 Shell/olfactory tubercle (OT) and D2 Shell/OT. Human D1 islands corresponded to specialized NHP MSN populations, including D1/D2 hybrid, D1 ICj, and D1 NUDAP (**Fig. 2E, Fig. S20**). A similar organization was observed in rodent data, which confirmed enrichment of *Grm8*-expressing MSNs in our human D1 islands (**Fig. 2F, Fig. S20**).

Finally, to assess how these discrete SpDs contributed to the genetic architecture of human traits, we applied s-LDSC^49^ regression using SpD-specific gene sets (**Table S7**). We found that MSN_1 and MSN_2 SpDs were significantly enriched (FDR < 0.1) for heritability associated with SCZ, bipolar disorder (BPD), and neuroticism. MSN_3 SpD showed enrichment for drinks per week and smoking cessation (**Fig. S21**). Taken together, these findings reveal transcriptionally distinct SpDs in the human NAc that reflect known anatomical features conserved across species with differential contribution to the heritability of neuropsychiatric and addiction-related traits.

### 2.3 Continuous spatial gradients define heterogeneity in MSN spatial domains

Discrete spatial clustering using *PRECAST*^52^ identified three MSN-enriched SpDs (MSN_1–3) that displayed inconsistent boundaries across clustering initializations with small log-fold changes in domain-specific DEGs (**Fig. 2**). These observations suggested that MSN organization in the human NAc may reflect continuous molecular variation rather than discrete compartments. To investigate this further, we applied *MERINGUE*^59^, a method based on topic modeling that identifies latent spatial gene expression patterns based on spatial autocorrelation and local neighborhood structure. We ran MERINGUE^59^ independently on each donor using only SRT spots assigned to MSN SpDs and clustered the resulting spatial patterns using a consensus similarity matrix that incorporated both gene- and domain-level information (**Fig. S22, Fig. S23, Fig. S24**). This approach revealed four reproducible transcriptional gradients, which we refer to as *MERINGUE* consensus patterns (MCPs), shared across donors (**Fig. 3A**).

**Fig. 3.**
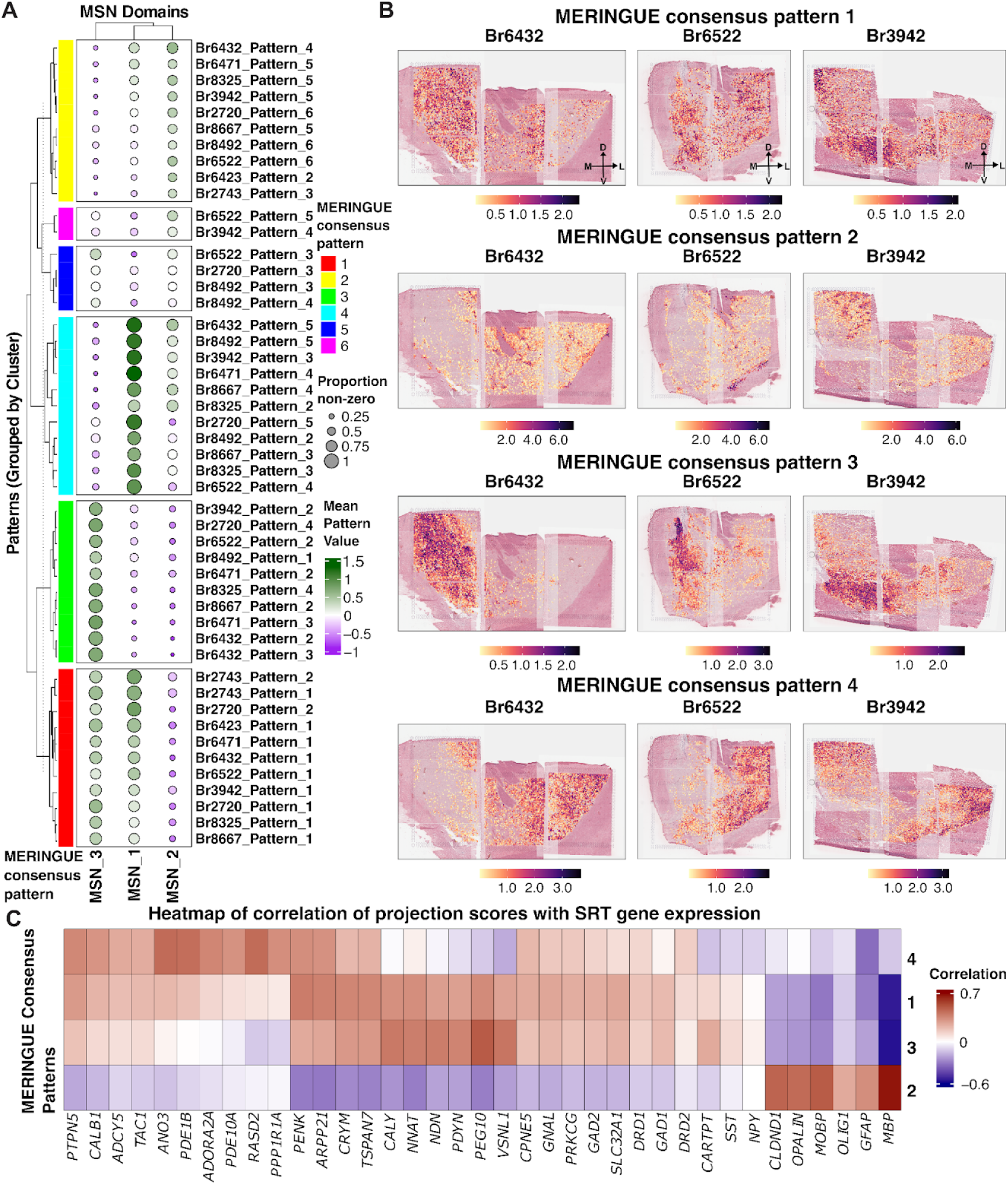
MERINGUE reveals reproducible transcriptional gradients within MSN SpDs. (**A**) Dot plot showing MERINGUE pattern scores across MSN spatial domains (SpDs). MERINGUE patterns inferred independently in each donor are grouped into four MERINGUE consensus patterns (MCPs 1-4) based on a consensus similarity incorporating gene- and domain-level information. Dot size reflects the proportion of non-zero pattern scores within each SpD and color represents the mean pattern score. (**B**) Spatial visualization of MCPs 1–4 across three representative donors (Br6432, Br6522, Br3942). MCP 1 is broadly distributed without directional bias. MCP 2 displays localized enrichment without clear anatomical orientation. MCP 3 localizes to medial regions and overlaps with MSN_3 SpD, while MCP 4 is enriched laterally and aligns with MSN_1 SpD. (**C**) Heatmap showing correlations between MCP scores and expression of selected genes, calculated from SRT data. MCP 2 is enriched for glial-associated genes (e.g., *MBP, MOBP, OLIG1*), MCP 3 is associated with shell-related (*PENK, CRYM, PDYN)* and neurodevelopmental genes (*NNAT, PEG10*), and MCP 4 with canonical DRD2 MSN and core-related markers (e.g., *CALB1, ADORA2A, PDE10A*). MCP 1 shows moderate correlation with similar gene sets, suggesting regions of mixed MSN subtype composition.

MCP 1 showed the highest average pattern scores in MSN_3 SpD, with moderate scores in MSN_1 SpD. MCP 2 transcriptionally aligned most closely with MSN_2 SpD. MCP 3 displayed the highest scores in MSN_3 SpD, while MCP 4 was strongest in MSN_1 SpD. We visualized these patterns at spatial resolution across donors spanning anterior and posterior positions (**Fig. 3B**). MCP 1 was broadly distributed and lacked directional bias, while MCP 2 showed localized enrichment without a clear anatomical gradient. MCP 3 localized to medial regions and overlapped with MSN_3 SpD, while MCP 4 was enriched laterally and aligned with MSN_1 SpD. The spatial positioning of MCPs 3 and 4 was consistent across donors, and reinforced our hypothesis of continuous transcriptional gradients across MSN SpDs.

To characterize the molecular features of each gradient, we examined genes most correlated with MCP scores (**Fig. 3C, Table S8, Methods**). MCP 2 was enriched for glial-associated genes, including *MOBP, MBP*, and *OLIG1*. MCP 3 was linked to shell-enriched neuropeptides in NHP^24^ and rodents^23^ (*CARTPT* and *PDYN), CALY*, which encodes a protein involved in dopamine receptor signaling^60^, and imprinted neurodevelopmental regulators (*PEG10, NNAT, NDN*)^61–63^. MCP 4 was associated with canonical DRD2 MSN and core-related genes, including *ADORA2A, RASD2, PDE1B, ANO3, CALB1, PTPN5*, and *PPP1R1A*, which have been linked to core and matrix compartments across species^23,24^. While MCPs 3 and 4 showed strong associations with distinct MSN SpDs, MCP 1 correlated with many of the same genes at lower magnitude, suggesting it may reflect regions with mixed MSN subtype proportions or transitional spatial environments. Together, these gradients capture continuous transcriptional heterogeneity among MSNs and align with glial, DRD1 MSN, and DRD2 MSN gene signatures.

To further examine the cellular composition underlying these gradients, we performed spot deconvolution using Robust Cell Type Decomposition (*RCTD*)^64^ with cell types identified using our paired snRNA-seq as a reference. Consistent with our earlier spatial registration analysis, the major cell types with non-zero estimated proportions across MSN SpDs included DRD1_MSN_A, DRD1_MSN_C, DRD1_MSN_D, DRD2_MSN_A, DRD2_MSN_B, Astrocyte_A and oligodendrocytes (Oligo) (**Fig. 4A**). DRD1_MSN_A and DRD2_MSN_A showed the highest weights in MSN_1 SpD, while MSN_2 SpD had increased contributions from Oligo. DRD1_MSN_C was most enriched in MSN_3 SpD. We observed similar SpD-level biases when stratifying samples by position along the anterior–posterior (A-P) axis, although the relative proportions of individual cell types varied across donors (**Fig. S25**). Despite these SpD-specific enrichments, many MSN subtypes were represented across all three SpDs, highlighting the spatial intermixing of DRD1 and DRD2 MSNs and supporting the notion that MSN organization reflects continuous, rather than discrete, cellular heterogeneity. To examine directional gradients in MSN cell type distribution, we mathematically projected *RCTD* inferred weights onto SRT spots (**Fig. 4B**). DRD2_MSN_A showed a clear lateral bias, while DRD1_MSN_C was more medial and enriched in MSN_3. To a lesser extent, in samples from the intermediate position, MSN_3 SpD also contained DRD1_MSN_D, a cell type found in adjacent D1 islands (**Fig. S26**).

**Fig. 4.**
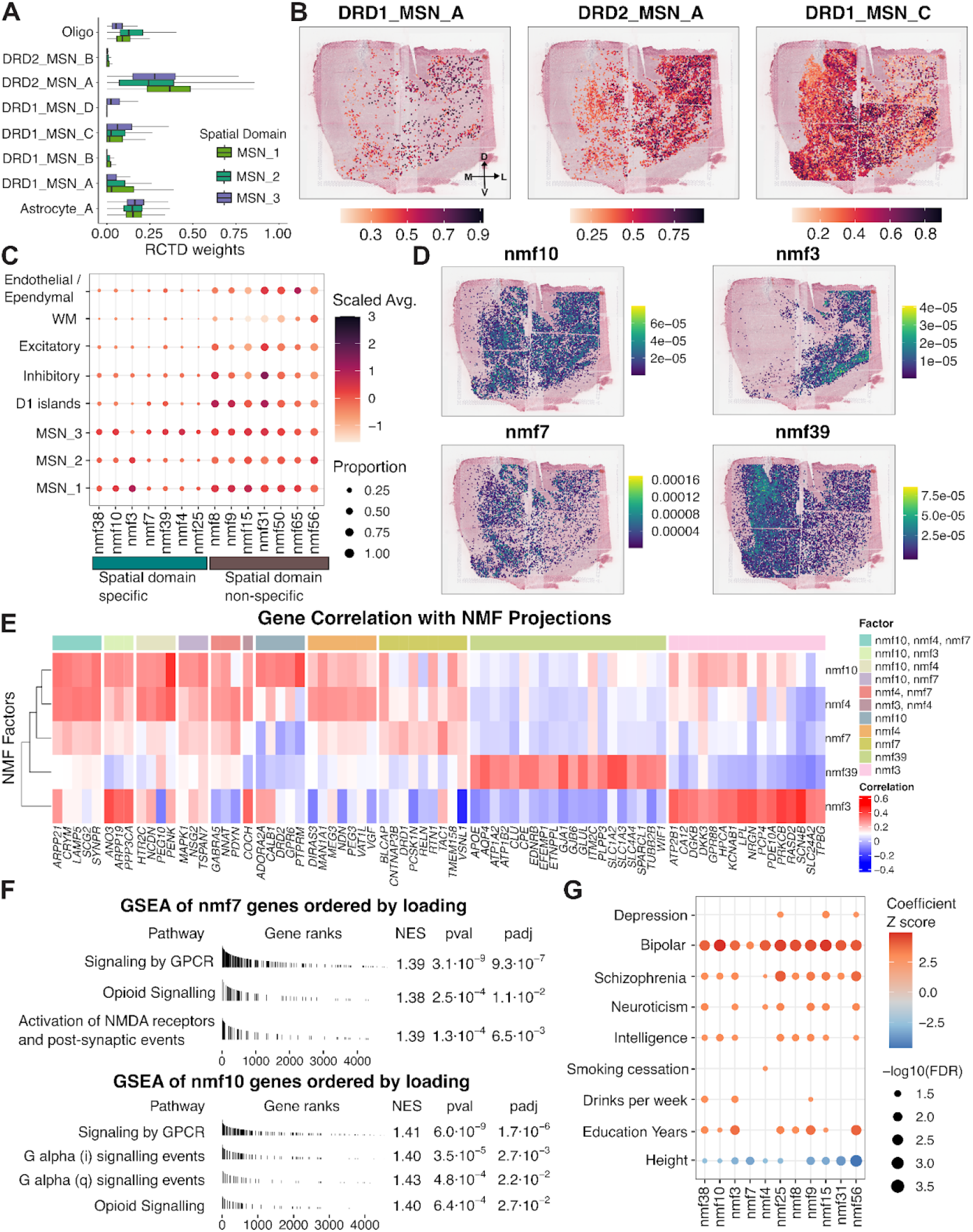
Integrative mapping of MSN subtypes, transcriptional programs, and trait associations in MSN SpDs. (**A**) Robust Cell Type Decomposition (RCTD) using paired snRNAseq-data shows relative contributions of major MSN subtypes, Astrocyte_A and oligodendrocytes within MSN_1-3 SpDs. DRD1_MSN_A and DRD2_MSN_A are enriched in MSN_1 SpD, oligodendrocytes in MSN_2 SpD, and DRD1_MSN_C in MSN_3 SpD. (**B**) Spatial projection of RCTD-derived weights in a representative sample (Br6522) show lateral bias of DRD1_MSN_A and DRD2_MSN_A, and medial bias of DRD1_MSN_C populations. (**C**) Non-negative matrix factorization (NMF) was applied to snRNA-seq data and 56 robust transcriptional factors were retained after removing sex-associated and sparsely represented components. To identify MSN-relevant programs, factors were required to exhibit ≥ 20% nonzero Visium projections and a mean scaled projection > 0.2 within at least one MSN domain. Factors displaying donor-specific biases or non-specific MSN domain representation were excluded, resulting in five MSN_1-3 SpD enriched factors (nmf3, nmf4, nmf7, nmf10, nmf39) used for downstream analyses. (**D**) Spot-level projections of 4 MSN-relevant transcriptional programs (nmf10, nmf3, nmf4, nmf39) for a representative donor (Br6522). Each factor exhibited a spatially distinct pattern. Nmf10 was broadly distributed across all MSN SpDs and nmf3 showed a lateral-ventral bias. nmf7 and nmf39 were medially biased. (**E**) The top genes most strongly associated with each factor were identified by correlating NMF projection weights with spatial gene expression levels, revealing distinct molecular programs. Nmf3 was linked to dopaminergic signaling modulators (*PDE10A, RASD2*), nmf10 to DRD2 MSN identity (*ADORA2A, DRD2*) and core-associated markers (*CALB1*), nmf4 to opioid peptides (*PENK, PDYN*) and serotonin receptors (*HTR2C*), nmf7 to DRD1 MSN identity (*DRD1, TAC1, RELN*), and nmf39 to astrocyte-related genes (*APOE, AQP4, GJA1*). (**F**) Gene set enrichment analysis (GSEA) is shown for nmf7 and nmf10, which were selected as representative DRD1- and DRD2-associated MSN transcriptional programs. nmf7 was enriched for “Activation of NMDA receptors and postsynaptic events” whereas nmf10 was enriched for “GPCR signaling”, “Opioid Signaling”, and “G-protein subunit–specific pathways”. (**G**) s-LDSC analysis of NMF-derived gene sets reveals significant enrichment (FDR < 0.05) of nmf10 and nmf3 for risk variants associated with SCZ and BPD. nmf3 additionally associated with drinks per week, and nmf4 showed significant enrichment for smoking cessation. Gene sets were defined by computing correlations between NMF factor loadings and snRNA-seq gene expression profiles, restricted to protein-coding, non-mitochondrial, non-ribosomal genes. For each factor, genes in the top 10% of correlation values were used as input for s-LDSC. Enrichment results were filtered for traits with known psychiatric, cognitive, or neurological relevance and FDR < 0.05.

To further our analysis of continuous transcriptional gradients in a manner agnostic to pre-defined MSN cell types, we next applied non-negative matrix factorization (NMF) to paired snRNA-seq data and projected the resulting factors onto SRT data. Using a cross-validation framework, we identified 66 as the optimal number of factors for capturing latent transcriptional structure (**Fig. S27**). After excluding two factors associated with donor sex (correlation > 0.3) and eight factors with sparse spatial representation, we focused on the remaining 56 factors (**Fig. S28**). To identify factors enriched in MSN SpDs, we filtered for those that exceeded both a 20% nonzero projection threshold and a mean scaled projection above 0.2 within at least one MSN SpD. This yielded a subset of MSN SpD-specific factors, including nmf38, nmf10, nmf3, nmf7, nmf39, nmf4, and nmf25, for downstream analysis (**Fig. 4C**). We next examined NMF factor projection weights grouped by MSN SpDs across A-P positions (**Fig. S29**). While some factors displayed broad distributions, others showed domain and donor-specific biases. For example, nmf38 and nmf10 were broadly expressed across SpDs, but nmf38 showed particularly strong projection in Br3942, indicating donor-bias. nmf3 was consistently enriched in MSN_1 and MSN_2 SpDs, with lower values in MSN_3 SpD across donors. In contrast, nmf7, nmf39, and nmf4 showed a bias toward MSN_3, though each exhibited weights in other SpDs in a sample-dependent manner. nmf25 did not show a consistent pattern across domains or A-P position. Based on these observations, we excluded nmf38 and nmf25 from downstream analyses and focused on the remaining factors (nmf3, nmf4, nmf7, nmf10, nmf39). We visualized the spatial distribution of these factors using spot-level projection plots across donors in the SRT data. nmf3 showed a consistent lateral bias, aligning with its enrichment in MSN_1 and MSN_2 SpDs. In contrast, nmf10 exhibited no clear directional gradient and was broadly distributed. nmf4, nmf7, and nmf39 were all medially biased, with nmf39 showing the strongest and most consistent medial enrichment across donors. The spatial profiles of nmf4 and nmf7 were more variable, displaying donor-specific patterns along the medial-lateral axis (**Fig. 4D, Fig. S30**).

To interpret the molecular programs underlying each factor, we computed the correlation between each gene’s expression and the factor’s spatial projection scores, and then examined the top 20 correlated genes (**Fig. 4E, Table S9**). nmf3 was characterized by intracellular modulators of dopaminergic signaling, including *PDE10A*^65^ *and RASD2*, a striatal-enriched GTP-binding protein with a putative role in SCZ^66^, alongside synaptic signaling genes such as *PRKCB*^67^ *and NRGN*^68^. These features point to nmf3 as a pattern integrating neuromodulatory input and downstream intracellular signaling to shape MSN excitability. nmf10 was defined by *ADORA2A, CALB1, DRD2*, and *PTPRM* and shared overlap with nmf4 genes, including *PEG10* and *PENK*, suggesting that nmf10 captures DRD2 MSN identity with a core bias despite being broadly distributed. In addition to *PEG10* and *PENK*, nmf4 was also associated with *PDYN* and *NNAT*. While *PDYN*^69^ *and PENK*^70^ *are opioid neuropeptides, NNAT*^63^ *and PEG10*^62^ are imprinted genes implicated in neuronal development and synaptic signaling. These genes suggest that nmf4 may reflect a neuromodulatory program combining opioid peptide signaling with imprinted gene expression. nmf7 was uniquely characterized by DRD1, TAC1, and RELN. TAC1 encodes a neuropeptide released from D1 MSNs, while *DRD1* is a defining feature of the direct pathway MSNs, together indicating a DRD1 MSN-associated factor. Notably, *RELN* was identified as a marker for cocaine activated ensembles in the rodent NAc^71^. Finally, nmf39 was associated with *APOE, AQP4, GJA1*, and *SLC1A2*, consistent with astrocyte-related transcriptional profiles. Together, these factors reveal spatially-defined axes of transcriptional variation in the MSN domains, including differences in dopamine signalling modulation, abundance of direct (DRD1) and indirect (DRD2) pathway neurons, cocaine and opioid-responsive signaling, and glial-MSN interactions.

Gene set enrichment analysis (GSEA)^72^ (**Methods**) revealed that all MSN-enriched factors were associated with “Signaling by GPCR,” with several also enriched for “Opioid Signaling” (nmf3, nmf4, nmf7, nmf10) and G-protein subunit–specific pathways (nmf4, nmf10) (**Fig. 4F, Fig. S31, Table S10**). nmf7 showed additional enrichment for “Activation of NMDA receptors and postsynaptic events,” consistent with its DRD1 MSN identity and the established role of D1 receptor activation in facilitating NMDA receptor signaling in other brain regions^73^. nmf39 was enriched for receptor tyrosine kinase signaling, synaptic transmission, and SLC-mediated transport, suggesting a potential glial-MSN communication axis. s-LDSC^49^ further revealed that nmf10 and nmf3 were associated with genetic risk for BPD and SCZ (**Table S11**). nmf3 was uniquely associated with drinks per week, while nmf4 showed significant enrichment for smoking cessation (**Fig. 4G**). Overall, continuous transcriptional gradients in MSN SpDs are supported by spatially-organized variation in MSN subtype abundance and neuromodulatory and synaptic signaling programs, with each axis of spatial variation showing distinct molecular signatures, spatial distributions, and genetic trait associations.

### 2.4 Cellular composition of D1 islands and inhibitory spatial domains

Given the distinct transcriptional profile of D1 islands, we next sought to delineate the cellular architecture and transcriptional heterogeneity of this SpD. Spatial registration between paired SRT and snRNA-seq data (**Fig. 2D**) identified DRD1_MSN_B and DRD1_MSN_D as the most transcriptionally correlated MSN populations with D1 islands. While this analysis identified an enrichment of DRD1_MSN_B and D cell types in D1 islands, it did not resolve how these cell types were spatially organized or how their contributions varied across individuals. To address these questions, we applied spot deconvolution and NMF to characterize the predicted cellular composition of D1 islands with spatial context.

D1 islands varied along the NAc A-P axis and were most prominent in anterior and intermediate samples, but nearly absent in posterior samples. Based on this observation, we focused our downstream analysis on a subset of four donors in which D1 islands clearly defined (**Fig. 5A**). We applied *RCTD* across all tissue sections and identified cell types enriched in the D1 islands by selecting those with a mean *RCTD* weight greater than 0.06 within this SpD. We identified three DRD1 MSN subtypes (DRD1_MSN_B, DRD1_MSN_C, and DRD1_MSN_D) as well as non-neuronal cell types with measurable contributions (**Fig. 5B**). DRD1_MSN_D showed a modest increase in intermediate samples located posteriorly, while DRD1_MSN_B and DRD1_MSN_C did not exhibit consistent trends (**Fig. 5C**). DRD1_MSN_D was broadly expressed throughout D1 islands, closely matching the domain boundaries, whereas DRD1_MSN_B displayed a more restricted pattern, enriched specifically in medial and ventral regions (**Fig. 5D, Fig. S32**). These spatial trends were consistent across donors, suggesting cellular heterogeneity among individual D1 islands.

**Fig. 5.**
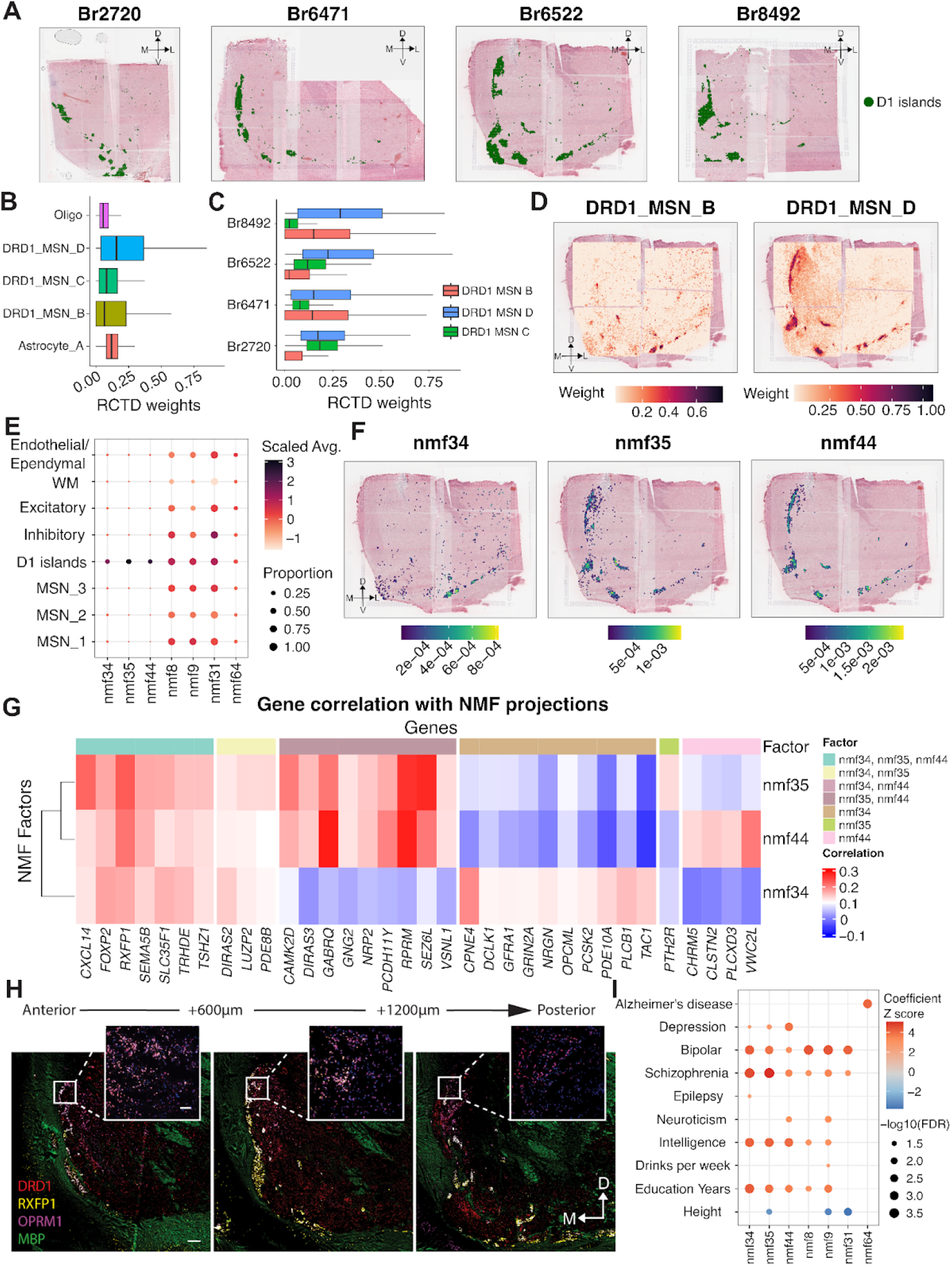
Cellular and transcriptional architecture of D1 islands. (**A**) Anatomical position of the D1 islands (green) across four donors (Br2720, Br6471, Br6522, Br8492), demonstrating localization along the ventro-medial border in intermediate NAc sections. (**B**) RCTD deconvolution weights showing cell types enriched within D1 islands. Cell types were selected based on having a mean *RCTD* weight > 0.06 in the D1 island SpD across donors, which included DRD1 MSN subtypes (DRD1_MSN_B, DRD1_MSN_C, DRD1_MSN_D), astrocytes, and oligodendrocytes. (**C**) Mean RCTD weights for DRD1_MSN_B, DRD1_MSN_C, and DRD1_MSN_D across D1 islands in each donor showing variable subtype contributions across the A-P axis. (**D**) Spatial distribution of DRD1_MSN_B and DRD1_MSN_D across the NAc. DRD1_MSN_D is broadly distributed throughout D1 islands, while DRD1_MSN_B is more restricted to ventromedial regions. (**E**) Dot plot showing NMF factor projection scores grouped by SpD. Only factors with scaled average > 0.2 and proportion > 0.2 in D1 islands are shown. Nmf34, nmf35, and nmf44 had non-zero projection scores uniquely in D1 islands. (**F**) Spatial projections of D1 island-associated NMF factors (nmf34, nmf35, and nmf44) confirm distinct localization patterns across tissue sections with nmf34 showing a more restricted pattern. (**G**) Heatmap of correlation between projection weights and gene expression in SRT data. *CPNE4* and *TAC1* correlate with nmf34; *SEZ6L* and *PTH2R* with nmf35; and *VWC2L, CLSTN2*, and *CHRM5* with nmf44. (**H**) Confocal image showing smFISH for D1 island marker genes (*DRD1, OPRM1, RXFP1)* and WM (*MBP)* across different A-P positions. Inset shows higher magnification of D1 islands. Scale bar 1,000 µm and 100 µm for low magnification and inset, respectively. (**I**) s-LDSC analysis of NMF-derived gene sets reveals significant enrichment (FDR < 0.05) of nmf34, nmf35, and nmf44 for risk variants associated with depression, SCZ, and BPD. nmf44 shows the strongest association with depression compared to SCZ and BPD. Gene sets were defined by computing correlations between NMF factor loadings and snRNA-seq gene expression profiles, restricted to protein-coding, non-mitochondrial, non-ribosomal genes. For each factor, genes in the top 10% of correlation values were used as input for s-LDSC. Enrichment results were filtered for traits with known psychiatric, cognitive, or neurological relevance and FDR < 0.05.

While spot deconvolution provided insight into the cellular composition of D1 islands, it was limited by reliance on predefined snRNA-seq labels. To identify continuous and data-driven transcriptional variation within D1 islands, we performed NMF on snRNA-seq data and identified three factors representing D1 islands, including nmf34, nmf35, and nmf44 (**Fig. 5E**). When we spatially visualized these factors, we found that nmf35 and nmf44 were broadly expressed across D1 islands, while nmf34 was spatially restricted to medial and ventral D1 islands (**Fig. 5F, Fig. S33**). These patterns closely mirrored those observed with spot deconvolution, reinforcing the presence of fine-grained molecular organization within these D1 interface islands. Next, we examined the gene composition of nmf34, nmf35, and nmf44, and identified a shared set of genes consistently associated with all three factors, including *FOXP2, NRG1, OPCML*, and *SLC35F1* (**Fig. S34**). These genes are broadly implicated in neurodevelopmental processes^74–76^ and linked to genetic risk for psychiatric disorders, such as SCZ^77^. Notably, *RBFOX1*, a neuronal splicing regulator frequently disrupted in neurodevelopmental syndrome^78^ was among the top 50 genes by loading for each program, reinforcing its shared relevance within D1 island MSNs. Beyond this shared set of D1 island genes, each factor displayed distinct transcriptional features suggesting heterogeneity among individual D1 islands. nmf34 was strongly correlated with *CPNE4* and *TAC1*, genes marking DRD1_MSN_B. nmf35 also showed moderate correlation with *CPNE4*, but was more enriched for *CAMK2D, SEZ6L*, and *PDE8B*, which are involved in synaptic plasticity^79^ and trafficking of kainate receptors^80^. nmf44, by contrast, captured a separate axis defined by high correlation with *VWC2L, CLSTN2*, and *EYA2*, pointing to roles in cell adhesion and plasticity^81–83^ within D1 islands.

To validate and extend these observations in a spatial context, we correlated SRT gene expression with NMF projection scores, assessing whether the gene co-expression patterns underlying latent factors in snRNA-seq are preserved in SRT profiles. This analysis recapitulated key trends observed in the snRNA-seq data (**Fig. 5G, Table S12**). *FOXP2, RXFP1*, and *SLC35F1* remained consistently correlated with all three factors, reinforcing their role as key components of D1 island transcriptional identity. *CPNE4* and *TAC1* again emerged as specific to nmf34, while *SEZ6L* and *PTH2R* were preferentially associated with nmf35. nmf44 was distinguished by association with *VWC2L, CLSTN2*, and *CHRM5*. The factor-specific association of *CPNE4* and *RXFP1* reflects a transcriptional division within D1 islands consistent with findings from the NHP striatum^24^. Analysis of broader gene set overlap revealed that nmf44 and nmf34 contributed the largest number of unique genes, while 62 genes were shared across all three factors (**Fig. S35**). These findings reinforce that while the D1 islands share a common transcriptional architecture, they also display heterogeneity characterized by partially overlapping, but distinct gene programs. Finally, we orthogonally validated the presence of spatially restricted D1 islands in NAc using smFISH for *DRD1, RXFP1*, and *OPRM1* (**Fig. 5H, Fig. S36**).

To further interpret the functional relevance of NMF factors enriched in D1 islands, we performed GSEA and s-LDSC. All three factors showed significant enrichment for the Reactome pathway^84,85^ “Activation of NMDA receptors and postsynaptic events”, which plays a role in morphological changes in MSNs in response to cocaine^86,87^. nmf34 was additionally enriched for “DAG and IP3 signaling” and “Activation of kainate upon glutamate binding”, pointing to second messenger cascades associated with synaptic activity in ventromedial D1 islands (FDR < 0.05; Fig. S37, Table S13). nmf35 showed stronger associations with “Axon guidance” and AMPA receptor signaling pathways, consistent with its broader spatial distribution and link to synaptic remodeling. nmf44, in contrast, was enriched for processes related to membrane organization and synaptic adhesion, including the “Dopamine neurotransmitter release cycle” and “Interaction between L1 and ankyrins” (FDR < 0.05), suggesting that it captures a distinct signaling axis within the D1 islands. We next asked whether these latent factors were enriched for genetic risk associated with neuropsychiatric traits. Using s-LDSC^49^, we found that nmf44 showed the strongest enrichment for depression risk loci (**Fig. 5I**), while nmf35 showed a weaker, but still nominally significant signal. All three factors, nmf34, nmf35, and nmf44 were enriched for SCZ and BPD risk loci (**Table S11**). These findings suggest that distinct transcriptional modes within D1 islands may differentially relate to the genetic architecture of neuropsychiatric traits.

Having characterized the cellular composition and transcriptional organization of MSN_1-3 SpDs and the D1 islands, we next examined how non-MSN populations are spatially positioned in relation to these SpDs. *RCTD*-based mapping of six GABAergic populations (Inh_A-F) revealed distinct spatial patterns (**Fig. S38**). Inh_E, which expressed *NPY, CORT*, and *SST*, formed dense pockets within MSN SpDs. This distribution is consistent with previous descriptions of *NPY*-expressing neurons modulating MSN excitability^88^. Inh_A showed a similar pattern, but was more diffusely distributed. Inh_C, enriched in *GLP1R* and *TAC3*, localized specifically to medial and dorsal regions of the NAc, overlapping with MSN_3 SpD and in close proximity to D1 islands (**Fig. S39**). In contrast, Inh_B, Inh_D, and Inh_F were peppered throughout the NAc, but also densely localized to neighboring regions adjacent to the NAc (**Fig. S40**). Notably, *CHAT*-expressing cholinergic neurons (Inh_D), which modulate MSN activity and dopamine release^89^, showed higher prevalence in regions neighboring D1 islands. Taken together, these findings suggest that D1 islands represent a specialized, transcriptionally heterogenous SpD with unique neighboring non-MSN populations that may impact addiction and neuropsychiatric traits.

### 2.5 Spatial mapping of Ligand-Receptor (LR) interactions associated with neuropsychiatric traits

Building on our s-LDSC analyses, which linked specific NAc cell types and SpDs to the heritability of neuropsychiatric traits, we next sought to identify molecular pathways that might underlie these genetic associations. We applied *LIANA*^*90*^ to our snRNA-seq data to infer candidate Ligand-Receptor (LR) interactions across cell types. We intersected these predictions with the OmniPath^91^ intercellular interaction network and filtered to LR interactions where at least one partner was a risk gene for depression, anxiety, substance dependence, or SCZ as defined by OpenTargets^92^. Across all traits, we identified both shared and disorder-specific LR pairs, revealing overlapping involvement of neuromodulatory, neuropeptide, and axon guidance signaling (**Fig. 6A, Fig. S41, Table S14**). Several interactions were shared across multiple disorders including neurotrophic signaling (*BDNF*-*NTRK2*), which supports synaptic plasticity and contributes to the stress response^93^ and drug-seeking^94^. Other shared interactions include tachykinin signaling (*TAC1*-*TACR3/DPP4*), which influences aversive learning^95^, and stress hormone signaling (*CRH-CRHR1*) which mediates dendritic atrophy in the accumbens of mice^96^ and stress response^97^. In contrast, trait-specific LR pairs revealed unique signaling interactions. Anxiety-associated interactions were enriched for cell adhesion (*JAM3-ITGB1*) and Wnt-signaling modulators (*DKK2-LRP5*). Depression-associated pairs spanned Notch (*JAG1-NOTCH2, DLL1-NOTCH3, DLL4-NOTCH3*), ephrin (*EFNA5-EPHA5, EFNA5-EPHA7, EFNA5-EPHA4*), and extracellular matrix pathways (*SLIT1-ROBO1, NTN1-DCC*). SCZ-specific pairs highlighted growth factor signaling (*TGFB2-TGFBR3*), and immune-related pathways (*IL18-IL1RAPL1*). Substance dependence-specific pairs included opioid and neuropeptide systems (*PENK-OPRM1, PDYN-OPRM1*), alongside chemokine (*CXCL12-DPP4*) and axon guidance signaling (*RGMA-NEO1*).

**Fig. 6.**
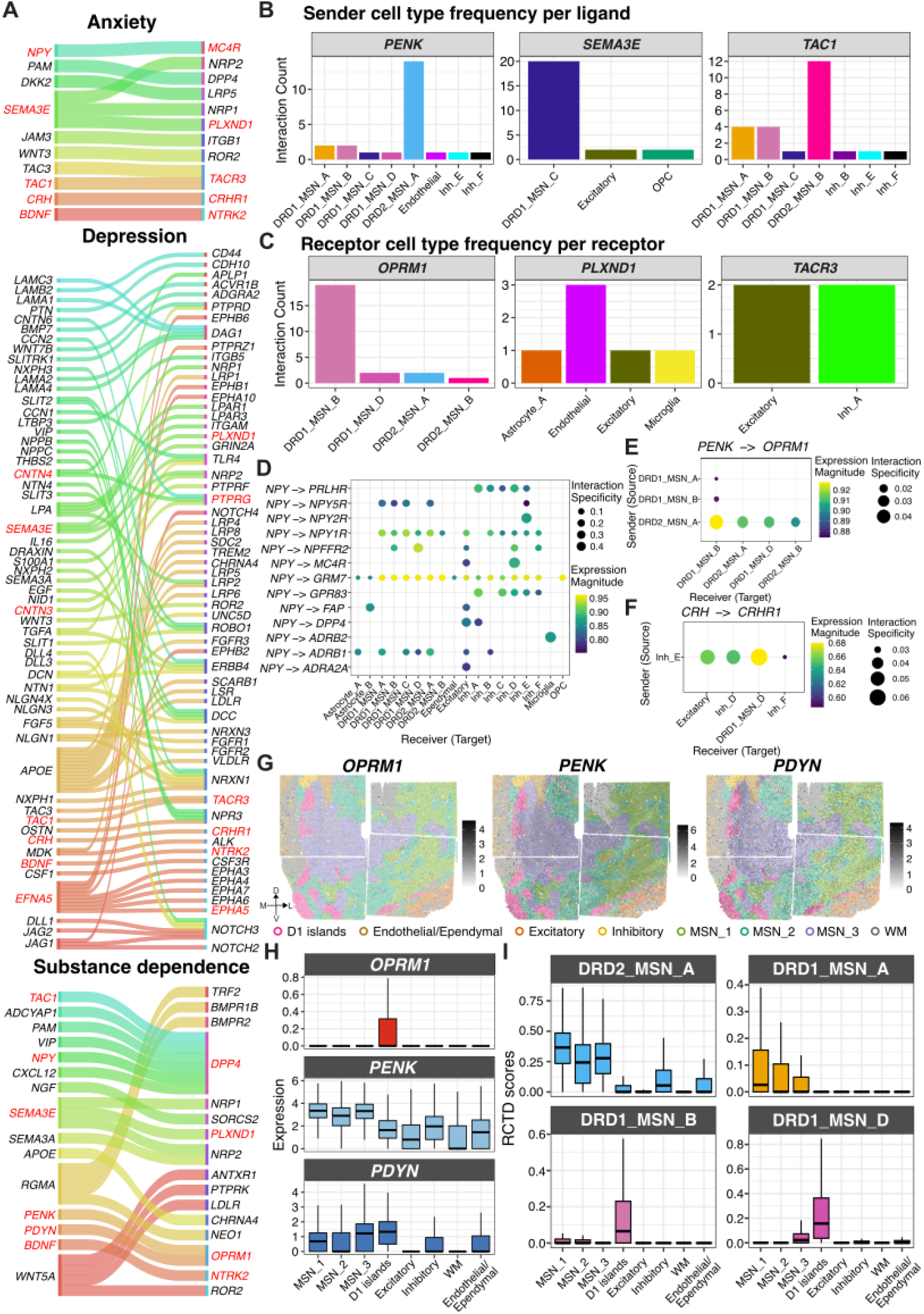
Trait-associated ligand–receptor (LR) interactions reveal molecular pathways underlying neuropsychiatric risk in the human NAc. (**A**) Sankey plots summarizing predicted LR interactions involving risk-associated genes for anxiety, depression, and substance dependence. *LIANA*-inferred LR pairs were filtered to include only those where at least one partner overlapped with risk loci from OpenTargets, revealing both shared and trait-specific signaling pathways. Notable shared interactions include *BDNF*-*NTRK2, TAC1*–*TACR3*/*DPP4*, and *CRH*–*CRHR1*. Trait-specific pathways include Notch, ephrin, and extracellular matrix-related signaling in depression, Wnt and adhesion-related signaling in anxiety, and opioid, chemokine, and axon guidance pathways in substance dependence. (**B**) Frequency of sender cell types for 3 selected ligands which exhibit spatially heterogeneous gene expression and are relevant across multiple traits (*PENK, SEMA3E*, and *TAC1*). *PENK* ligands originated predominantly from DRD2_MSN_A, *SEMA3E* from DRD1_MSN_C, and *TAC1* from DRD2_MSN_B, DRD1 MSNs and selected inhibitory populations. (**C**) Frequency of receiver cell types for selected receptors (*OPRM1, PLXND1, TACR3*). *OPRM1*-mediated signaling was enriched in DRD1_MSN_B populations localized to D1 islands, while *PLXND1* receptors were expressed in endothelial and non-neuronal cell types, and *TACR3* in excitatory and Inh_A neurons. (**D**) Predicted *NPY*-associated LR interactions reveal diverse target specificity. While broad interactions such as *NPY*-*GRM7* span MSN, excitatory, and inhibitory populations, more selective signaling was observed for *NPY*-*MC4R* (excitatory and Inh_D) and *NPY*–*DPP4* (excitatory and Inh_A). (**E**) Among *PENK*-*OPRM1* interactions inferred by *LIANA, PENK* ligands originated almost exclusively from DRD2_MSN_A and targeted *OPRM1* in DRD1_MSN_B populations localized to D1 islands. (**F**) CRH–CRHR1 signaling was mediated primarily by Inh_E neurons, with receptor expression enriched in DRD1_MSN_D cells within D1 islands, as well as excitatory neurons and Inh_D cholinergic populations. (**G**) Spatial maps of *OPRM1, PENK*, and *PDYN* expression in a representative donor (Br6522) reveal enrichment of *OPRM1* in D1 islands, widespread *PENK* expression across MSN_1–3 SpDs, and elevated *PDYN* in medial MSN_3 SpD. (**H**) Domain-level quantification of LR expression shows *OPRM1* is highly enriched in D1 islands, while *PENK* is broadly expressed across MSN SpDs and *PDYN* peaks in D1 islands and medial MSN_3 SpD. (**I**) RCTD-inferred cell type weights confirm that *PENK*-*OPRM1* interactions originate primarily from DRD2_MSN_A and target *OPRM1*-expressing DRD1_MSN_B/D populations in D1 islands, whereas *PDYN* senders (DRD1_MSN_A) are distributed outside medial MSN_3 SpD, suggesting that *PDYN* expression in this medial region arises from a subset of DRD1_MSN_A cells proximal to D1 islands.

Next, we examined the cell types that contributed most strongly to LR interactions shared across traits with a focus on *PENK, SEMA3E*, and *TAC1*, three ligands that were also key molecular features in our SRT analysis (**Fig. 6B**). Filtering for high-confidence interactions, we found *PENK* ligands originated almost exclusively from DRD2_MSN_A, identifying indirect pathway MSNs as the predominant sender population for enkephalinergic signaling. *SEMA3E* ligands originated from DRD1_MSN_C, and *TAC1* senders were DRD1 MSN subtypes, with additional contributions from inhibitory populations. On the receptor side (**Fig. 6C**), *LIANA* predicted that *OPRM1*-mediated interactions occurred in the DRD1_MSN_B cell population, which localized to D1 islands (**Fig. 5B,D**). *PLXND1*-mediated interactions occurred in endothelial and non-neuronal populations, while *TACR3*-mediated interactions occurred in excitatory neurons and Inh_A neurons.

*NPY* interactions (*NPY-MC4R* and *NPY-DPP4*) also emerged as risk-associated LR pairs across multiple traits (**Fig. 6D**). While broad interactions such as *NPY-GRM7* were detected across MSN and inhibitory cell types, we also observed receptor- and cell type-specific signaling. For example, *NPY-ADRB1* and *NPY-NPY5R* involved different, yet overlapping MSN populations suggesting that *NPY* engages the β1-adrenergic receptor and Y5 receptor differently across MSN subpopulations. By contrast, *NPY-MC4R* interactions were more restricted, targeting excitatory neurons and Inh_D (cholinergic) populations, while *NPY-DPP4* pairs involved excitatory neurons and Inh_A populations.

Next, we sought to delineate the specific sender-receiver pairs underlying *PENK*-*OPRM1* and *PDYN*-*OPRM1* signaling. Among the identified interacting pairs, *PENK* ligands predominantly originated from DRD2_MSN_A and targeted *OPRM1* in DRD1_MSN_B populations located in D1 islands (**Fig. 6E**). In contrast, *PDYN*-*OPRM1* signaling involved a single interacting pair, with DRD1_MSN_A predicted as the exclusive sender population (sca.LR score∼0.87, natmi.edge specificity∼0.05). Additionally, we found evidence that D1 islands may also play a role in stress-related signaling through *CRH*-*CRHR1* interactions (**Fig. 6F**). Here, Inh_E interneurons emerged as the predominant *CRH* senders, while receptor expression was enriched in DRD1_MSN_D (localized to D1 islands), excitatory neurons and Inh_D (cholinergic) populations, which is consistent with previous work showing that CRH influences the physiology and functional diversity of cholinergic neurons^98,99^. Collectively, these findings suggest that D1 islands integrate both opioid-related and stress hormone signaling, supporting a role in reward and affective processing.

To spatially map LR interactions, we integrated our snRNA-seq findings with SRT data. We characterized both contact-dependent interactions and paracrine signaling. For contact-dependent signaling, we focused on membrane-bound contactin ligands (*CNTN3, CNTN4, CNTN6*) that interact with *PTPRG*, and play a pivotal role in axon guidance and synapse formation^100^. *LIANA*-inferred interactions revealed distinct sender populations, but largely convergent receiver cell types (**Fig. S42**). Using SRT, we identified spots co-expressing ligands and receptors, and quantified their distribution across spatial domains. Spots co-expressing either *CNTN3-PTPRG, CNTN4-PTPRG*, or *CNTN*6-*PTPRG* were differentially distributed across SpDs with a higher proportion of *CNTN4*-*PTPRG* co-expressing spots in the D1 islands, MSN_1, and Inhibitory SpDs. Because coexpressing spots often contained multiple cell types, we used *RCTD*-derived weights to identify contributing populations and assess whether *LIANA*-predicted interactions were supported by inferred cell types (**Fig. S42**). The two approaches were largely in agreement. For example, for *CNTN4*-*PTPRG*, the highest co-occurrence was observed among DRD2_MSN_B, Inh_A, and Inh_F populations, consistent with the predicted involvement of DRD2_MSN_B as a primary sender (**Fig. S42**). Additionally, we identified co-localization of cell types, such as DRD1_MSN_A with Inh_A, and DRD1_MSN_D with Inh_F, which were distinct from *LIANA* predicted interactions for *CNTN6*-*PTPRG*. Thus, integrating SRT with *LIANA*-inferred interactions allowed us to refine cell type relationships and add spatial context essential for interpreting contact-mediated signaling.

Given the well-established role of opioid signaling in the NAc and the association of *OPRM1* with addiction, we focused on spatially mapping *PENK*-*OPRM1* and *PDYN*-*OPRM1* paracrine signaling (**Fig. 6G**). By quantifying LR expression across SpDs (**Fig. 6H**), we found that *OPRM1* was enriched within D1 islands with minimal expression in other domains. *PENK* was broadly distributed with relatively consistent levels across MSN_1-3 SpDs and lower expression in inhibitory and non-neuronal regions. *PDYN* was slightly higher in D1 islands and MSN_3 SpD, consistent with its partial overlap with *OPRM1*-expressing regions. *RCTD* deconvolution weights corresponding to DRD2_MSN_A (the principal *PENK* sender as predicted by *LIANA*) were elevated across all three MSN SpDs, with modestly higher weights in MSN_1 SpD, mirroring the expression patterns of *PENK*. Both *PENK* and *PDYN* signaling converge on *OPRM1*-expressing DRD1_MSN_B populations, which exhibited the highest deconvolution weights in the D1 islands as expected (**Fig. 6I**). *PENK* also targets *OPRM1*-expressing DRD1_MSN_D populations, which likewise localize to D1 islands (**Fig. 6I**). In contrast, while *LIANA* inferred DRD1_MSN_A as the main sender for *PDYN*, the highest weights for DRD1_MSN_A were observed in MSN_1 and MSN_2 SpDs, i.e. outside the medial MSN_3 SpD (**Fig. 6I**) where *PDYN* expression peaks (**Fig. 6H**). This suggests that while DRD1_MSN_A cells are less prevalent in MSN_3 SpD, it is possible that they express more *PDYN* in this SpD. Our findings highlight the significance of integrating SRT, which provides spatial expression patterns that cannot be resolved solely by snRNA-seq-based inference of interacting cell types. In conclusion, these findings reveal that trait-associated signaling in the human NAc emerges from coordinated interactions across diverse cell populations and demonstrate how integrating SRT and snRNA-seq data improves our understanding of LR interactions involved in neuropsychiatric traits and addiction.

### 2.6 Spatial correlates of response to acute and volitional morphine

We next aimed to demonstrate the utility of our integrated SRT and snRNA-seq data for linking transcriptional signatures from external datasets to spatially organized molecular circuits in the human NAc, providing a framework that can be broadly applied across diverse experimental contexts. Following the characterization of neuropeptide and μ-opioid receptor signaling in NAc cell types and SpDs, we next examined which SpDs exhibit the strongest transcriptional responses to morphine. To do so, we leveraged snRNA-seq data generated by Reiner et. al^27^ from the rat NAc, which profiled transcriptional changes across three paradigms: acute morphine injection, volitional morphine self-administration, and yoked saline (control). Notably, this dataset captured morphine induced transcriptional changes across DRD1 and DRD2 MSNs, interneurons, astrocytes, and oligodendrocytes. For each paradigm, we split DEGs into upregulated and downregulated genes and tested the over-representation of these rat gene sets with human SpD marker genes (**Methods**). For acute morphine, oligodendrocyte DEGs were significantly enriched in WM, with upregulated genes showing strong over-representation. Upregulated D1R MSN-2 genes were enriched across MSN_1-3 SpDs with the strongest enrichment in MSN_3 SpD. In contrast, downregulated D2R MSN-3 genes were enriched in MSN_1 and MSN_2 SpDs (**Fig. S43A**). For volitional morphine self-administration, upregulated DEGs showed stronger enrichment within MSN_1 SpD, particularly for D1R MSN-2, D1R MSN-3, and MSN-2. We also identified significant enrichment of upregulated D2R MSN-2, D2R MSN-4, and MSN-2 DEGs within D1 islands (**Fig. 7A**).

**Fig. 7.**
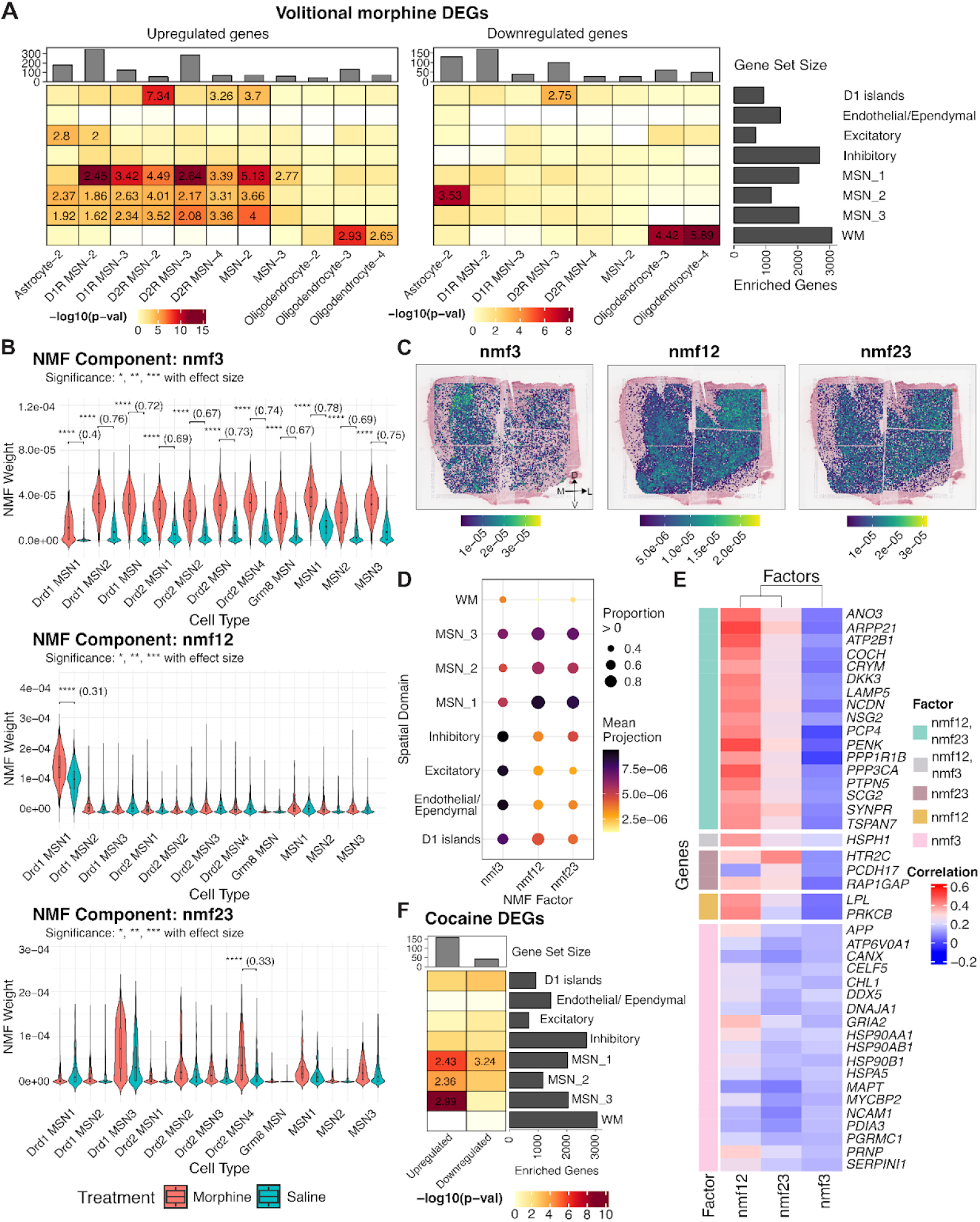
Spatial mapping of morphine- and cocaine-associated transcriptional programs in the human NAc. (**A**) Enrichment of DEGs associated with volitional morphine intake from rat NAc snRNA-seq ^27^ across human NAc SpDs. Upregulated DEGs identified in multiple rat cell types, including D1R MSN-2, D1R MSN-3, D2R MSN-3, and MSN-2 were enriched in human MSN_1 SpD. In contrast, upregulated rat genes identified in D2R MSN-2 showed the strongest enrichment in the human D1 islands, with additional, more moderate enrichment observed across MSN_2 and MSN_3 SpDs. Downregulated DEGs detected in Astrocyte-2 were enriched in MSN_2. (**B**) Violin plots showing factor weights for 3 morphine-associated factors inferred from MSN-restricted rodent snRNA-seq data^27^ (nmf3, nmf12, and nmf23), grouped by cell type. Each plot compares morphine- and saline-exposed cells, with all three factors exhibiting significantly higher weights in cells from the volitional morphine condition. Nmf3 shows broadly distributed weights across multiple MSN subtypes, whereas nmf12 and nmf23 display more cell type-specific weighting patterns. (**C**) Spatial projections of morphine-associated factors onto SRT data (Br6522) revealed distinct anatomical patterns. (**D**) Quantification of factor projection weights across SpDs revealed distinct patterns of anatomical bias. Nmf12 and nmf23 were broadly distributed across MSN_1–3 SpDs. Nmf3 exhibited a more widespread pattern, with moderate weights across multiple MSN domains and elevated contributions in D1 islands, inhibitory, excitatory, and endothelial/ependymal SpDs. (**E**) Heatmap showing the top genes most positively correlated with each factor’s spatial projection scores, calculated based on correlations between NMF projection weights and spatial gene expression. Nmf12 was associated with dopaminergic and plasticity-related genes, including *PENK, ARPP21, PTPN5*, and *PRKCB*. Nmf23 captured a mixed program combining dopamine-related genes, such as *ARPP21, PENK*, and *PTPN5*, with a serotonergic component marked by *HTR2C*. Nmf3 reflected a generalized stress and metabolic response program, characterized by genes such as *HSPH1, HSP90AA1*, and *NCAM1*. (**F**) Enrichment of cocaine-associated DEGs derived from rat snRNA-seq ^26^ mapped onto human SpDs. Upregulated DEGs from rat Drd1 MSN 1 were enriched in human MSN_1-3 SpDs.

While DEG enrichment analysis identifies morphine-induced transcriptional changes within individual cell types and SpDs, it does not capture transcriptional patterns that are shared across MSN populations. To investigate the presence of these patterns, we applied NMF to MSN-restricted expression data and identified separate morphine-associated factors for both acute and volitional morphine exposure.

For acute morphine, we identified two morphine-associated factors: nmf12, which was most weighted in DRD1 MSN-1, and nmf3, which exhibited loadings broadly across multiple MSN cell types (**Fig. S43**), and projected these factors into the SRT data. nmf12 showed broad representation across all three MSN SpDs, whereas nmf3 exhibited a subtle medial bias and extended into WM (**Fig. S43**). Rodent genes with the highest loadings in nmf12 included (**Table S15**) *Ntrk2* encoding TrkB, the high affinity receptor for *BDNF*, which promotes growth and survival of MSNs^101,102^; *Pde7b*, which inhibits dopaminergic death^103^; and *Sik2*, which regulates neuronal survival under stress^104^. Genes whose expression was most strongly correlated with nmf12 projections in the SRT data (**Table S16**) included *PENK* and other genes that modulate dopamine signaling (e.g., *PRKCB* and *PTPN5*)^105,106^. Thus nmf12 is a morphine-associated factor which captures neuronal survival and dopaminergic signaling while nmf3 captures a generalized morphine-associated factor which reflects stress response, protein folding, and metabolic regulation indicated by the high loadings of rodent genes *Zmym4, Eor1b*, and *Hsp90aa1*^107,108^.

For chronic morphine self-administration, we identified three morphine-associated factors (nmf3, nmf12, and nmf23) with varying contributions to different MSN cell types (**Fig. 7B**). When projected into the SRT data, each factor displayed distinct anatomical patterns (**Fig. 7C**). While the cell type-specific factors nmf12 and nmf23 exhibited the highest weights in MSN SpDs, especially MSN_1, the non cell type-specific factor, nmf3, displayed highest weights in inhibitory, excitatory, D1 islands, and endothelial/ependymal SpDs (**Fig. 7D**). Finally, we interpreted these factors based on the rodent genes with the highest factor loadings (**Fig. S44, Table S15**), and human genes whose spatial expression was most strongly correlated with factor projections (**Fig. 7E, Table S16**). Consistent with the acute morphine analysis, nmf12 captured a dopamine- and plasticity-associated program, while nmf3 reflected a generalized stress and metabolic response based on overlap with top loaded rodent genes. In contrast, nmf23 highlighted a distinct morphine-associated signature characterized by genes including Pcdh9, Tenm3, Htr2c, and *Opcml. Tenm3* plays a role in topographical patterning of thalamostriatal projections^109^ and *Htr2c* encodes the 5-HT2C serotonin receptor, activation of which has been shown to selectively inhibit morphine-induced dopamine efflux in the NAc^110^. Human spatial expression correlations for nmf23 overlapped partially with nmf12, including *ANO3, COCH, CRYM, PENK*, and *PTPN5*, but also showed exclusive associations with *HTR2C* and *PCDH17*. Together, these findings suggest that while nmf23 shares dopaminergic and synaptic plasticity signatures with nmf12, it also captures a distinct serotonergic component.

Finally, we performed a similar analysis with cocaine-specific transcriptional signatures in rat snRNA-seq data^26^. Among all cell types, only Drd1 MSN 1 exhibited more than 25 DEGs (FDR < 0.05) when comparing cocaine-vs. saline-treated animals. This rodent cell type was most transcriptionally similar to our human DRD1_MSN_A population (**Fig. S10**) and human MSN_1 SpD (**Fig. 2F**). Gene set enrichment analysis showed that upregulated cocaine-associated DEGs were significantly enriched across MSN_1, MSN_2 and MSN_3 SpDs (**Fig. 7F**). Notably, upregulated DEGs shared by MSN_1 and MSN_3 SpDs included *PDE7B, ARPP21*, and *GRIN1*, highlighting their collective involvement in dopaminergic and glutamatergic signaling pathways that regulate synaptic plasticity and reward-related behaviors. However, upregulated genes also included MSN_1 SpD-specific genes, such as *TAC1*, which forms local circuits that influence stress-induced anhedonia behavior^111^ and avoidance response^112^. To extend this analysis, we also examined NMF patterns (k = 30) using acute cocaine snRNA-seq data, and identified two factors, nmf15 and nmf28, which were significantly more loaded in cocaine cells compared to saline cells (**Fig. S46, Table S15, Table S16**). nmf15 captured a synaptic plasticity and dopaminergic signaling program, with similarities to morphine-associated nmf12, and its spatial projections were distributed across MSN SpDs. In summary, these analyses reveal morphine- and cocaine-associated transcriptional programs that span MSN cell types and SpDs, demonstrating the capacity of these circuits to coordinate drug-responsive pathways while also uncovering specialized roles for D1 islands and *TAC1*+ neurons. More broadly, this work establishes a roadmap for integrating external functional datasets with human spatiomolecular data to dissect conserved and drug-specific molecular adaptations within the NAc.

## 3 Discussion

Here we generated the first comprehensive spatiomolecular atlas of the human NAc by integrating snRNA-seq with SRT of postmortem NAc tissue across ten neurotypical adult donors. We identified 20 transcriptionally distinct cell types, including multiple DRD1 and DRD2 MSNs, several classes of non-MSN neurons, and non-neuronal populations. Analysis of SRT data revealed continuous gradients of MSNs across the mediolateral axis and transcriptionally distinct D1 islands on the ventromedial NAc border. Cross-species comparisons demonstrated conservation of both MSN cell types and spatial features, including D1 islands corresponding to *RXFP1+* and *CPNE4+* islands in the rodent^26,35^ and NHP^24^. Although we did not observe discrete core and shell compartments, MSN gradients displayed transcriptional profiles that were core-like or shell-like, indicating strong alignment with known organizational features described in the NHP. Leveraging this spatiomolecular atlas, we linked specific spatial domains (SpDs) to neuropsychiatric traits and identified trait-relevant ligand-receptor interactions. Notably, SRT data enabled the localization of candidate interacting cell types and improved our interpretation of contact-dependent signaling. Finally, we projected transcriptional programs associated with morphine and cocaine exposure from rodent NAc datasets onto the human SRT data revealing spatial patterns of drug-responsive gene expression across MSN populations.

In addition to defining the spatial architecture of the human NAc, we found evidence that its transcriptionally defined SpDs vary along the A-P axis. D1 islands were prominent in anterior and intermediate sections, but largely absent in posterior regions. This is consistent with studies showing the distribution of transcriptionally defined MSN subtypes varies systematically along the A-P axis in the mouse striatum^23^. Beyond these discrete SpD differences, our SRT analyses revealed spatial organization that was not entirely captured by the classic core vs. shell framework. *MERINGUE* and NMF-based analyses uncovered continuous gradients of MSN identity spanning the medial-lateral axis, demonstrating intermixing of DRD1- and DRD2-expressing MSN populations. Continuous gradients of MSNs and spatial gradients across the NAc have been previously described in the rodent NAc^19,23,29^. Chen et al.^23^ showed that sharp core vs. shell boundaries are most prominent in posterior regions of the mouse NAc, whereas the anterior NAc exhibits more graded transitions between MSN subtypes^23^. Consistent with this, the continuous gradients we observed in the human NAc likely reflect the anatomical positions represented in our dataset. Together, these findings suggest that MSN diversity in the human NAc is organized along continuous spatial axes, with gradients of gene expression that vary across both medial-lateral, dorsoventral, and potentially the A-P axis. Given the importance of A-P position, future studies should systematically sample the human NAc along the full extent of the A-P axis within the same donors to delineate donor-specific effects from A-P position. It will also be crucial to explore transcriptional changes in the NAc over brain development to understand how MSN heterogeneity arises during development, and its stability across the lifespan. Finally, the continuous gradients identified by *MERINGUE* and NMF analysis highlight the need for higher resolution imaging-based SRT data to precisely characterize the organization of co-occurring MSN subtypes. The datasets generated here will provide a critical foundation for designing custom gene panels for follow up SRT studies at cellular resolution.

The spatial organization of the human NAc has functional implications, which are evident within the specialized D1 islands. These transcriptionally distinct compartments are enriched for μ-opioid receptor (*OPRM1*) expression and receive inputs from *PENK* and *PDYN* expressing neurons, positioning them as central hubs for integrating opioid signaling. Their medial bias aligns with rodent studies identifying a rostrodorsal hotspot in the medial shell that mediates opioid-induced hedonic enhancement^36^. D1 islands map to specific MSN cell types, including DRD1_MSN_B and DRD1_MSN_D, which share transcriptional features with rodent Grm8 neurons characterized by *Chst9* and *Tshz1* expression^35^. Consistent with this, Andraka et al. demonstrated that Grm8 neurons form discrete clusters along the medial and ventral borders of the rodent NAc shell^35^. Further, registration with annotated NHP cell types ^24^ confirmed that D1 islands were molecularly distinct from dorsal striatum striosomes and instead showed greater similarity to the D1 NUDAP population. Within human D1 islands, we observed heterogeneity, with distinct *RXFP1* and *CPNE4* enriched islands, consistent with findings from He et al.^24^ who identified two transcriptionally distinct island subtypes in NHP accumbens. Collectively, these observations suggest that D1 islands represent an evolutionarily conserved and potentially functionally diverse compartment, where molecular heterogeneity may reflect specialization in processing opioid and neuromodulatory signals. Future studies combining high-resolution SRT with rodent or NHP connectivity mapping studies will be essential for refining the structural and functional delineation of these islands and for determining whether *RXFP1*+ and *CPNE4*+ D1 islands represent distinct opioid pathways.

The contribution of distinct NAc cell types and SpDs to neuropsychiatric risk is likely mediated by cell-cell signaling. Trait-associated ligand-receptor (LR) analyses identified neuromodulatory and opioid-related pathways such as *PDYN*–*OPRM1* linked to substance dependence risk. While DRD1_MSN_A neurons were predicted to be the primary *PDYN* senders, SRT data revealed that *PDYN* expression was highest in the medial MSN_3 SpD, where DRD1_MSN_A neurons contributed less compared to the lateral MSN_1 SpD based on cell-type deconvolution weights. This apparent discrepancy suggests heterogeneity within DRD1_MSN_A populations, with medial subpopulations potentially expressing higher levels of *PDYN* than their lateral counterparts and thereby disproportionately influencing dynorphinergic and opioid signaling. These findings underscore the importance of incorporating spatial context when modeling intercellular communication and align with prior functional studies demonstrating spatially segregated roles of dynorphin circuits. In particular, activation of dynorphinergic neurons in the ventral NAc shell induces aversive, dysphoric behavior, whereas dorsal shell activation promotes positive reinforcement^113^.

Consistent with the role of the NAc in modulating responses to opioids, we projected rodent-derived latent factors associated with morphine and cocaine exposure onto human SRT data. We found spatially distinct transcriptional signatures related to biological processes, such as stress response, synaptic plasticity, and neuromodulation rather than enrichment in specific MSN cell types or spatial domains. These findings highlight the latent susceptibility of multiple NAc spatial domains to either morphine or cocaine exposure. Interestingly, DRD2-associated transcriptional programs mapped onto D1 islands, suggesting shared adaptive responses across MSN subtypes rather than misassignment of cell identity. However, because many of these transcriptional responses are not cell type-specific, it is difficult to resolve which specific MSN subtypes or SpDs exhibit the strongest effects. This challenge is further compounded by the fact that our data is obtained from neurotypical individuals, and it is possible that genes not expressed in a particular cell type or SpD in neurotypical individuals could be robustly expressed in individuals with a SUD diagnosis. Important future directions include generating paired snRNA-seq and SRT datasets from human samples in the context of substance use disorders. Future studies could also establish an integrated spatial reference of the human NAc and computationally infer the anatomical positioning of cells from rodent or NHP case-control datasets, enabling the identification of drug-induced transcriptional changes within anatomically-defined cell types and SpDs. Finally, factorization-based transfer learning approaches could be broadly extended to integrate diverse datasets, including joint snRNA-seq and anterograde/retrograde connectivity tracing experiments, facilitating the integration of molecular, spatial, and functional information within a unified reference atlas.

In conclusion, these findings advance our understanding of the molecular and spatial organization of the human NAc, revealing how heterogeneous cell types, discrete SpDs, and continuous gradients interact within a complex anatomical framework. By integrating snRNA-seq and SRT data, we provide insights into how the structural organization of the NAc shapes neuromodulatory signaling, opioid sensitivity, and links to neuropsychiatric risk. Future directions such as high-resolution spatial profiling across the A-P axis, improved integration with case-control datasets, and connectivity mapping in rodents and NHP will be essential for resolving how transcriptional, spatial, and circuit-level characteristics converge to support the complex functions of the NAc in motivation and reward.

## 4 Methods

### 4.1 Postmortem human tissue samples

Postmortem human brain tissue from neurotypical adult donors of European ancestry (*N* = 10) were obtained at the time of autopsy following informed consent from legal next-of-kin, through the Maryland Department of Health IRB protocol #12–24, and from the Department of Pathology at Western Michigan University Homer Stryker MD School of Medicine, the Department of Pathology at University of North Dakota School of Medicine and Health Sciences, and the County of Santa Clara Medical Examiner-Coroner Office in San Jose, CA, all under the WCG protocol #20111080. Using a standardized strategy, all donors were subjected to clinical characterization and diagnosis. Macroscopic and microscopic neuropathological examinations were performed, and subjects with evidence of significant neuropathology were excluded. Additional details regarding tissue acquisition, processing, dissection, clinical characterization, diagnoses, neuropathological examination, RNA extraction and quality control (QC) measures have been previously published^114^. Demographic information for all neurotypical control donors included in transcriptomics studies is listed in Supplementary Table 1. Each tissue block was dissected from frozen coronal slabs at the level of the ventral striatum with a visible caudate nucleus, putamen, anterior limb of the internal capsule, and, in some specimens, optic chiasm and external globus pallidus (**Fig. S2, Fig. S3, Fig. S4**). Using a hand-held dental drill, tissue blocks of approximately 20 × 25 mm were dissected, encompassing the entire NAc. Tissue blocks were stored in sealed cryogenic bags at −80°C until cryosectioning.

### 4.2 Tissue processing and quality control

All tissue blocks were subjected to quality control by H&E to ensure dissected blocks contained the entire NAc and surrounding landmarks (caudate nucleus and internal capsule as the dorsal landmarks, white matter tracts of the forebrain as the ventromedial landmarks, and putamen/external globus pallidus as the lateral landmarks). Briefly, fresh frozen tissue blocks were acclimated at -14^°^C for 30 minutes inside the cryostat (Leica CM3050s), mounted on a round chuck with Optimal Temperature Compound (TissueTek Sakura, Cat #4583), and ∼50 µm of tissue was trimmed and discarded to achieve a flat surface. Several ∼10 µm sections were collected on pre-chilled microscope slides (VWR SuperFrost Microscope Slides, Cat #48311703). H&E staining was performed according to the manufacturer’s instructions as previously described ^57^, and images were acquired using an Aperio CS2 slide scanner (Leica). To prepare tissue for the 10x Genomics Visium assay, areas of the blocks corresponding to the NAc were scored with a razor blade vertically in ∼6.5 mm strips to match the width of the Visium capture areas. Because each NAc sample occupied 2-5 capture arrays, multiple non-overlapping, vertically-adjacent or partially-overlapping horizontally-adjacent 10 µm tissue strips containing NAc were mounted onto pre-chilled Visium Spatial Gene Expression slides (part number 2000233, 10x Genomics) to cover the entire NAc structure. Following successful completion of the Visium assay, to prepare tissue for snRNA-seq (10x Genomics Chromium), blocks were acclimated in the cryostat, trimmed to a flat surface, and 100 µm sections of the NAc were collected in a DNA LoBind 2 ml tube (Eppendorf, Cat#022431048) for a total of ∼100 mg of tissue. Tubes were stored at −80°C until the Chromium assay.

### 4.3 Spatially resolved transcriptomics (SRT) data generation

Visium Spatial Gene Expression slides were processed as previously described^57^. Tissue optimization experiments were performed according to the manufacturer’s protocol (CG000160, revision B, 10x Genomics) to ensure optimal permeabilization time. Briefly, eight NAc tissue sections were exposed to permeabilization enzymes for 3 to 36 min. cDNA synthesis was performed using a fluorescently labelled nucleotide (CG000238, revision D, 10x Genomics). The slide was then coverslipped and fluorescent images were acquired at 10x magnification with a TRITC filter (ex 550nm/em 600nm) on a Cytation C10 Confocal Imaging Reader (Agilent). The optimal permeabilization time of 18 min was selected for all subsequent experiments.

Two to five 10μm sections from each of the 10 NAc blocks were collected onto one or two 10x Visium Gene Expression slides, and processed according to the manufacturer’s instructions (10x Visium Gene Expression protocol number CG000239, Rev G) as previously described^57^. Briefly, H&E staining was performed (protocol CG000160, revision B, 10x Genomics), after which slides were coverslipped and high-resolution, brightfield images were acquired on a Leica CS2 slide scanner equipped with a 20x/0.75NA objective and a 2x doubler. Following the removal of the coverslips, tissue was permeabilized, cDNA synthesis was performed, and sequencing libraries were generated for all capture areas following the manufacturer’s protocol. Libraries were loaded at 300 pM and sequenced on a NovaSeq 6000 (Illumina) at the Johns Hopkins Single Cell Transcriptomics core according to the manufacturer’s instructions at a minimum depth of 60,000 read pairs per spot.

### 4.4 Single molecule fluorescence *in situ* hybridization (smFISH)

An independent non-neurotypical donor was obtained for smFISH experiments (Br4032; female; age 59.2 years; post-mortem interval (PMI) = 43 hours; RIN = 8.3). Fresh-frozen NAc tissue was sectioned at 10 µm, adhered to glass slides, and stored at −80°C. *In situ* hybridization was performed using the RNAScope Multiplex Fluorescent Reagent Kit v2 (Cat #323100, ACD, Hayward, California) as previously described^115^. To summarize, frozen tissue sections were fixed in 10% Neutral Buffered Formalin (NBF) solution (Ct # HT501128-4L, Sigma-Aldrich, St. Louis, Missouri) for 30 minutes at room temperature. A series of ethanol dehydration steps was performed followed by pretreatment with hydrogen peroxide for 10 minutes at room temperature and treatment with Protease IV for 30 minutes. The following probe combination was used to identify D1 islands: *Hs-MBP* (Cat #411051-C1), *Hs-RXFP1* (Cat #422821-C2), *Hs-DRD1* (Cat #524991-C3), *Hs-OPRM1* (Cat #410681-C4). Following probe labeling, tissue sections were stored overnight in 4× SSC buffer. Signal amplification was performed using AMP1–3 reagents, followed by fluorescent labeling with Opal dyes (1:500 dilution; Perkin Elmer, Waltham, MA) assigned as follows: Opal 520 for channel 1 (*MBP*), Opal 620 for channel 2 (*RXFP1*), Opal 690 for channel 3 (*DRD1*), and Opal 570 for channel 4 (*OPRM1*). Nuclear counterstaining was performed with 4′,6-diamidino-2-phenylindole (DAPI, ACD, Hayward, California) and slides were mounted with Fluoromount-G. Tissue sections were imaged using an AX Nikon Ti2-E confocal fluorescence microscope equipped with NIS-Elements software (AR6.10.01), utilizing a 2x objective (Nikon PLAN APO λ D 2x/0.1) and a 40x objective (Nikon Plan Apo Lambda S 40×/1.5). Images were spectrally unmixed to isolate and subtract the lipofuscin autofluorescence signal prior to analysis.

### 4.5 Single molecule fluorescence *in situ* hybridization with Immunofluorescence (smFISH-IF)

To combine smFISH with immunofluorescence (IF) staining, slides with frozen tissue sections (10 µm) were removed from –80°C and fixed in pre-chilled 10% NBF for 15 minutes at 4°C. Dehydration was performed sequentially in 50%, 70%, and two 100% ethanol incubations at room temperature. Hydrogen peroxide was applied for 10 minutes at room temperature to quench endogenous peroxidases. Primary antibodies anti-rabbit TUJ1 (1:500) (Cat # 18207, Abcam, Waltham, MA) and anti-chicken MAP2 (1:500) (Cat# 92434, Abcam, Waltham, MA) were diluted in co-detection diluent and applied to each section. Slides were incubated overnight at 4°C in a humidified chamber. The following day, sections were re-fixed in 10% NBF for 30 minutes at room temperature, washed, and treated with Protease IV for 30 minutes at room temperature. *SLC17A7*-specific (Cat #415611-C1, ACD, Hayward, California) RNAScope probes were applied and slides were incubated at 40°C for 2 hours. Signal amplification was performed using the RNAScope Fluorescent Multiplex V2 protocol, including sequential incubations with AMP1, AMP2, and AMP3 reagents at 40°C. HRP-C1 was used to detect and amplify the *SLC17A7* signal, followed by Opal dye 520 (1:500 dilution; Perkin Elmer, Waltham, MA) incubation for 30 minutes at 40°C. Secondary antibodies anti-rabbit Alexa Fluor 555 (Cat #A21429, Invitrogen, Waltham, MA) and anti-chicken Alexa Fluor 647 (Cat #A78952, Invitrogen, Waltham, MA), both 1:400, were applied for 40 minutes at room temperature in the dark. Slides were counterstained with DAPI (ACD, Hayward, California) and mounted with Fluoromount-G (Cat #00-4958-02, ThermoFisher). Final imaging was performed within 24 hours. Tissue sections were imaged using an AX Nikon Ti2-E confocal fluorescence microscope equipped with NIS-Elements software (AR6.10.01), utilizing a 2x objective (Nikon PLAN APO λ D 2x/0.1) and a 40x objective (Nikon Plan Apo Lambda S 40×/1.5). Images were spectrally unmixed to isolate and subtract the lipofuscin autofluorescence signal prior to analysis.

### 4.6 snRNA-seq data generation

Using 100 μm thick cryosections collected from each donor, we conductedsnRNA-seq using 10x Genomics Chromium Single Cell Gene Expression V3.1 technology. Approximately 70-100 mg of tissue was collected from each donor, placed in a pre-chilled 2 mL microcentrifuge tube (Eppendorf Protein LoBind Tube, Cat #22431102), and stored at −80°C until the time of the experiment (see Tissue Processing and Quality Control). Nuclei preparation was performed as previously described ^25^ Briefly, cryosections were incubated in chilled Nuclei EZ Lysis Buffer (MilliporeSigma #NUC101) in a glass dounce. Sections were homogenized using 10-20 strokes with both loose and tight-fit pre-chilled pestles. Homogenates were filtered through 70 μm mesh strainers and centrifuged at 500g for 5 minutes at 4°C using a benchtop centrifuge. Nuclei pellets were resuspended in fresh EZ lysis buffer, centrifuged again, and resuspended in wash/resuspension buffer (1x PBS, 1% BSA, 0.2U/μL RNase Inhibitor). Final nuclei were washed in wash/resuspension buffer and centrifuged a total of 3 times. For each donor, we split the total population of nuclei into two separate tubes to label half of the population with propidium iodide (PI) and the other half with both NeuN and PI. We labeled the latter population of nuclei with Alexa Fluor 488-conjugated anti-NeuN (MilliporeSigma cat. #MAB377X) diluted 1:500 in nuclei stain buffer (1x PBS, 3% BSA, 0.2U/μL RNase Inhibitor), by incubating at 4°C with continuous rotation for 1 hour. Proceeding NeuN labeling, nuclei were washed once in stain buffer, centrifuged, and resuspended in wash/resuspension buffer. All nuclei were labeled with PI at 1:500 in the wash/resuspension buffer and subsequently filtered through a 35 μm cell strainer.

We performed fluorescent-activated nuclear sorting (FANS) using a MoFlo Legacy Cell Sorter at the Johns Hopkins Flow Cytometry and Immunology Core. Gating criteria were selected for whole, singlet nuclei (by forward/side scatter), G0/G1 nuclei (by PI fluorescence), and neuronal nuclei (by Alexa Fluor 488 fluorescence). First, we sorted 14,000 nuclei based on PI+ fluorescence to include both neuronal and non-neuronal nuclei from each donor. We then sorted 14,000 additional nuclei into a separate tube based on both PI+ and NeuN+ fluorescence to enrich for neurons. This resulted in a final N=20 for snRNA-seq (one PI+ and one PI+NeuN+ sample for each donor). Samples were collected over multiple rounds, each containing 2-3 donors for 4-6 samples per round. All samples were sorted into reverse transcription reagents from the 10x Genomics Single Cell 3′ Reagents kit (without enzyme). Reverse transcription enzyme and water were added after FANS to bring the reaction to full volume. cDNA synthesis and subsequent library generation were performed according to the manufacturer’s instructions for the Chromium Next GEM Single Cell 3’ v3.1 (dual-index) kit (CG000315, revision E, 10x Genomics). Samples were sequenced on a Novaseq6000 (Illumina) at the Johns Hopkins University Single Cell and Transcriptomics Sequencing Core targeting 50,000 read pairs per nucleus.

### 4.7 snRNA-seq data processing and quality control

FASTQ files for snRNA-seq libraries from 20 samples were aligned to the human genome (GRCh38/Hg38, Ensembl release 98), using 10x Genomics software^116^, *cellranger count* v7.2.0. Raw feature-barcode files were analyzed in R (v4.3.2) using the Bioconductor^117,118^ suite of single-cell transcriptomics analytical packages. Droplets containing nuclei were identified via the `emptyDrops()` function from the *DropletUtils*^119,120^ package (v1.22.0) using a data-driven `lower` threshold. Quality control metrics were calculated using the `addPerCellQC()` function provided by the scuttle^121^ package (v1.12.0). High-quality nuclei were identified by assessing the total number of UMIs, number of detected features, and the percentage of reads mapping to the mitochondrial genome. This dataset contains samples sorted with PI+ to collect intact nuclei and PI+NeuN+ to enrich for neuronal nuclei. Neuronal nuclei contained more genes than non-neuronal nuclei and applying a single median absolute deviation (MAD) threshold to all samples would result in the loss of high-quality nuclei in neuronally enriched samples. To avoid discarding high-quality neuronal nuclei, the MAD thresholds were calculated using sort type (PI+NeuN+ vs. PI+) as the batch. For the total number of UMIs or the number of detected features, we applied a MAD threshold of 3 for PI+NeuN+ sorted samples and 1 for PI+ sorted samples. The `isOutlier()` function provided by the scater^121^ package v1.30.1 was used to perform QC with adaptive thresholds. When performing QC with the `isOutlier()` function, the `log=` argument was set to `TRUE` for library size and `FALSE` for the number of detected genes. Distributions for the percentage of reads mapping to the mitochondrial genome were centered on 0 and adaptive thresholds would result in the loss of several hundred high-quality nuclei. Thus, we discarded nuclei with >5% of reads mapping to the mitochondrial genome. In addition to the total number of UMIs, number of detected features, and percentage of reads mapping to the mitochondrial genome, preliminary dimensionality reduction and clustering identified a cluster of low-quality neurons with abnormally low numbers of detected features and total UMIs. These low-quality neurons were removed from the analysis. Following quality control of key metrics, we next performed doublet detection within each sample with the `computeDoubletDensity()` function provided by the scDblFinder^122^ package v1.16.0. Nuclei with a doublet score ≥5 were removed from further analysis. Following quality control, our dataset contained 36,601 genes and 103,785 nuclei. Nuclei contained an average UMI count of 17,304 and an average number of detected features of 4,634.

### 4.8 snRNA-seq feature selection, dimensionality reduction, and clustering

Following the removal of low-quality nuclei, feature selection was conducted by first fitting a binomial model to gene counts across all cells and calculating a deviance residual for each gene with the `devianceFeatureSelection()` function provided by the scry^123^ package (v1.14.0). Following the calculation of deviance statistics, Pearson residuals were calculated from the binomial model with `nullResiduals()` function provided by the scry^123^ package (v1.14.0). Genes were then ranked by decreasing deviance and the top 4,000 deviant genes, termed highly deviant genes (HDGs), were used for downstream dimensionality reduction. The binomial Pearson residuals of the 4,000 HDGs were then used for PCA and the top 100 principal components (PCs) were kept. Preliminary clustering analysis revealed the presence of sample-specific clusters. To correct for these batch effects, we ran *harmony*^*124*^ within the PC space with the sample ID as the batch via the `RunHarmony()` function from *harmony*^124^ (v1.2.0). Following batch correction, graph-based clustering was performed with `buildSNNGraph()` *from scran*^125^ (v1.30.0) with *k*=10 neighbors within the *harmony* corrected PC space. Following the construction of the shared nearest neighbor graph, cells were clustered using `cluster_louvain()` from *igraph*^*126*,*127*^ (v1.6.0) with a resolution value of 1, to yield 27 preliminary clusters. Following QC, dimensionality reduction, and clustering log_2_-normalized counts were finally calculated. First, preliminary clusters were used to calculate size factors with the `computeSumFactors()` function provided by *scran*^125^. Size factors were then used to calculate log_2_-normalized expression values with the `logNormCounts()` *function provided by scuttle*^121^. *To visually inspect clusters, we generated a t*-distributed stochastic neighbor embedding (*t*-SNE) using the top 100 PCs. Visual inspection of enrichment of literature-derived marker genes overlaid on the *t*-SNE identified that several preliminary clusters spanned single, previously identified cell types. To further investigate the relationship between clusters, pairwise modularity scores were calculated between preliminary clusters using the `pairwiseModularity()` function from *bluster*^128^ (v1.11.4) with `as.ratio=TRUE`. Additionally, preliminary cluster-specific marker genes were calculated by performing pairwise *t*-tests between each cluster with the `findMarkers()` function from *scran*^125^. First, related clusters were identified using pairwise modularity scores. Related clusters that were not marked by meaningful differences in literature-derived marker genes were merged. For example, preliminary clusters 1 and 2 had a high pairwise modularity score and were marked by *MBP, MOBP, ST18*, and *PLP1*, marker genes for oligodendrocytes. Thus, clusters 1 and 2 were merged and called Oligodendrocytes. Similarly, pairwise modularity suggested that preliminary clusters 3, 5, and 14 were related.

### 4.9 Identification of differentially expressed genes corresponding to underlying cell types in snRNA-seq data

Before performing non-pseudobulk differential expression (DE) testing, all genes with 0 counts for every cell were removed from further analysis. Additionally, cells assigned to the `Neuron_Ambig` cluster were removed before DE testing. Non-pseudobulk DE testing was performed as previously described^25^. Briefly, to identify cluster-specific marker genes, two separate methods were used to perform non-pseudobulk DE testing. First, pairwise *t*-tests were performed between each cluster with the `findMarkers()` function from *scran*^125^. The second method utilized the `findMarkers_1vAll()` function from *DeconvoBuddies*^129^ (v0.99) to perform cluster-versus-all-other-nuclei DEG testing. For both methods, a design matrix specifying the sample was included to control for any sample-specific gene expression differences that may affect DE testing.

### 4.10 *SRT* data processing and quality control

#### Visium raw data processing

Sample slide images were first processed using *VistoSeg* software^130^. *VistoSeg* was used to divide the Visium sample slides into individual images using the `splitSlide` function. This takes one large image and separates them into distinct capture areas, one for each area on the slide, labeled at A1, B1, C1, and D1. The individual images from *VistoSeg* were then aligned with the slide capture areas in the *Samui Browser* version (v1.0.0-next.24)^131^. This allows for the alignment of the image with the spots captured on the slide. Sample slides were then processed using the *SpaceRanger* (v2.1.0) software from 10x Genomics (https://www.10xgenomics.com/support/software/space-ranger/). Which takes the JSON output from the *Samui Browser*^*131*^, the sample image, and associated FASTQ files to generate spatial feature counts for a sample.

#### Image stitching

For each donor, the high-resolution images for all corresponding capture areas were loaded and manually aligned in *ImageJ*’s^132^ graphical user interface to match the relative tissue orientations of each capture area during imaging. *ImageJ* produces a JSON file containing a matrix corresponding to an affine transformation to apply to each capture area to produce the proper merged orientation for each donor. This matrix was read into *Python* 3.10, and the transformations were applied to the spatial coordinates stored in the `pxl_row_in_fullres` and `pxl_col_in_fullres` columns of *SpaceRanger*’s `tissue_positions_list.csv` file for each capture area (https://www.10xgenomics.com/support/software/space-ranger/). A combination of *NumPy* (v1.24.4)^133^ and *PIL* (v10.0.0)^134^ was used to also apply the transformations to the full-resolution H&E images, ultimately constructing one larger multi-channel image per donor such that each of a donor’s capture areas were reduced to grayscale and stored in unique channels. Using the Python API provided by *Samui Browser* (v1.0.0-next.24)^131^, the transformed images and spatial coordinates were loaded in the *Samui Browser*^131^, one donor at a time (**Fig. S5A**). Importantly, the *Samui Browser*^131^ enables viewing of full-resolution images, whereas memory constraints prevent full-resolution images from being possible to view in *ImageJ*^132^. This enables *Samui Browser*^131^ to have greater spatial precision in viewing image features, motivating its use in addition to *ImageJ*^132^ in this workflow. Once in the browser, the “ROI (region of interest) annotation” feature was used to label the centroids, vertices, or edges of shared histological features for each pair of overlapping capture areas, noting that a given capture area may be included in one or more pairs (**Fig. S5B**). A minimum of 4 pairs of features were selected by visual inspection for each overlap, to evenly sample points throughout the full region of overlap. These ROIs were exported to CSV using *Samui Browser*^131^ and imported in Python, where custom code was used to find the optimal rotations and translations of one capture area relative to the other in its pair (**Fig. S5C**). Here, an optimal adjustment is defined as one that minimizes the final average distance between the features in one capture area relative to the same features in its overlapping capture area. To find the optimal translation, a vector was computed for each shared feature, mapping from the location from one capture area to the other. The optimal adjustment was the mean of these vectors, depicted in the center (**Fig. S5C**). To find a rotation, line segments were found between each pair of features within each capture area. Each pair of features yielded an angle between the line segments; the optimal rotation angle was defined as a weighted average of the individual angles, where weights were given by the geometric mean of the intersecting line segments’ lengths (**Fig. S5C**). After determining the optimal adjustments, a higher-accuracy matrix of transformations was exported to CSV, much like the matrix contained in the JSON output from *ImageJ*^132^. This enabled the same Python-based workflow to be *executed* again^133,134^, adjusting spatial coordinates and each capture area’s images with greater precision; the *Samui Browser*^131^ API was again used to visually verify the improvements at full resolution. Finally, an RGB version of the merged image was downscaled to “high resolution” SpaceRanger-compatible images (max 2,000 pixels on the X or Y axis) and saved for downstream use in a *SpatialExperiment* object (v1.10.0)^135^.

#### Defining array coordinates for merged images

Following Visium’s hexagonal array design for a single capture area, `array_row` and `array_col` (“array coordinates”) are columns in *SpaceRanger*’s `tissue_positions_list.csv` file representing each Visium spot’s row and column index (https://www.10xgenomics.com/support/software/space-ranger/). By definition, `array_row` holds integer values from 0 to 77, while `array_col` holds integer values from 0 to 127. After performing image stitching (Materials and Methods: Image stitching), there is a need for new array coordinates relative to the new stitched image. The original array coordinates cannot reflect how capture areas may have been arbitrarily rotated and translated to reconstruct the true tissue arrangement across the capture areas. To redefine these array coordinates, we employed an algorithm that assigns each whole tissue section a new Visium-like array, in place of the individual capture areas that comprise the tissue section (**Fig. S6**). Like a true Visium array, each “artificial spot” is a distance of 100 microns from each of its neighbors; array rows and columns are defined such that the minimum and maximum values for `pxl_row_in_fullres` and `pxl_col_in_fullres` of any capture area for one donor correspond to values of 0 for `array_col` and `array_row`, respectively (**Fig. S6**) as in the original `tissue_positions_list.csv` file. Array rows and columns are added as necessary, generally well above the ordinary respective maximum values of 77 and 127, to cover the area spanned by all original spots for a whole tissue section. Each original spot is then mapped to the nearest artificial spot by Euclidean distance to redefine the `array_row` and `array_col` values used downstream in clustering via *PRECAST* (v1.6.3)^52^. In the default implementation for Visium data, *PRECAST*^52^ *uses array coordinates for finding a spot’s neighbors but is agnostic to the fact that more than one artificial spot may occupy the same array coordinates, as occurs in regions of overlap between capture areas*.

#### Determining overlapping spots in Visium arrays

For select donors such as Br6432, we observed that a particular tissue region was repeated in more than one capture area. We defined spots which belonged to distinct capture areas which mapped to the same tissue position as overlapping spots and considered them to be technical replicates. To identify these overlaps, all capture arrays for each donor were aligned using the refined affine transformations derived during image stitching (Materials and Methods: Image stitching). Overlapping spots were identified using the *visium_stitcher* (v0.0.3) package^136^, and a spot was marked overlapping if it fell within a distance of 1. 5 × the median nearest neighbor distance of an existing spot from the first listed slide’s native spot spacing. We then determined the optimal order of merging by comparing the UMI counts for all possible pairs of overlapping capture arrays, and prioritized capture arrays in which the overlapping spots had a higher minimum UMI count, while marking spots from the lower-depth capture array for exclusion by creating an `exclude_overlapping` flag. Following this QC-based prioritization, all capture arrays for a particular donor were merged using visium_stitcher()^136^ along with annotations of which overlapping spots were of lower quality.

#### Constructing a SpatialExperiment object

First, `read10xVisiumWrapper()` from *spatialLIBD* (v1.15.1)^58^ was used to read the gene expression data, images, and spatial coordinates from the outputs of Spaceranger, run on each capture area, ultimately producing a *SpatialExperiment*^135^ object. Next, the transformed spatial coordinates and donor-level, high-resolution merged images from the image-stitching process were read in (Materials and Methods: Image stitching). These donor-level (whole tissue section) images were read into R with `SpatialExperiment::readImgData()`^135^, overwriting the capture-area-level images from `spatialLIBD::read10xVisiumWrapper()`^58^. Conceptually, each sample in the *SpatialExperiment*^135^ is one donor; in practice, this meant that the `sample_id` variable in the `colData()` of the *SpatialExperiment*^135^ referred to donor, coordinates stored in the `spatialCoords()` slot were the transformed values from image stitching, the `imgData()` slot contained donor-level images, and the `array_row` and `array_col` columns in the `colData()` were redefined to encompass entire tissue sections (Materials and Methods: Defining array coordinates for merged images). However, `colData()` columns called `sample_id_original`, `pxl_row_in_fullres_original`, `pxl_col_in_fullres_original`, `array_row_original`, and `array_col_original`, were retained, referring to the capture-area level equivalents read in directly from `read10xVisiumWrapper()`^58^. To visualize this stitched data, we developed custom R code^37^ that is now available through *spatialLIBD*^58^ >=v1.17.3 by supplying the `is_stitched = TRUE` argument to `vis_gene()` or `vis_clus().`

#### Visium quality control and count normalization

Our initial data consisted of 189,661 Visium spots and preliminary QC led to the exclusion of 50 spots with no detected gene expression, and 11,815 spots which were not in tissue regions, leaving 177,804 spots for further QC. For the remaining spots, we computed per-spot QC metrics including library size (`sum_umi`), number of detected genes (`sum_gene`), proportion of mitochondrial gene expression (`expr_chrM_ratio`), and nuclei counts from VistoSeg^130^ (`Nmask_dark_blue`). To assess data quality, we visualized the distribution of key QC metrics using both spatial spot plots (**Fig. S12, Fig. S13**) and global distributions across all spots, individual capture arrays, and donors (**Fig. S14, Fig. S15**). We observed that both `sum_umi` and `expr_chrM_ratio` were systematically lower in white matter when compared to gray matter, consistent with previous studies^57^ showing reduced RNA content in white matter, and higher mitochondrial activity in neurons^137^. Because these biologically driven differences could confound automated outlier detection, we avoided using global statistical cutoffs for low-expression filtering. Instead, we identified low-quality spots by using a combination of expression and spatial criteria. Specifically, 1,214 spots with `sum_umi` < 250 and `edge_distance` < 6 were excluded from downstream analysis. The `edge_distance` was estimated per capture area using the untransformed `array_row` and `array_col` attributes and computing the euclidean distance, as was done previously^138^. Additionally, we detected 584 local expression outliers by using SpotSweeper^139^ (v0.99.1), which leverages local neighborhood-based similarity to identify expression outliers. In summary, we retained a total of 176,013 spots across 10 donors, with a median of 18,536.5 spots per donor. Br8667, was the donor with the fewest number of spots, which was 8,747. The median number of spots per capture area was 4,674.5.

After spot-level QC, we removed genes with zero expression in any donor, to only include features that were detected across all samples, resulting in 22,375 genes retained. For normalization, we used the *scuttle*^*121*^ framework. First, per-spot size factors were computed using `computeLibraryFactors()`, which accounts for differences in sequencing depth. These size factors were then used to perform log-transformation of normalized counts with `logNormCounts()`, yielding normalized expression suitable for downstream dimensionality reduction and spatial domain identification.

### 4.11 SRT feature selection and unsupervised identification of spatial domains

#### Finding spatially variable genes

*nnSVG* (v1.4.2)^51^ was used to find spatially variable genes (SVGs) for each donor. First, we subset the *SpatialExperiment*^135^ object which was processed as detailed in **Materials and Methods: SRT data processing and quality control** to only include spots from a single donor. We then performed QC on the selected spots by excluding overlapping spots, and running `filter_genes()` from *nnSVG*^51^. Default parameters that retained non-mitochondrial genes with at least 3 counts in 0.5% of spots were used. Next, spots with no gene counts were dropped. Log-normalized counts were re-computed on the filtered data using `computeLibraryFactors()`, followed by `logNormCounts()` from *scuttle*^121^.

We then ran `nnSVG()`^51^ in two stages. In the first stage, we used default parameter values and generated SVGs which were used as input for *PRECAST*^52^ (v1.6.3) clustering with k=2 (**Materials and Methods: Clustering with PRECAST**). *PRECAST*^52^ (k=2) cluster assignments effectively partitioned white and gray matter (**Fig. S16**). However, several top SVGs included genes that corresponded to broad differences between white and gray matter instead of capturing expression heterogeneity that occurs within these domains as previously observed in brain tissue^57,138,140^ (**Table S4**). Thus, in the second stage, we included the cluster assignments from *PRECAST*^52^ (k=2) as a covariate while detecting SVGs, via `nnSVG(X=model.matrix (∼precast_cluster))`^51^. For both stages of identifying SVGs, the individual donor-level SVGs were aggregated into a summary list of SVGs that was applicable to the entire dataset. First, SVGs were filtered to only retain genes with significant spatial variability across all donors (`padj < 0.05`). Next, the `rank` metric for each SVG was averaged across all donors, and SVGs were re-ranked according to this mean value (**Table S4**). The top 2,000 genes based on aggregate rank were used as input features for downstream clustering and identification of spatial domains (SpDs).

#### Unsupervised clustering of spatial transcriptomics data

*PRECAST* (v1.6.3)^52^ was employed to jointly cluster spots across all donors. The *SpatialExperiment*^*135*^ object (**Materials and Methods: Constructing a SpatialExperiment object**) was individually subset to each donor to form a list of *Seurat* (v5.0.0) objects^141–145^, expected by the `seuList` parameter to `CreatePRECASTObject()`from *PRECAST*^52^. The `row` and `col` columns in the metadata of each donor’s *Seurat* object were set to the redefined donor-level (whole tissue section) `array_row` and `array_col` values (**Materials and Methods: Defining array coordinates for merged images**). We used the top 2,000 SVGs identified from *nnSVG*^51^ (**Materials and Methods: Finding spatially variable genes**) as input features via the `customGenelist` parameter, while also setting `selectGenesMethod=NULL`. All other gene-filtering parameters were set to their default values, while noting that the provided SVGs had been similarly filtered. Importantly, *PRECAST*^52^ was run in two configurations. For k=2, we used SVGs that were obtained using default settings, since these genes captured global expression patterns separating regions of white and grey *matter*. For k ≥ 3, we used SVGs obtained after adjusting for *PRECAST*^52^ *k*=2 cluster assignments, which better captured finer spatial heterogeneity within white and gray matter. For clustering, `AddAdjList (platform = “Visium”)` was used to treat each donor as a Visium sample. This resulted in PRECAST^52^ identifying more than six neighbors for a spot, since more than one spot may occupy identical array coordinates. Next, `AddParSetting(Sigma_equal = FALSE, maxIter = 30)` was invoked, following recommendations provided in the software documentation (https://feiyoung.github.io/PRECAST/articles/PRECAST.BreastCancer.html)^52^, and finally `PRECAST()` was run for all integer values of k between and including 2 and 15. To evaluate the robustness of spatial clustering, we performed five independent random starts for each k using distinct seeds.

#### Pseudobulk differential expression analysis

For each random initialization and clustering resolution (k), we performed pseudobulk aggregation to identify differentially expressed genes (DEGs) across the inferred unsupervised clusters. The processed SpatialExperiment^135^ object was loaded and cluster assignments were appended to the `colData()`. Spots which did not have a corresponding cluster assignment (`NA`) were discarded. Pseudobulk expression profiles were generated using `aggregateAcrossCells()` from the scuttle^121^ R package as previously described^146^, and counts were grouped by both *PRECAST*^52^ cluster and capture area and summed. This approach preserved within-donor spatial variation while reducing technical noise from individual spots. Metadata including donor identity, capture area, age, sex, diagnosis, and the number of aggregated spots was retained for downstream modeling. Next, pseudobulked samples obtained by aggregating across fewer than 50 spots were excluded. Genes were then filtered by using `filterByExpr()` from the edgeR R package (v4.0.3)^147^ considering both cluster-level and capture-area level grouping and default parameters. Only genes which were not found to be expression outliers at the cluster-level were retained for DEG testing. Pseudobulk samples were then normalized using `calcNormFactors()` from the edgeR R package^147^ with the TMM method^148^ and log-transformed using `cpm()` from edgeR^147^ with a `prior.count=1`. Finally, pseudobulk samples with fewer than 2,000 detected genes and clusters with fewer than 10 pseudobulked samples were removed. DEG testing was performed using the registration framework from spatialLIBD^58^. First, we constructed a design matrix with `registration_model()`^58^ and included donor sex and slide number as covariates. We did not include donor age as a covariate, since it was linearly dependent with sex and slide number. We accounted for sample-level correlation using `registration_block_cor()`^58^, treating the capture array variable as a blocking factor. Next, we performed differential expression testing using three approaches, including, (1) one-vs-all enrichment tests (`registration_stats_enrichment`), (2) pairwise comparisons (`registration_stats_pairwise`), and (3) ANOVA tests (`registration_stats_anova`) when k ≥ 3. Significance was determined at FDR < 0.05, and a list of significant genes were obtained using `sig_genes_extract_all()`.

#### Spatial domain annotation

*The PRECAST*^52^ clustering resolution (k) and random initialization were selected prior to spatial domain annotation. Across k=3-15 and five random starts, the log-likelihood was directly derived from *PRECAST*^52^ outputs (**Fig. S17A**), and the degree of freedom (df) was estimated based on the number of spatial domains (k), latent dimensions (q), number of genes (p), the rank of low-dimensional embeddings (r_max), and variance-covariance structures used by PRECAST^52^ in inferring the clusters. The BIC was then computed as follows,

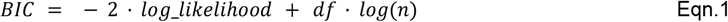

where *n* is the total number of spots. Ideally, the clustering resolution which minimizes BIC corresponds to the optimal trade-off between model fit and model complexity^53^. Based on the BIC, several candidate resolutions emerged (**Fig. S17B**). To further narrow these down, we examined the concordance in cluster assignments between overlapping spots which can be considered technical replicates. Concordance decreased with increasing k by 1 until k = 9 (**Fig. S17C**), and then began to increase again. Among the candidate solutions, k=10 provided the best balance between BIC-based model fit, technical replicate concordance, and biological interpretability, as it was the smallest resolution at which D1 islands could be reliably resolved using previously established marker genes. To account for variability across multiple random initializations, we computed the Rand index^149^ between the cluster assignments across the five random starts for a resolution of k=10, and found random start 3 displayed the highest overall concordance showing agreement with random start 1 (Rand index = 0.67) and random start 2 (Rand index = 0.64) (**Fig. S18B**). Based on these findings, the cluster assignments from random start 3, and k=10 were selected for downstream analyses including domain annotations, DEG testing, and spatial registration. To obtain biologically interpretable spatial domains (SpDs) from cluster assignments, *PRECAST*^52^ clusters were merged and annotated based on cluster-specific DEGs obtained from the one-vs-all tests described in **Materials and Methods: Pseudobulk differential expression analysis**. Canonical marker gene expression was examined to merge clusters based on transcriptional similarity. First, clusters 6 and 10 were merged into a single endothelial/ependymal domain based on the enrichment of vascular and choroid plexus genes including *CLDN5* and *TTR*. Similarly, clusters 2 and 9 were combined into a unified white matter (WM) domain based on elevated expression of oligodendrocyte- and myelination-associated genes such as *MBP, MOG*, and *PLP1*. After merging, eight SpDs were identified, including MSN_1-3, D1 islands, inhibitory, excitatory, endothelial/ependymal, and WM. Finally, DEGs specific to each SpD were identified using pseudobulk-based DEG testing described in **Materials and Methods: Pseudobulk differential expression analysis**.

### 4.12 Spatial registration of SRT SpDs to human NAc cell types

To assign putative cellular identities to the SpDs identified in **Materials and Methods: SRT feature selection and unsupervised identification of spatial domains**, spatial registration was performed by correlating SpD-level differential expression signatures with cell type-specific transcriptional profiles derived from paired human NAc snRNA-seq data as previously described by Huuki-Myers et al.^57^. A pre-processed *SingleCellExperiment*^118^ object containing cell type annotations (`CellType.Final`) generated as described in **Materials and Methods: snRNA-seq feature selection, dimensionality reduction, and clustering** was used as the reference, and all cells labeled `Neuron_Ambig` were excluded.

Spatial registration was performed using two complementary frameworks. In the neurons-only registration, only annotated neuronal populations were retained, including DRD1 and DRD2 MSNs, excitatory and inhibitory neurons. In the all-cell types registration, all annotated cell types were included, including glial, endothelial, and ependymal cells.

First, for both frameworks, cell type-specific enrichment statistics were computed using the `registration_wrapper()` from *spatialLIBD* (v1.17.8)^58^, using `CellType.Final` as the registration variable. This function facilitated convenient preprocessing of snRNA-seq data by aggregating pseudobulk counts per donor, performing quality control and normalization, and conducting one-vs-all differential expression tests adjusted for age and sex. The resulting enrichment model produced *t*-statistics for each gene across all annotated cell types, generating transcriptional profiles which served as reference signatures for spatial registration.

For integration with SRT-derived SpDs, SpD-specific enrichment *t*-statistics were obtained from pseudobulk differential expression modeling (**Materials and Methods: Pseudobulk differential expression analysis**). Pearson correlation coefficients were then computed between SpD-level and cell-type-level enrichment *t*-statistics using `layer_stat_cor(model_type=“enrichment”, top_n=100)` from *spatialLIBD*^58^. The union of the top 100 marker genes per cell type was used to focus the registration on the most informative features. Separate correlation matrices were generated for the neuron-restricted and all-cell type analyses, which were reordered for interpretability based on a curated hierarchy of SpDs and cell types. Visualization was performed using ComplexHeatmap (v2.18.0)^150,151^, producing heatmaps showing transcriptional similarity between SpDs and cell types.

### 4.13 Cross-species registration to map human NAc cell types and spatial domains to rodent and NHP NAc cell types

#### Cross-species registration of human NAc snRNA-seq data

To assess the conservation of human NAc cell types across species, cross-species registration was performed using reference NAc snRNA-seq data from rodents^26^ and non-human primates (NHPs)^24^. Since cross-species comparisons that include all annotated cell types are often dominated by global transcriptional patterns reflecting broad differences between neuronal and non-neuronal populations, the analysis was performed in two complementary modes: (1) using all annotated cell types and (2) restricting to neuronal cell types only. This dual framework enabled the characterization of broad transcriptional conservation and finer resolution mapping of related neuronal subtypes. The cell types present in the rodent and NHP reference datasets used are summarized below.

- *Rodent data*: Cell types include multiple DRD1 MSN subtypes (Drd1-MSN-1, Drd1-MSN-2), DRD2 MSN subtypes (Drd2-MSN-1, Drd2-MSN-2), Drd3-MSN, Grm8-MSN, Glutamatergic neurons, GABAergic neurons, Pvalb-, Sst-, and Chat-interneurons, as well as non-neuronal cell populations such as astrocyte, microglia, polydendrocyte, and mural cells.
- *NHP data*: Two reference datasets were obtained from GEO (GSE167920)^24^. The first, `GSE167920_Results_full_processed_final.rds` contained annotations spanning both neuronal and non-neuronal populations including, DRD1 MSNs, DRD2 MSNs, interneurons, astrocytes, microglia, oligo precursor cells, oligodendrocytes, mural/ fibroblast, and endothelial cells. The second dataset, `GSE167920_Results_MSNs_processed_final.rds` focused exclusively on MSNs, and included populations such as D1/D2 Matrix, D1/D2 Striosome, D1/D2 Shell/OT, D1 NUDAP, D1 ICj, and D1/D2 Hybrid.

For the neurons-only analysis, the human dataset was processed differently depending on the reference species to match the resolution of the available annotations. When comparing to the rodent reference, the human dataset was subset to include DRD1 and DRD2 MSNs, excitatory neurons, and six inhibitory interneuron subtypes (Inh_A-Inh_F) to match the boarder neuronal diversity represented in the rodent reference. In contrast, when aligning to the NHP data, the human dataset was restricted to only include the six MSN subtypes.

First, for each mode, DEGs were identified within the human snRNA-seq data using the cell-type enrichment model (`registration_stats_enrichment`) described in **Materials and Methods: Spatial registration of SRT SpDs to human NAc cell types**. Within each annotated cell type, genes were filtered based on FDR < 0.05 and logFC > 0, ranked by logFC, and the top 250 DEGs were selected. The union of these genes across all selected cell types was then used to define the feature set for cross-species mapping. Next, the gene symbols for the filtered gene set were mapped to their respective rodent and NHP homologs to enable cross-species comparisons. For rodent mappings, homologous relationships were identified using the `HOM_AllOrganism.rpt` file from the Jackson Laboratory MGI database (https://www.informatics.jax.org/downloads/reports/index.html), retaining only unique one-to-one human-rat gene matches. For NHP mappings, orthologs were identified using the `orthologs()` function from the *babelgene* R package (v22.9)^152^ function with `species = “Macaca mulatta”`, producing human-macaque gene correspondences. Following ortholog mapping, the final DEG set was filtered to include only genes expressed in both the human and reference datasets. Finally, for each species and analysis mode, we computed the *t*-statistics for each gene *g* and cell type *c* using a one vs. all framework previously used by Phillips et al.^26^ and described in Eqn. 2.

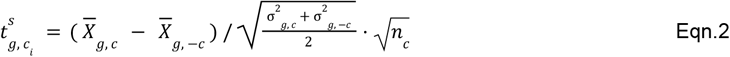

Where, 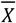 is the mean log-normalized gene expression, σ^2^ is the variance of the log-normalized expression, and *n* is the number of nuclei assigned to cell type *c*. To quantify transcriptional similarity between species, Pearson correlation coefficients were computed between the human *t*-statistics and the corresponding rodent or NHP *t*-statistics across the union of orthologous DEGs for each pair of cell types. The resulting correlation matrices were reordered according to curated hierarchies of cell types within each species to enhance interpretability and visualized using *ComplexHeatmap*^150,151^. Two sets of heatmaps were generated per species: one summarizing transcriptional similarity across all annotated cell types and another restricted to neuronal populations only.

#### Cross-species registration of human NAc SRT data

To map human NAc SpDs identified from SRT data to transcriptionally corresponding cell populations in other species, SpD-specific transcriptional signatures were compared against cell type-specific signatures obtained from rodent^26^ and NHP^24^ datasets. This analysis mirrored the snRNA-seq cross-species framework previously described in **Materials and Methods: Cross-species registration of human NAc snRNA-seq data**, with the key distinction that SpD-level enrichment statistics from SRT pseudobulk modeling (**Materials and Methods: Pseudobulk differential expression analysis**) were used instead of cell type-specific DEGs. For each SpD, DEGs were identified using the one-vs-all enrichment tests (FDR < 0.05, logFC > 0), ranked by logFC, and the top 250 genes per SpD were selected. Gene symbols from the union DEG set were mapped to their respective rodent and NHP orthologs as described previously (**Materials and Methods: Cross-species registration of human NAc snRNA-seq data**), retaining only one-to-one matches, and further filtered to only include genes expressed in both the SRT and reference datasets. For each SpD, *t*-statistics were computed using the approach described in Eqn. 2 and were correlated with rodent or NHP cell-type *t*-statistics derived from their respective snRNA-seq datasets. Pearson correlation coefficients were calculated across the shared ortholog set to quantify transcriptional similarity between each SpD and each reference cell type.

To capture both global and fine-grained patterns of transcriptional similarity between domains and reference cell types, the analysis was conducted in two modes: (1) All SpDs, and (2) restricting to neuronal SpDs. For rodent comparisons, the neuronal mode included MSN_1–3, D1 islands, excitatory, and inhibitory SpDs to match the broader neuronal diversity represented in the reference dataset. For NHP comparisons, the neuronal analysis was restricted to MSN_1–3 and D1 islands, reflecting the MSN-specific annotations available in the NHP dataset. Correlation matrices were reordered based on curated SpD and reference hierarchies to improve interpretability and visualized using *ComplexHeatmap*^150,151^.

### 4.14 *MERINGUE* to uncover latent factors of spatial variation in MSN domains

Discrete spatial clustering using *PRECAST*^52^ identified three MSN SpDs (MSN_1–3), but variability in SpD boundaries and small log fold-changes of SpD-specific DEGs suggested the presence of continuous molecular variation rather than clearly distinct compartments. To investigate these gradients we applied *MERINGUE*^59^ (v1.0), a spatial factorization framework which identifies latent spatial gene expression patterns by leveraging spatial autocorrelation and local neighborhood structure.

#### *Inferring MERINGUE* factors in each donor

For each donor, *MERINGUE*^59^ was run independently using only Visium spots assigned to MSN SpDs. Spatial neighbors were defined using the `getSpatialNeighbors()` function in *MERINGUE*^59^, which constructs a binary spatial adjacency matrix (*W*) by thresholding pairwise Euclidean distances between transformed Visium spot coordinates (**Materials and Methods: SRT data processing and quality control**). A distance cutoff of 500 units in the transformed coordinate space was applied, such that pairs of spots within this distance were considered neighbors (*W*_*i*,*j*_ = 1), while more distant pairs were set to *W*_*i*,*j*_ = 0. This resulted in a median number of 6 neighbors per spot. Spatially autocorrelated genes were identified using `getSpatialPatterns()`, which computes Moran’s I statistics^153^ from log-normalized SRT expression to quantify the similarity of gene expression between neighboring spots, where higher values indicate genes forming spatially gradients. Genes showing significant spatial autocorrelation were retained using `filterSpatialPatterns()`, which performs a Local Indicator of Spatial Association (LISA) test^154^ and selects genes such that at least 5% of spots significantly contribute to the spatial pattern (FDR < 0.05). Using this filtered set of spatially patterned genes, we computed the spatial cross-correlation (SCC) matrix to quantify similarity between spatial patterns of individual genes. Intuitively, the SCC measures how strongly two genes co-vary across space while accounting for the spatial neighbor structure defined by *W*. If two genes are consistently co-expressed in nearby spots, the corresponding SCC value will be high. This SCC matrix was then used as input to `groupSigSpatialPatterns()`, which hierarchically clusters genes with similar spatial patterns. Finally, pattern scores were obtained by z-scoring each gene’s expression across spots, averaging the standardized expressions of all genes within a gene cluster, and using Akima interpolation^155^ to fill in empty regions.

#### Grouping factors across donors to obtain *MERINGUE* consensus patterns

After identifying donor-specific *MERINGUE*^59^ factors, we introduce a new approach to integrate these patterns across donors by defining *MERINGUE* consensus patterns (MCPs) that capture reproducible transcriptional gradients. For each donor, we first extracted the set of genes assigned to each *MERINGUE*^59^ *f*actor from `groupSigSpatialPatterns()`. Gene-level similarity between *MERINGUE*^59^ factors was quantified using the Jaccard index. A Jaccard index close to 1 indicates a strong overlap in spatially patterned genes between two factors, whereas a value near 0 reflects minimal similarity (**Fig. S22**). In parallel, we estimated domain-level similarity by computing the average factor scores within each MSN domain (MSN_1–3) and correlating these domain averaged profiles across patterns (**Fig. S23**). This captures whether a particular pair of *MERINGUE*^59^ factors inferred across distinct donors are enriched across similar MSN domains. We then combined the gene-level Jaccard similarities and domain-level correlations into a consensus similarity matrix (**Fig. S24**), calculated as the mean of the two similarity metrics. Hierarchical clustering using Ward’s method implemented in `hclust()`^156^ was performed on the consensus similarity matrix to construct a dendrogram relating *MERINGUE*^59^ factors. The resulting tree was cut into discrete groups using `cutree()`^157^ with k = 6, yielding four reproducible MCPs shared across donors. To avoid redundancy from multiple *MERINGUE*^*59*^ factors inferred in the same donor within a single MCP, we retained at most one representative factor per donor per MCP. Representative factors were chosen by selecting MCP-donor pairs with exactly one contributing factor. The only exception to this rule was the inclusion of Br6432_Pattern_3 in MCP3 which facilitated comparisons of spatial biases exhibited by the MCPs across the A-P axis.

### 4.15 Spot deconvolution with Robust Cell Type Deconvolution (*RCTD*)

The 10x Visium platform quantifies gene expression from spots that typically contain mixtures of multiple nuclei, necessitating computational deconvolution to infer cell-type composition at the spot-level resolution.To map the heterogeneous spatial organization of neuronal and non-neuronal populations across transcriptionally defined domains, we applied *RCTD* from the *spacexr* R package (v2.2.1)^64^. We selected *RCTD*^64^ because previous benchmarking studies^146^ demonstrated that *RCTD*^64^ achieves the highest concordance with immunofluorescence-based cell counts when compared to other deconvolution methods including *Tangram*^158^ and *cell2location*^159^. Intuitively, *RCTD*^64^ integrates paired snRNA-seq and SRT data by fitting a poisson-lognormal hierarchical model that accounts for gene-specific platform effects thereby facilitating the accurate estimation of cell-type weights for each Visium spot.

#### Identification of marker genes and generation of *RCTD* reference

snRNA-seq data from all 10 donors was jointly processed to identify marker genes for each cell type and generate a unified reference for *RCTD*^64^ deconvolution. A processed *SingleCellExperiment*^118^ object containing curated cell type annotations (**Materials and Methods: Identification of differentially expressed genes corresponding to underlying cell types in snRNA-seq data**) was imported and nuclei labeled `Neuron_Ambig` were excluded from downstream analysis. Gene identifiers were converted to HGNC symbols, and normalized log-transformed counts were re-computed using `logNormCounts()` function from the scuttle^121^ R package. Ribosomal and mitochondrial genes were excluded prior marker gene identification. Marker genes used in deconvolution were identified using the `scoreMarkers()` function from *scran*^125^. Genes with mean AUC > 0.75 are selected as robust markers, sorted by decreasing AUC, and combined across all cell types to create a unified set of highly informative features. This curated marker set was used to construct a reference object using the `Reference()` function from *RCTD*^64^, which includes filtered counts, cell-type annotations, and total UMI.

#### Pre-processing of SRT data and PRECAST model initialization

Previously processed SRT data (**Materials and Methods: SRT data processing and quality control**), containing filtered raw counts was imported and reformatted by converting gene identifiers to gene symbols and standardizing gene symbols using `make.names()` to ensure uniqueness. Donor-specific datasets were then generated by subsetting the *SpatialExperiment*^135^ object based on the `sample_id` variable, retaining only SRT spots corresponding to each donor. From each donor-specific dataset, raw UMI counts, spatial coordinates (`spatialCoords`), and total UMI counts per spot were extracted to construct a *SpatialRNA* object using *spacexr*^64^. *Each SpatialRNA* object was paired with the marker-gene-restricted snRNA-seq reference to initialize a RCTD model using the `create.RCTD()` function. We specified `UMI_min = 2` to exclude spots with low gene expression and `MAX_MULTI_TYPES = 5` to allow for the model to consider up to five potential contributing cell types per spot.

#### Cell type deconvolution and visualization

*RCTD*^64^ inference of cell-type weights was run independently in each donor. The previously initialized *RCTD* object was loaded, followed by deconvolution using the `run.RCTD()` function from the *spacexr* R package (v2.2.1)^64^. We ran *RCTD*^64^ in multi doublet mode (`doublet_mode = “multi”`), which enables the model to penalize overfitting. During inference, *RCTD*^64^ integrates the spot-level SRT expression matrix with the paired snRNA-seq-derived reference to estimate the posterior probabilities of cell-type contributions per spot. Following *RCTD*^64^ deconvolution, estimated cell-type weights were extracted from the all-weights matrix (`results$all_weights`), representing the fractional contribution of each cell type found in the reference to each spot. These weights were appended to the `colData()` of the processed *SpatialExperiment*^135^ object and visualized using the `vis_gene()` function from *spatialLIBD*^58^. In addition to spot plots, domain-aggregated *RCTD*^64^ weights were computed by averaging spot-level weights within each spatial domain, and the proportion of spots in each domain with non-zero weights was estimated.

### 4.16 Integration of snRNA-seq with SRT using non-negative matrix factorization

Non-negative matrix factorization (NMF)^160^ is an unsupervised dimensionality reduction approach that decomposes a non-negative gene expression matrix *A* (genes x nuclei), into two lower rank non-negative matrices *W* and *H* as shown in Eqn. 3.

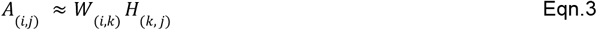

Here, *k* specifies the factorization rank and represents the number of latent transcriptional programs that are inferred from the data. The matrix *W* captures the contribution of each gene to a particular transcriptional *program*, whereas *H* represents the activation of these programs across nuclei. Importantly both *W* and *H* are constrained to only include non-negative values, which enables an additive parts-based representation of the data. In the context of snRNA-seq data, NMF enables the discovery of *de novo* transcriptional programs without relying on discrete predefined cell types or marker genes. Previous studies have successfully leveraged NMF to infer continuous transcriptional programs from snRNA-seq data and integrate these with spatially resolved transcriptomics (SRT), thereby linking single-nucleus resolution with spatial context ^146^. Building on this framework, we applied NMF to the human NAc snRNA-seq data to identify latent transcriptional programs and projected the resulting factor loadings onto our SRT data.

#### Pre-processing snRNA-seq and SRT data

A processed *SingleCellExperiment*^118^ object containing count data and cell type annotations (**Materials and Methods: Identification of differentially expressed genes corresponding to underlying cell types in snRNA-seq data**) was loaded, and nuclei labeled `Neuron_Ambig` were excluded prior downstream analysis. To ensure compatibility with the SRT data, analysis was restricted to genes expressed in both the snRNA-seq data and the pre-processed *SpatialExperiment*^135^ object containing the SRT data (**Materials and Methods: SRT data processing and quality control**). Following gene-filtering, snRNA-seq and SRT counts were log-normalized using `LogNormalize()` from Seurat (v5.0.1)^141–145^ with a scaling factor of 10,000.

#### Determining the optimal number of latent factors

The optimal number of latent factors (*k*) was determined using `cross_validate_nmf()` from the *singlet* package (v0.99.6)^161^. Candidate values of *k* tested include 5, 10, 20, 30, 40, 50, 75, 100, and 125. For each rank three independent replicates were examined, and in each replicate a random subset of 20% of matrix entries was withheld during training and used to compute held-out test error. Models were fit using an iterative coordinate descent framework, with L1-sparsity regularization on *W* of 0.1, while *H* was not regularized. A convergence tolerance of 1×10^−3^ and a maximum of 100 iterations were used for each replicate. Model performance for each *k* was evaluated by averaging the held-out test error across replicates. The final rank (*k* = 66) was selected as the smallest value beyond which additional factors provided minimal improvement in test error while maintaining stability across replicates.

#### Inferring snRNA-seq derived factors and projection to SRT data

Using the optimal rank (*k* = 66), the final NMF model was fitted on log-normalized snRNA-seq data using the *RcppML* package (v0.5.6)^161^. The `nmf()` function was run with `k=66` factors, L1 regularization parameter of 0.1 applied to *W* (`L1=0.1`), non-negativity constraints on *W* and *H* (`nonneg=TRUE`), convergence tolerance of 1×10^−6^ (`tol`), and a maximum of 1,000 iterations (`maxit`). A fixed random seed (`seed=1135`) was used to ensure reproducibility. Zero values in the input matrix were not treated as missing during training (`mask_zeros=FALSE`), and diagonalization was introduced to scale factors in *W* and *H* to sum to 1 (`diag=TRUE`).

After estimating the final NMF model, the genes x factors matrix *W* was used to project latent transcriptional programs onto the SRT data. Program activities for each Visium spot were computed using the *projectR* package (v1.19.2)^162^, which projects the log-normalized SRT expression matrix (*A*’) onto the learned *W* matrix, generating a factors x spots activity matrix (*H*’) as shown in Eqn. 4.

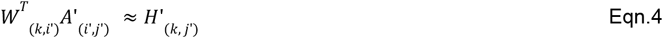

To facilitate comparability across spots, factor projections were normalized such that values for each spot summed to 1. The resulting projection matrix, [*nmf*1, *nmf*2, · · ·, *nmf*66] were appended to the `colData()` of the *SpatialExperiment*^135^ object and used in downstream visualization and interpretation.

#### Selecting SpD-associated factors

NMF-derived latent factors and their projections were summarized by SpD by computing the proportion of SRT spots with nonzero factor weights and the *z*-scored mean factor weight within each group. Two factors (nmf26 and nmf45) were excluded due to correlations with donor sex (|*r* > 0. 3), and an additional eight factors were removed because they were detected in fewer than 200 SRT spots. Factors were considered SpD-associated when they exhibited > 20% prevalence and scaled average activity > 0.2. This led to the identification of seven factors associated with D1 islands, and 14 factors which were associated with MSN_1-3. After excluding factors which mapped to multiple SpDs in a non-specific manner, we identified a set of MSN-specific factors including nmf3, nmf4, nmf7, nmf10, nmf25, nmf38, and nmf39. Additionally, we identified that nmf34, nmf35, and nmf44 mapped specifically to D1 islands.

#### Interpretation of NMF-derived factors

To functionally interpret the NMF-derived latent factors, we combined multiple complementary approaches and examined gene loadings, expression correlations, and performed pathway enrichment analyses. First, we identified the top-loaded genes in the *W* matrix (genes x factors) for each factor of interest, ranking genes by their contribution to the corresponding transcriptional program. Next, we computed Pearson correlations between factor scores (*H*) and log-normalized snRNA-seq expression values, using these correlations to rank genes and identify those most strongly associated with each factor within the snRNA-seq data. To complement this, we correlated factor projections in the spatial transcriptomics data (*H*’) with the SRT log-normalized expression matrix to identify genes spatially co-varying with factor activity across Visium spots.

Finally, gene set enrichment analysis (GSEA) was performed using the *fgsea* package (v1.28.0)^72^ with Reactome pathway annotations. For each factor, genes were ranked by their NMF weights (*W*), and `fgseaMultilevel()` was used to compute enrichment scores by iteratively walking through the ranked list and increasing a running-sum statistic when pathway genes were encountered and decreasing it otherwise. The multilevel sampling strategy employed by `fgseaMultilevel()` adaptively estimates extremely small *p*-values with improved accuracy compared to standard permutation-based GSEA. Normalized enrichment scores (NES) and adjusted p-values were calculated to determine pathway significance. Analyses were restricted to positively weighted genes (`scoreType=“pos”`) and limited to pathways containing between 15 and 500 genes.

### 4.17 Stratified linkage disequilibrium analysis to examine enrichment of trait heritability for cell types, spatial domains, and continuous factors

Stratified linkage disequilibrium score regression (s-LDSC)^49^ was applied to determine whether the genetic risk for a range of neuropsychiatric and behavioral traits was preferentially associated with genes linked to a specific cell type, spatial domain, or NMF-derived factors as detailed previously^146^. The analysis followed the LDSC guidelines (https://alkesgroup.broadinstitute.org/LDSCORE), using the baseline LD (v2.2)^163^ model to control for LD between SNPs in our focal annotation and SNPs annotated by any of the 97 baseline functional categories. HapMap Phase 3^164^ and European-ancestry 1000 Genomes Phase 3^165^ SNPs served as the reference panel. Enrichment was quantified from *z*-scores of per-SNP heritability, and statistical significance was assessed using Benjamini-Hochberg correction. We used an FDR threshold of < 0.1 for cell-type and SpD annotations and < 0.05 for NMF-derived factors.

#### Selection of genes linked to snRNA-seq cell types and SRT spatial domains

To generate annotations for cell types and SRT-derived spatial domains (SpDs), we constructed pseudobulk expression profiles by summing raw counts across all nuclei within each cell type or across all spots within each SpD. For snRNA-seq, nuclei labeled `Neuron_Ambig` were excluded prior to aggregation. Genes were filtered using `filterByExpr()`from the *edgeR*^147^ package to retain protein-coding genes with sufficient counts and duplicated genes were removed. Counts were normalized by counts per million (CPM) normalization, and relative expression for each gene was computed as its CPM divided by the total CPM within the corresponding cell type or SpD. Genes within the top 10% of relative expression were selected for cell type annotations, while the top 15% were selected for SpD annotations. Genomic coordinates for these genes were obtained from the hg19 reference annotation, and intervals were defined as 100 kb upstream of the transcription start site and 100 kb downstream of the transcription end site. Regions on chrX, chrY, and chrMT were excluded, and the resulting BED files were used to construct annotations, where SNPs falling within these intervals were assigned a value of 1 and 0 otherwise.

#### Selection of genes linked to NMF-derived factors

For NMF-based annotations, factor-specific gene sets were derived from the log-normalized snRNA-seq expression matrix and the final fitted NMF model. Across both the expression and factor scores matrix *(H)*, genes were filtered to only retain protein-coding genes. Further, any genes with duplicated gene names were excluded. For each factor-gene pair, the Pearson correlation was computed between the factor scores *(H)* and log-normalized gene expression across all nuclei to generate a genes x factors correlation matrix. The top 10% of genes ranked by correlation were selected as factor-specific genes. Genomic intervals were defined using hg19 coordinates, extending 100 kb upstream of the transcription start site and 100 kb downstream of the transcription end site, while excluding regions on chrX, chrY, and chrMT. BED files were created for each factor and the corresponding annotation was constructed by assigning SNPs within the defined intervals a value of 1 and 0 otherwise.

### 4.18 Analysis of trait-associated ligand-receptor (LR) analysis in snRNA-seq and SRT data

Building on *s-LDSC*^*49*^ analyses that linked nucleus accumbens (NAc) cell types, spatial domains, and NMF derived factors to trait heritability, we next examined cell-cell signaling within the NAc as a potential mechanism through which genetic risk is mediated. We (i) inferred interacting sender/receiver cell types from snRNA-seq data, (ii) prioritized ligand–receptor (LR) pairs in which at least one partner is a trait-risk gene, and (iii) integrated our findings with SRT data to localize putative sender/receiver pairs within tissue.

#### Inference of LR interactions in snRNA-seq data

A processed *SingleCellExperiment*^118^ object (**Materials and Methods: Identification of differentially expressed genes corresponding to underlying cell types in snRNA-seq data**) was used to infer cell-cell communication with LIANA (v0.1.14)^90^. First, nuclei labeled `Neuron_Ambig` were removed. Next, cell type labels defined in `CellType.Final` were used as the group identifiers for *LIANA*^90^ and gene identifiers were converted to symbols to match *OmniPath*^91^. Finally, *LIANA*^90^ was run with `liana_wrap()` using default settings (`methods: NATMI`^166^`, Connectome`^167^`, logFC, SCA`^168^`, CellPhoneDB`^169^`; resource: Consensus derived from OmniPath`^91^ ) to score candidate LR pairs for each ordered sender and receiver cell type combination. The ranks obtained across the different methods were then combined with `liana_aggregate()` using robust rank aggregation, yielding a consensus rank and corresponding significance used for prioritization.

#### Trait-informed prioritization

Risk gene sets for depression, anxiety, substance abuse, and schizophrenia were obtained from *the OpenTargets* platform (https://www.opentargets.org/)^92^. The reported evidence included `gwasCredibleSets`, `geneBurden`, `eva`, `genomicsEngland`, `gene2Phenotype`, `uniprotLiterature`, `uniprotVariants`, `orphanet`, `clingen`, stored as numeric scores for each gene-trait pair. “`No data`” entries were set to 0 and, for each trait, a per-gene maximum evidence score was computed across these metrics. Genes with a maximum evidence score > 0.1 were retained as trait risk genes.

To integrate ligand-receptor (LR) biology, we queried the *OmniPath* intercellular database^91^ and extracted a directed L-R network (`transmitters: category “ligand”; receivers: category “receptor”`) and gene symbols began with “HLA” were removed to avoid MHC artifacts. For each trait, we compiled all directed LR pairs in which at least one partner was a risk gene and intersected these sets with LIANA^90^ -inferred interactions by matching directed identities, producing a consensus list supported by both our snRNA-seq data and trait relevance.

#### Contact-dependent CNTN3/4/6–PTPRG signaling in SRT

Since *CNTN3/4/6*-*PTPRG* signaling were identified as trait relevant LR pairs, we filtered *LIANA*^90^ predictions to only retain highly confident interactions (aggregate rank ≤ 0.01) to examine the interacting cell types by visualizing the expression magnitude (`sca.LRscore`) and interaction specificity (`natmi.edge_specificity`). Additionally, CNTN3, CNTN4, and *CNTN6* are membrane-bound contactins that act as axon-associated cell-adhesion ligands, and *PTPRG* is a transmembrane receptor. Thus, their interaction is contact-dependent and should be most evident where ligand- and receptor-expressing cells are in close proximity. To localize these interactions in the tissue, SRT spots were classified using log-normalized expression: a spot “expressed” a gene if its value was > 0. Each spot was then assigned to one of four categories for a given LR pair: co-expressing (ligand and receptor present), ligand-only, receptor-only, or neither. For each SpD we computed the proportion of spots co-expressing the ligand and receptor, the ligand-only, and the receptor-only. Further, to identify which cell types co-localize in co-expressing spots *RCTD*^64^-derived weights (**Materials and Methods: Spot deconvolution with Robust Cell Type Deconvolution (*RCTD*)**) were used. Specifically, if *W* is a matrix of *RCTD*^64^*-*weights of dimensions spots *x* cell types, *and S* _*LR*_ is the set of spots that co-express ligand and receptor for a given LR pair, and *S*_*ctrl*_ is the complementary set of spots that do not co-express the ligand and receptor, we computed the cell type co-occurence for *S* and *S* as given by Eqn. 5 and Eqn. 6.

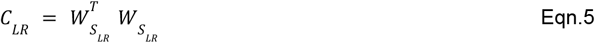

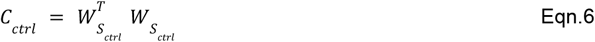

After zeroing the diagonal and upper diagonal entries, the lower triangle entries were normalized to sum to 1. The enrichment ratio (*R*) was then computed as shown in Eqn. 7.

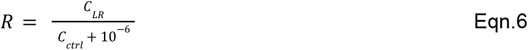

Entries of *R* > 1 indicates that the corresponding cell type pairs co-occured more often in spots which co-expressed the ligand and receptor in comparison to spots which do not. This analysis was performed independently for each LR pair.

#### SpD-specific localization of sender and receiver cell types for OPRM1 signaling

*We focused on OPRM1* (the μ-opioid receptor) signaling because it is central to reward processing, and both *PENK*-*OPRM1* and *PDYN*-*OPRM1* interactions were identified by applying *LIANA*^90^ to the snRNA-seq data, and independently prioritized as a trait-relevant LR pair for substance dependence (Materials and Methods: Trait-informed prioritization). Since *PENK* and *PDYN* are secreted neuropeptides, these interactions are consistent with short-range paracrine signaling that may span adjacent spatial domains (SpDs). Thus, to determine which SpDs are involved in *PENK*-*OPRM1* and *PDYN*-*OPRM1* signaling, we first examined the expression of *PENK, PDYN*, and *OPRM1* across the eight identified SpDs. Next, we visualized the contribution of *LIANA*^*90*^ predicted sender and receiver cell types across SRT spots after filtering to high-confidence interactions (aggregate rank ≤ 0.01). Specifically, we compared the *RCTD*^*64*^*-*derived weights of these cell types across SpDs (**Materials and Methods: Spot deconvolution with Robust Cell Type Deconvolution (RCTD)**). Since DRD1_MSN_A was the sole sender population for PDYN, and DRD2_MSN_A was the predominant sender of PENK signaling, we examined *RCTD*^64^*-*derived weights corresponding to these cell types. Additionally, we visualized *RCTD*^*64*^*-*derived weights corresponding to the two primary receiver populations DRD1_MSN_B and DRD1_MSN_D.

### 4.19 Gene set enrichment of differentially expressed genes in response to morphine and cocaine

To map genes that exhibit drug induced transcriptional differences onto SpD-specific differentially expressed genes (SpD-DEGs), we integrated rodent NAc snRNA-seq case-control datasets and asked whether genes responsive to morphine or cocaine exposure are over-represented among SpD-DEGs. We used two external datasets which profiled the transcriptional responses across acute and chronic (volitional) exposure paradigms in the rodent NAc in response to morphine^27^ and cocaine^26^. Both studies reported transcriptional differences between cells derived from morphine/cocaine-exposed subjects and paired controls across multiple annotated cell types. Thus, we obtained gene lists indexed by cell type, drug, and exposure paradigm.

Within each study, exposure paradigm, and cell type, we split genes into upregulated and downregulated gene sets based on the sign of the reported logFC, yielding one list per cell type, drug, exposure duration, and direction of transcriptional change. Rat gene symbols were mapped to human gene symbols as described previously using the MGI homology reference (`HOM_AllOrganism.rpt`), and the human gene symbols were converted to Ensembl IDs. Because the chronic cocaine condition contained too few DEGs, we did not stratify the cocaine DEG results across exposure paradigms, and instead used the cocaine DEG results corresponding to both the acute and chronic data. After applying a filter requiring each gene list to have 25 or more genes, to ensure sufficient power for over-representation analysis, we found that only a single cell type (Drd1_MSN_A) contained more than 25 DEGs in Phillips et al^26^. Hence, for the cocaine data, we refer to genes as either “cocaine upregulated” or “cocaine downregulated”, while implicitly noting that these DEGs were derived from the Drd1_MSN_A cell type across both exposure paradigms.

Over-representation analysis was performed using `gene_set_enrichment()` from *spatialLIBD*^58^. Briefly, this function takes an input list of genes and previously computed enrichment statistics derived from pseudobulk aggregation and one-vs-all enrichment testing (**Materials and Methods: Pseudobulk differential expression analysis**). These inputs are used to generate a 2 x 2 contingency matrix summarizing the overlap between the input gene list and SpD-DEGs, which were defined as genes with FDR < 0.05 and *t*-statistic > 0. This contingency matrix was used to perform a one-sided Fisher’s test, and the resulting odds ratio (OR) and *p*-value were reported.

### 4.20 Examining continuous transcriptional programs in response to morphine and cocaine in SRT data

To characterize drug responsive transcriptional programs that can occur across multiple cell types, we applied NMF to rodent NAc snRNA-seq case-control datasets and derived latent factors that capture coordinated gene expression changes induced by morphine or cocaine exposure. We then projected the inferred factors to SRT data to spatially localize program activity, providing a complementary analysis to our earlier test of over-representation of morphine/cocaine DEGs in SRT-derived SpDs.

#### Input data and feature alignment

The input human NAc SRT data was processed as described in Materials and Methods: SRT data processing and quality control. We inferred latent factors to capture drug-responsive transcriptional changes in three case-control datasets: acute morphine and chronic morphine self-administration^27^, and acute cocaine^26^, each with matched saline controls. Prior to applying NMF, we first focused our analysis on MSNs. For cocaine we retained nuclei annotated to one of the following cell types including “Drd1-MSN-1”, “Drd1-MSN-2”, “Drd2-MSN-1”, “Drd2-MSN-2”, “Drd3-MSN”, and “Grm8-MSN”. For morphine we retained nuclei annotated as either “Drd1 MSN1”, “Drd1 MSN2”, “Drd1 MSN3”,”Drd2 MSN1”, “Drd2 MSN2”, “Drd2 MSN3”, “Drd2 MSN4”, “Grm8 MSN”, “MSN1”, “MSN2”, or “MSN3”. Rat gene symbols were mapped to human gene symbols as described previously using the MGI homology reference (`HOM_AllOrganism.rpt`) and only unique one-to-one mappings were retained. The mapped human genes were then intersected with genes detected in the SRT data, and all downstream analyses were restricted to this shared gene set. After subsetting both the rodent snRNA-seq and SRT counts to this shared gene set, the expression data were log-normalized with *Seurat*^141–145^ `LogNormalize()` (v5.0.1, scale factor 10,000).

#### Inferring factors and testing drug association

For each rodent case-control cohort (acute morphine, chronic morphine self administration, and acute cocaine), we inferred latent factors with NMF implemented in *RcppML* (v0.5.6)^161^ on the MSN-restricted log-normalized expression matrix as previously described in **Materials and Methods: Inferring snRNA-seq derived factors and projection to SRT data**. Specifically, we ran `nmf()` after excluding genes with no detected expression across all selected nuclei, with the number of latent factors (`k`) set to 30. All other functional parameters were set as before (`L1=0.1, mask_zeros=FALSE, nonneg=TRUE, diag=TRUE, tol=1e-6, maxit=1000, seed=1135)`. The NMF results yielded a gene loadings matrix *W* _*r*_ and a factor score matrix *H*_*r*_ across all nuclei.

To identify NMF factors which were associated with drug exposure, we compared the factor scores between drug exposed nuclei and their matched saline controls within each MSN subtype, using the Wilcoxon rank sum test and Benjamini-Hochberg multiple testing correction. Factors were identified as drug responsive when the corresponding Wilcoxon effect-size > 0.3 and FDR < 0.05. We identified two factors associated with acute morphine including nmf3 and nmf12. For volitional morphine, we identified three drug associated factors including nmf3, nmf12, and nmf23. Finally, nmf5 and nmf28 were found to be associated with acute cocaine.

#### Projection of rodent-derived factors to SRT data

We projected the rodent snRNA-seq derived gene loadings (*W*_*r*_ ) to human NAc SRT data using a linear model as previously detailed in Materials and Methods: **Inferring snRNA-seq derived factors and projection to SRT data**. Briefly, for each case-control cohort, we read the NMF results from *RcppML*^161^, and extracted the gene loadings matrix *W*_*r*_ where the rows correspond to rodent genes and the columns to individual factors. First, the rows were mapped to human genes as previously described. The rows of *W*_*r*_ were then re-ordered to match the gene names in the rows of the log-normalized SRT expression matrix. Finally, factor projections across SRT spots was computed using the *projectR* package (v1.19.2)^162^, to obtain 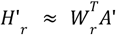, a factors x spots matrix. For comparability across spots, each column of 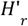 was normalized to sum to 1. The resulting projections [*nmf*1, *nmf*2, …, *nmf*30] were appended to the `colData()` of the *SpatialExperiment*^*135*^ object for downstream visualization and interpretation.

We visualized per-spot projection weights, and then summarized the projections by SpD as detailed in **Materials and Methods: Interpretation of NMF-derived factors** by computing (1) the proportion of spots with non-zero projection weights, and (2) z-scored mean factor weight. Finally, we correlated SRT gene expression with each projected factor and examined the top loaded rodent genes for the same factor to functionally characterize morphine and cocaine responsive programs.

## 4.21 Data and code availability

Raw data generated as a part of this study have been deposited on the Gene Expression Omnibus (GEO) with accession numbers GSE307586 and GSE307587. Processed data and all code used to perform the analyses in this paper is publicly available through Github at https://github.com/LieberInstitute/spatial_NAc, and have been permanently archived on Zenodo (doi: https://doi.org/10.5281/zenodo.17089020). Large processed data files which exceeded Github size limits are available through Globus (https://research.libd.org/globus/) under jhpce#spatial_NAc. Unless noted otherwise, all analyses were performed using R (v4.3.2)^170^, Bioconductor (v3.18)^171^, and Python (v3.12.11). ImageJ (v2.9.0)^132^ was used for image alignment. We also provide two interactive websites to explore the findings presented in this work which can be accessed at https://research.libd.org/spatial_NAc/#interactive-websites under Visium for the spatial transcriptomics data, and iSEE for the snRNA-seq data. The Visium application uses *spatialLIBD*^58^ to interactively visualize gene expression, RCTD^64^ cell type deconvolution scores, and *MERINGUE*^59^ and NMF factor projection scores overlaid on H&E images for all ten donors. The *iSEE*^*172*^ application visualizes annotated cell types in user-defined lower dimensional embeddings, and gene expression across cell types.

## 4.22 Acknowledgements, Funding, Authorship Contributions

## Acknowledgements

We thank the LIBD neuropathology team, particularly James Tooke and Amy Deep-Soboslay, for curation of the brain samples and assistance with tissue dissections. *We thank Dr. Hao* Zhang at the Johns Hopkins Cell Sorting Core facility and the Johns Hopkins University Single Cell and Transcriptomics Sequencing Core. We thank Dr. Melissa Grant-Peters for consultation on ligand receptor analyses. We are grateful to Dr. Brian Herb and Dr Seth Ament, University of Maryland School of Medicine for their advice and feedback on transfer learning and factorization approaches. We thank Dr. Benjamin Reiner, University of Pennsylvania Perelman School of Medicine for sharing snRNA-seq data from experiments involving acute and volitional morphine exposure in rodents. We thank the staff and physicians at the brain donation sites, and the generosity of the brain donors and their families, without whom this work would not be possible. Finally, we thank the families of Connie and Steve Lieber and Milton and Tamar Maltz for their generous support. Portions of some figures were created with Biorender.

## Author Contributions

Conceptualization: KM, SCH, KRM

Methodology: PR, SVB, NJE, TMH, LCT

Software: NJE, RAM, LCT

Validation: SVB, YD, IDR

Formal Analysis: PR, RAP, NJE, SH

Investigation: SVB, MRV, YD, IDR, KDM, SCP

Resources: JEK, TMH

Data Curation: PR, NJE, RAM, HRD, MT, SCP, LCT

Writing – original draft: PR, SVB, RAP, KRM

Writing – review & editing: PR, SVB, RAP, NJE, SCP, TMH, LCT, AB, KM, SCH, KRM Visualization: PR, SVB, RAP, NJE

Supervision: LCT, AB, KM, SCH, KRM, SH

Funding acquisition: KM, SCH, KRM

Project Administration: KM, SCH, KRM

