## Supplementary Materials for "Spatiomolecular mapping reveals anatomical organization of heterogeneous cell types in the human nucleus accumbens"

### Supplementary Figures

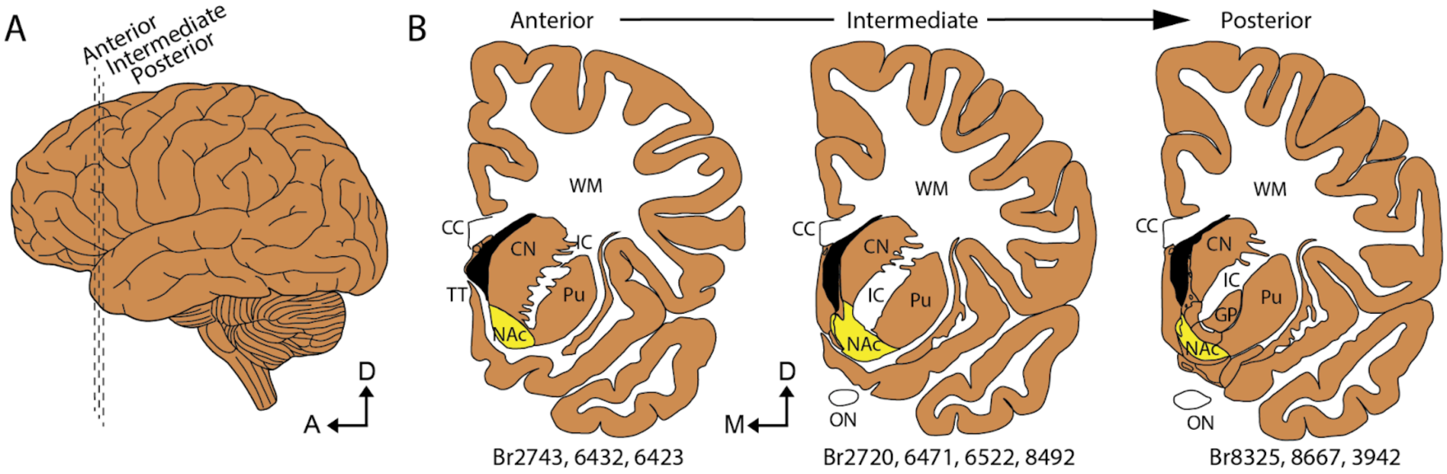


**Supplementary Figure. 1. Schematic representation of three anatomical levels sampled across the NAc.** (**A)** Schematic of the lateral side of the brain with three levels (anterior, intermediate, and posterior) of coronal sections represented by dashed lines. A - anterior; D - dorsal. (**B)** Three schematics of hemisected coronal brain slabs at the anterior, intermediate, and posterior extent of the NAc. CC - corpus callosum; CN - caudate nucleus; D - dorsal; GP - globus pallidus; IC - internal capsule; M - medial; NAc - nucleus accumbens; ON - optic nerve; Pu - putamen; TT - taenia tecta; WM - white matter. Brain (Br) numbers for donors at each level are listed below.

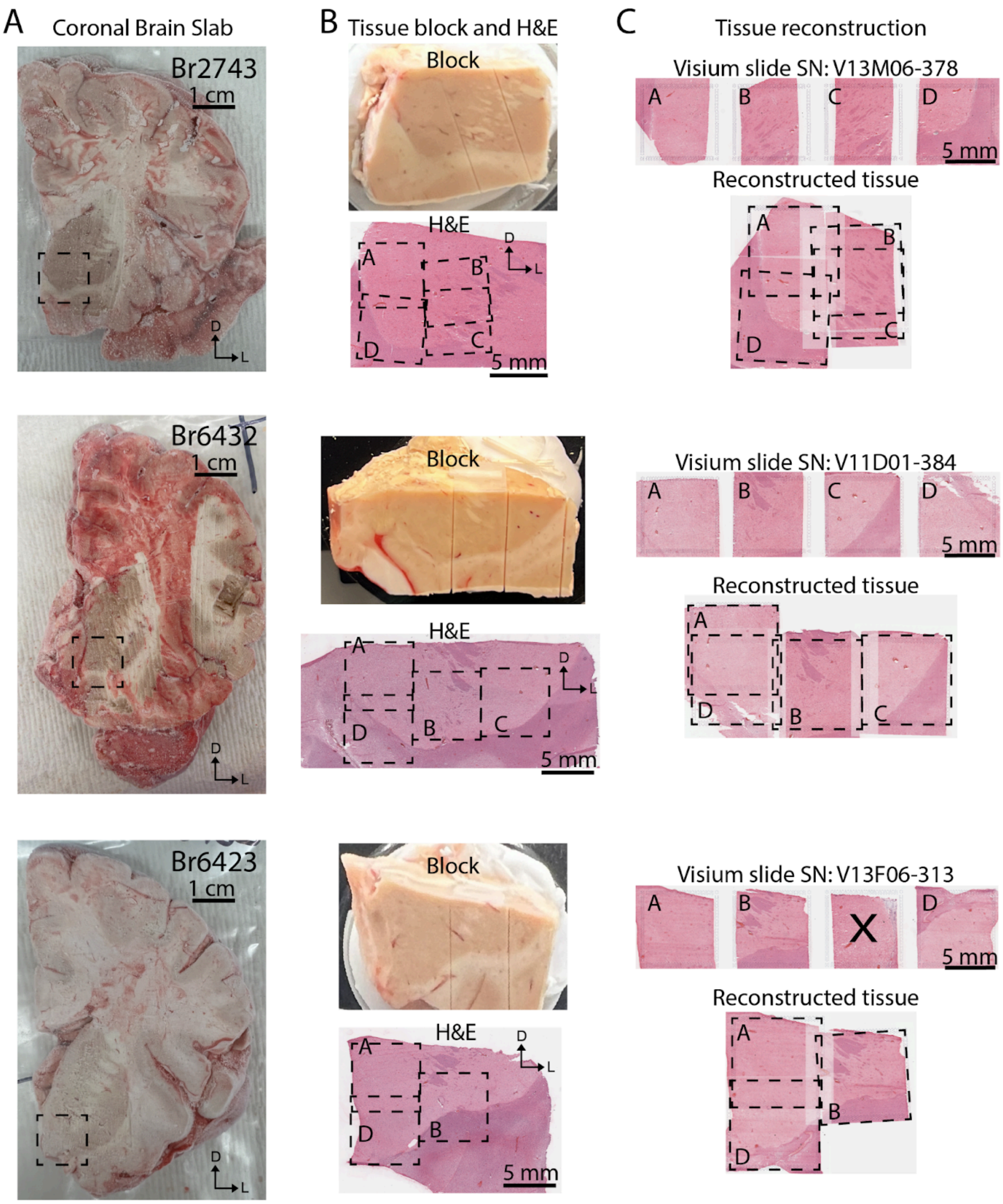


**Supplementary Figure. 2. Postmortem human brain dissections for the anterior NAc.** **(A)** Fresh frozen coronal postmortem human brain slabs from n=3 neurotypical control donors containing the anterior NAc. Dashed boxes indicate the location of the dissected blocks in **(B)**. Neuroanatomical orientation is indicated by arrows: D - dorsal; L - lateral. **(B)** Dissected scored tissue blocks and H&E quality control images for each anterior NAc donor. Dashed boxes indicate the approximate area placed on 3 to 4 Visium capture areas in **(C)**. Letters A-D represent each Visium capture area on a slide. Neuroanatomical orientation is indicated by arrows: D - dorsal; L - lateral. **(C)** H&E stained sections from each donor, as placed on Visium slides with slide number annotated (top). Letters A-D designate each Visium capture area. Reconstructed anterior NAc via stitching the tissue from each individual Visium capture area (bottom). X indicates discarded tissue from unrelated experiments.

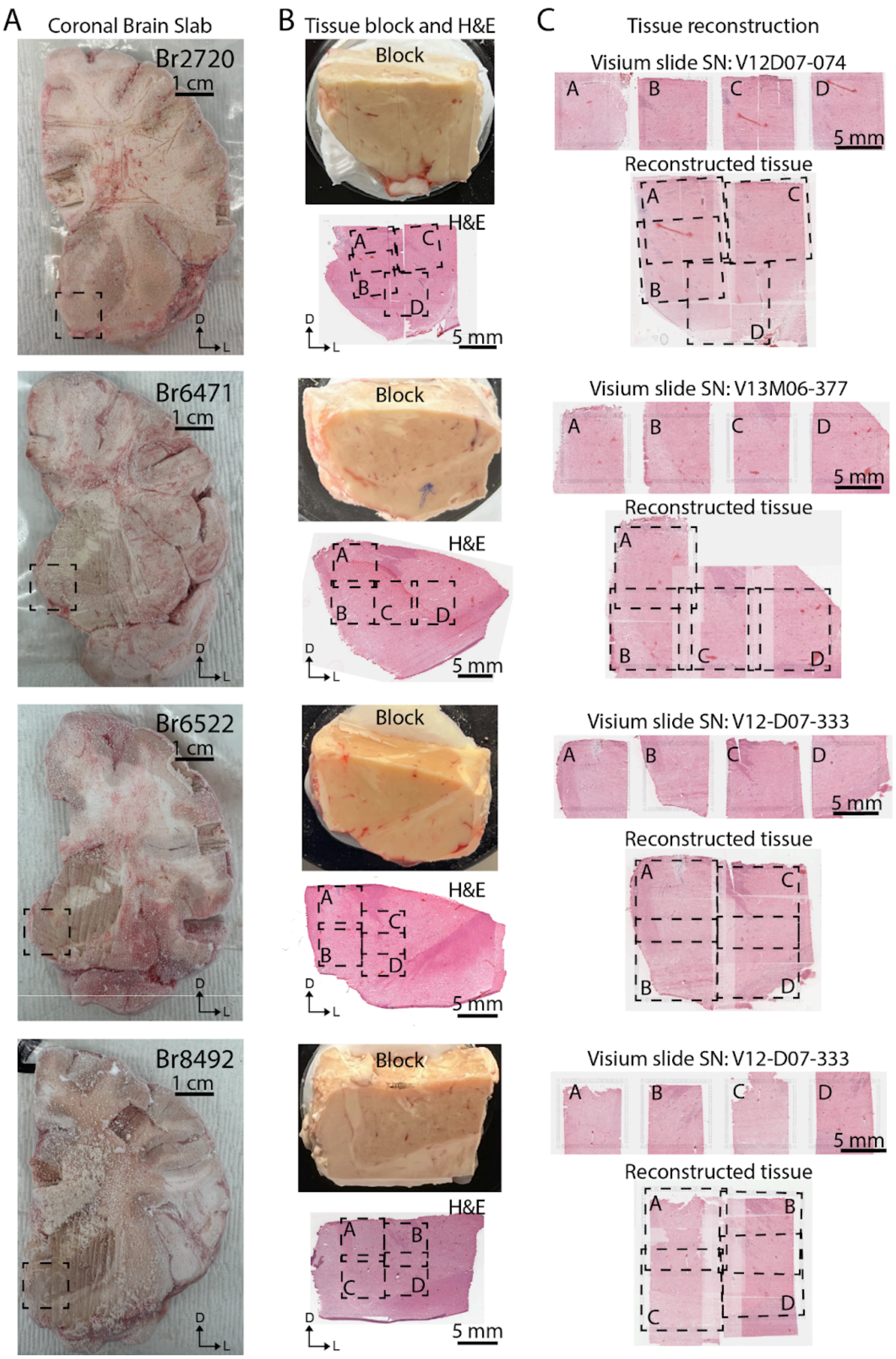


**Supplementary Figure. 3. Postmortem human brain dissections for the intermediate NAc. (A)** Fresh frozen coronal postmortem human brain slabs from n=4 neurotypical control donors containing the intermediate NAc. Dashed boxes indicate the location of the dissected blocks in **(B)**. Neuroanatomical orientation is indicated by arrows: D - dorsal; L - lateral. **(B)** Dissected scored tissue blocks and H&E quality control images for each intermediate NAc donor. Dashed boxes indicate the approximate area placed on 4 Visium capture areas in **(C)**. Letters A-D represent each Visium capture area on a slide. Neuroanatomical orientation is indicated by arrows: D - dorsal; L - lateral. **(C)** H&E stained sections from each donor as placed on Visium slides with slide number annotated (top). Letters A-D designate each Visium array. Reconstructed intermediate NAc via stitching the tissue from each individual Visium capture area (bottom).

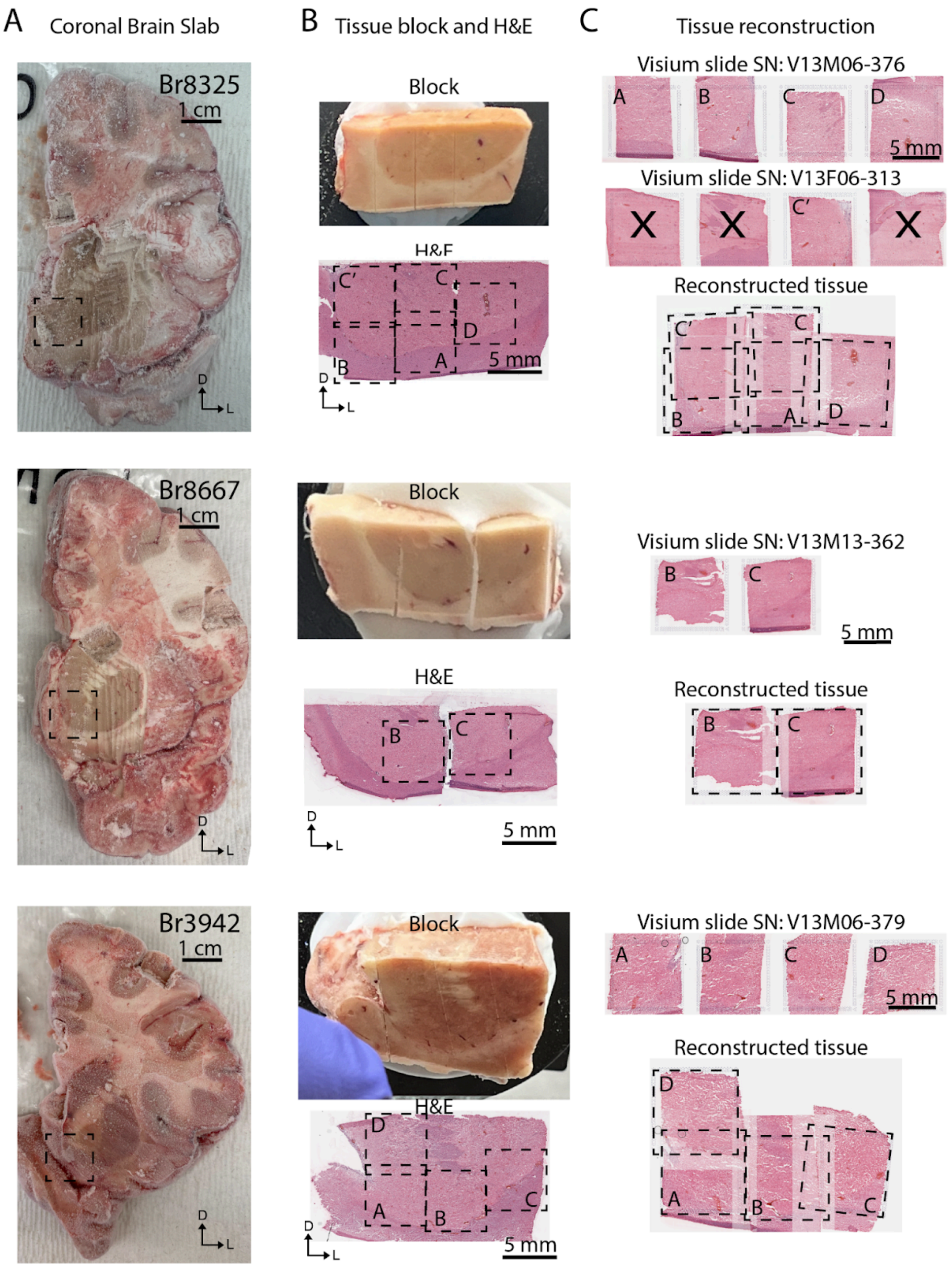


**Supplementary Figure. 4. Postmortem human brain dissections for the posterior NAc.**
**(A)** Fresh frozen coronal postmortem human brain slabs from n=3 neurotypical control donors containing the posterior NAc. Dashed boxes indicate the location of the dissected blocks in **(B)**. Neuroanatomical orientation is indicated by arrows: D - dorsal; L - lateral. **(B)** Dissected scored tissue blocks and H&E quality control images for each posterior NAc donor. Dashed boxes indicate the approximate area placed on 2 to 5 Visium capture areas in **(C)**. Letters A-D represent each Visium capture area on a slide. Neuroanatomical orientation is indicated by arrows: D - dorsal; L - lateral. **(C)** H&E stained sections from each donor as placed on Visium slides with slide number annotated (top). Letters A-D designate each Visium capture area. Reconstructed posterior NAc via stitching the tissue from each individual Visium capture area (bottom). X indicates discarded tissue from unrelated experiments.

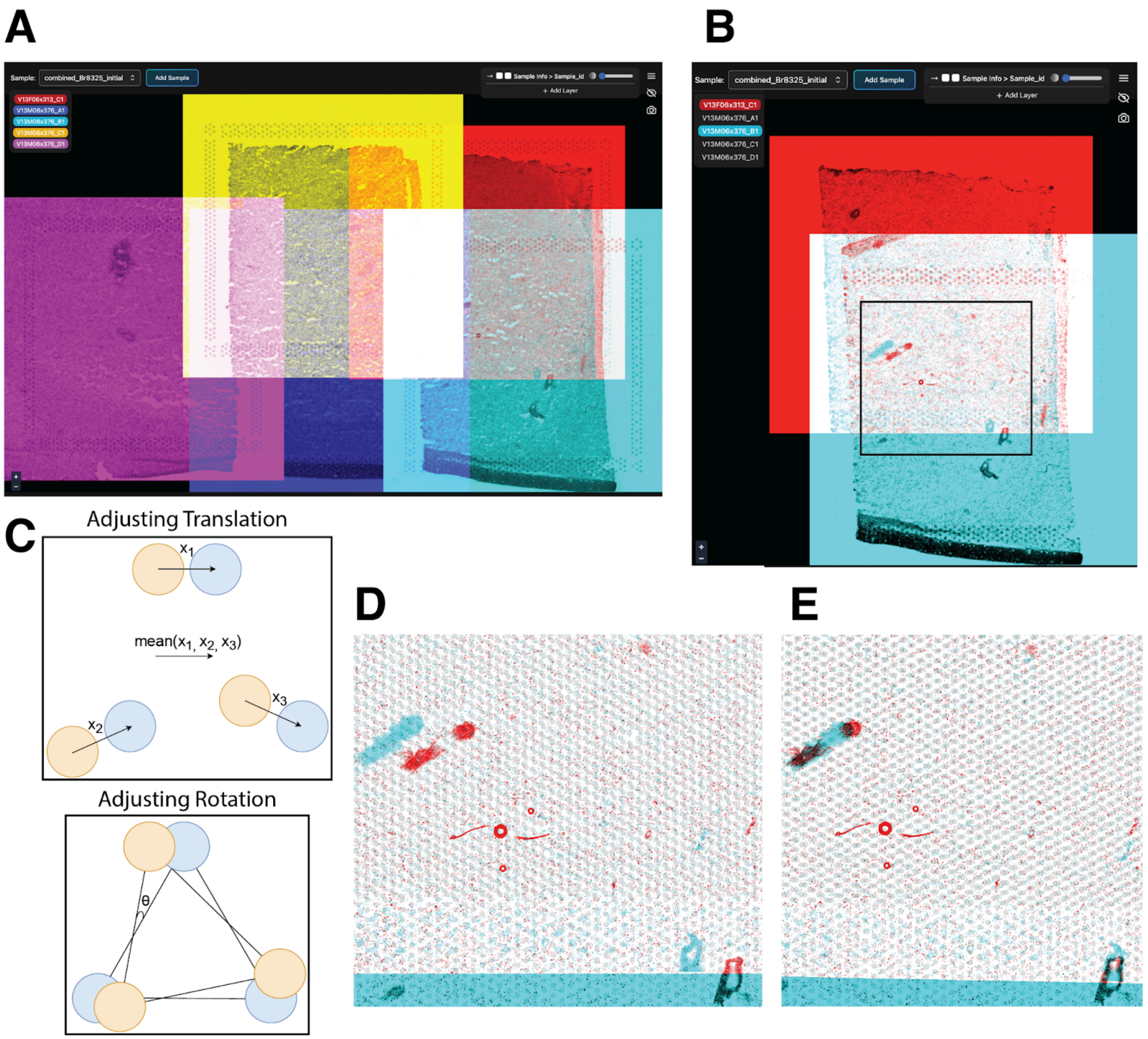


**Supplementary Figure. 5. Using *Samui Browser* and *Python* to refine image transformations from *ImageJ*.** (**A**) Each of Br8325's four capture areas were stitched following transformations in *ImageJ*, and placed in separate image channels to easily toggle individual capture areas on and off with *Samui Browser*. (**B**) The channels for capture areas V13F06-313_C1 and V13M06-376_B1 were toggled on, and a small region in their overlap is outlined in black, shown more closely in (**D**) and (**E**). (**C**) Example graphic illustrating how an optimal translation and rotation can be applied to three pairs of shared features, similarly to those depicted in **(D)**, from two overlapping capture areas (depicted in orange and blue, **Methods: Image stitching**). (**D**) Image features, such as blood vessels, are depicted from a small region of overlap between capture areas V13F06-313_C1 (in blue) and V13M06-376_B1 (in red), using *ImageJ* transformation estimates. In transparent gray, spots from both capture areas show the actual scale of image features, which can be seen to be out of alignment. (**E**) The same region of overlap between V13F06-313_C1 and V13M06-376_B1 is shown after an optimal adjustment to the initial *ImageJ* transformations is applied (**Methods: Image stitching**); image features align more closely.

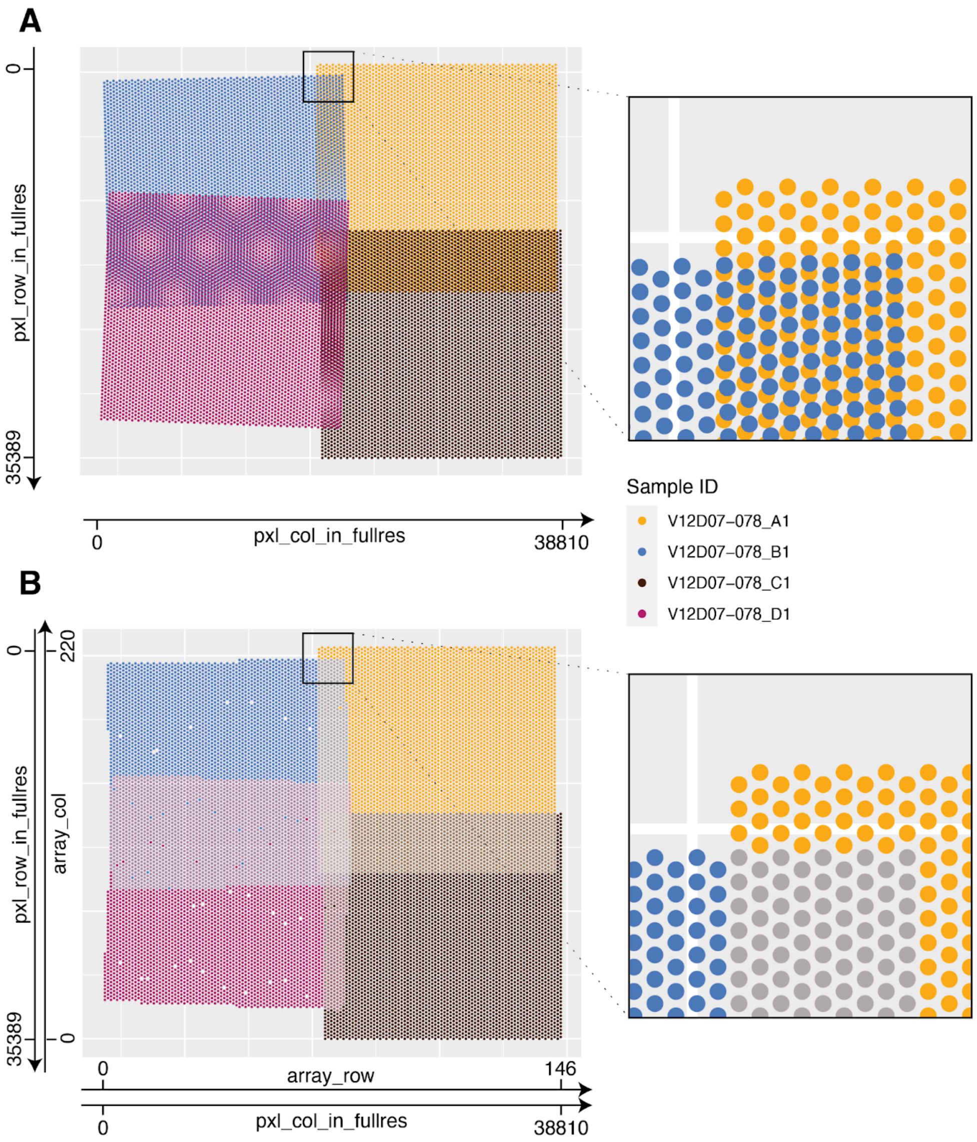


**Supplementary Figure. 6. Redefining array coordinates for stitched images**. (**A**) The exact positions of spots for Br8492 after stitching its capture areas with the *Samui Browser-Python* workflow (**Materials and Methods: Image stitching**). Capture areas V12D07-078_A1 and V12D07-078_B1 overlapped, but in general, their spots did not precisely align due to relative rotations and translations (shown on the right). As a result, `array_row` and `array_col` were poorly defined after merging and were omitted from the X and Y axes, respectively. (**B**) The adjusted spot positions for Br8492 after fitting to an artificial array encompassing all four capture areas (**Materials and Methods: Defining array coordinates for merged images**). Spots from (**A**) were mapped to the nearest artificial spot location, constrained both to match the hexagonal structure of a typical Visium capture area and match the standard inter-spot distance of 100 microns. This potentially resulted in two or more spots from (**A**) being assigned the same new array coordinates (depicted here by adjusting transparency on the left and as gray spots on the right). Notably, as is standard with *SpaceRanger* array coordinates specifications, `array_row` increases with the value of `pxl_col_in_fullres`, while `array_col` decreases with increasing `pxl_row_in_fullres`.

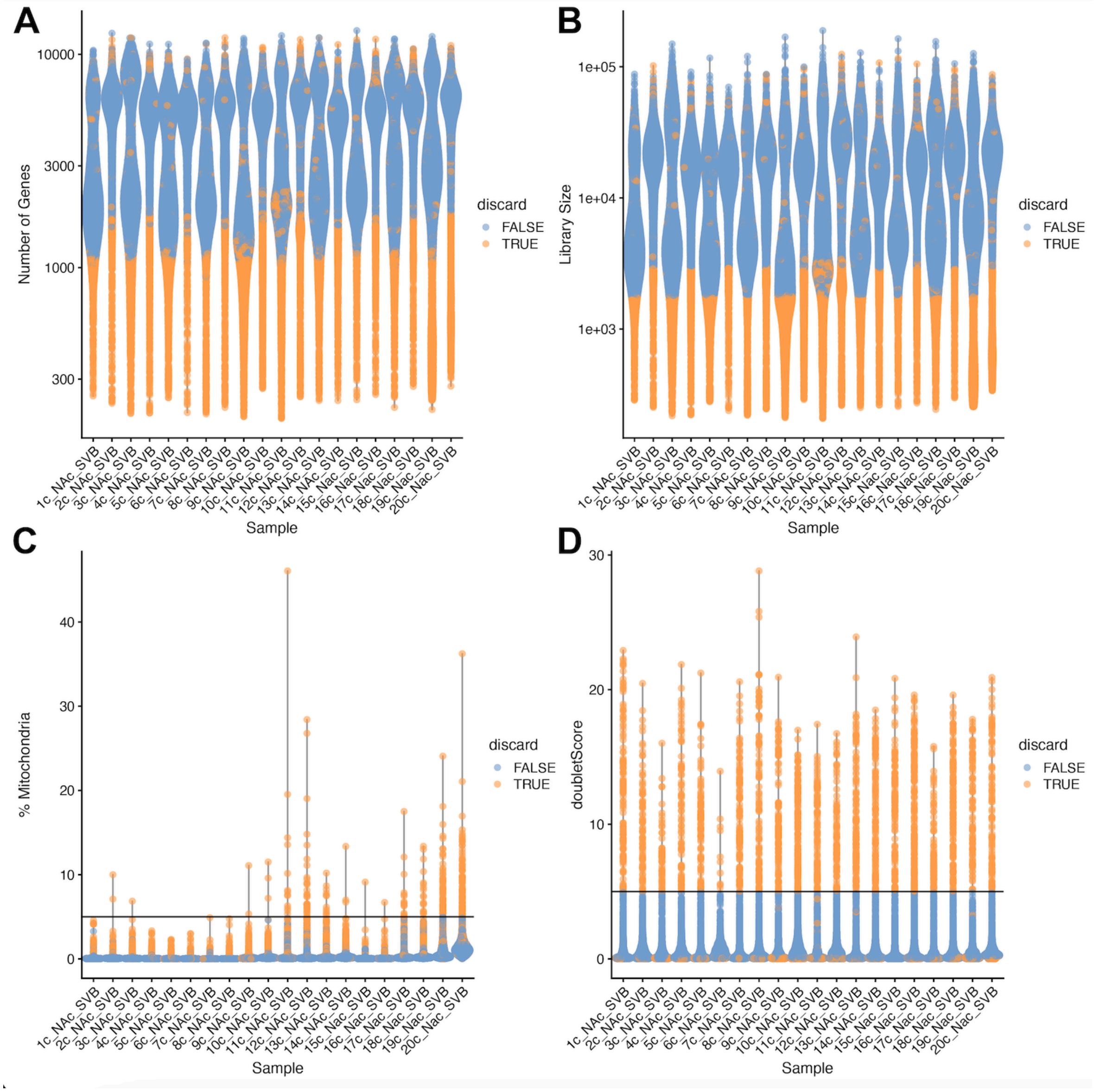


**Supplementary Figure 7. Quality control for snRNA-seq samples.** Quality control was performed using a combination of fixed and adaptive thresholds. Violin plots for (**A**) number of genes (**B**) library size (**C**) Percentage of reads mapping to mitochondrial genome and (**D**) `doubletScore` by snRNA-seq sample. Points are colored according to whether they were retained or discarded based on set criteria (see **Methods**).

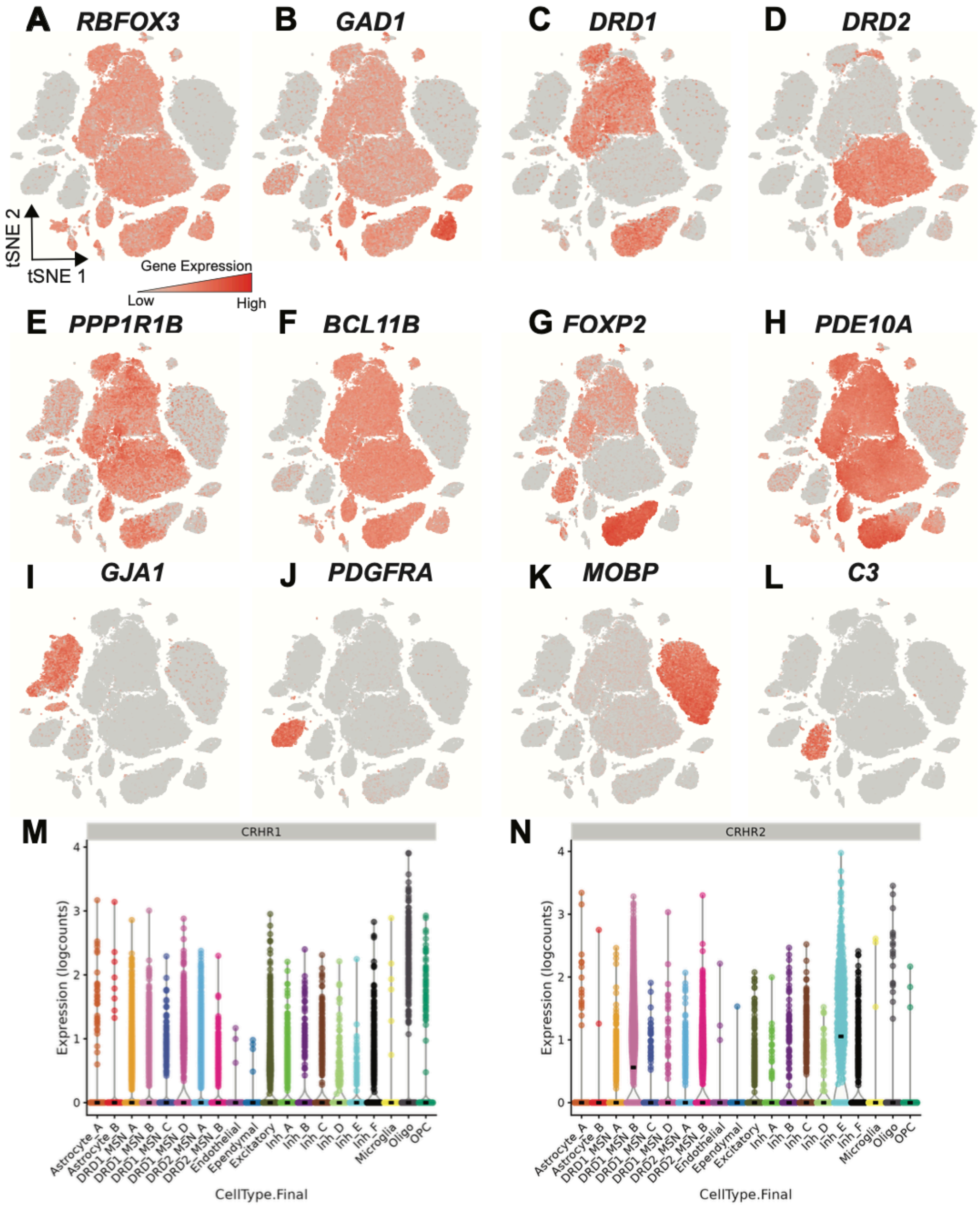


**Supplementary Figure. 8. Distribution of selected genes across cell types identified in the adult human NAc.** Feature plots colored by expression of neuronal marker genes, (**A**) *RBFOX3* and (**B**) *GAD1*. Medium spiny neurons (MSNs) constitute a large proportion of neurons within the NAc and are marked by expression of (**C**) *DRD1* (**D**) *DRD2* (**E**) *PPP1R1B* (**F**) *FOXP2* (**G**) *BCL11B*, and (**H**) *PDE10A*. In addition to neuronal subtypes, the human NAc contains non-neuronal cell types marked by expression of (**I**) *MOBP*, (**J**) *GJA1*, (**K**) *PDGFRA*, and (**L**) *C3*. Violin plots showing distribution of expression for (**M**) *CRHR1* and (**N**) *CRHR2*.

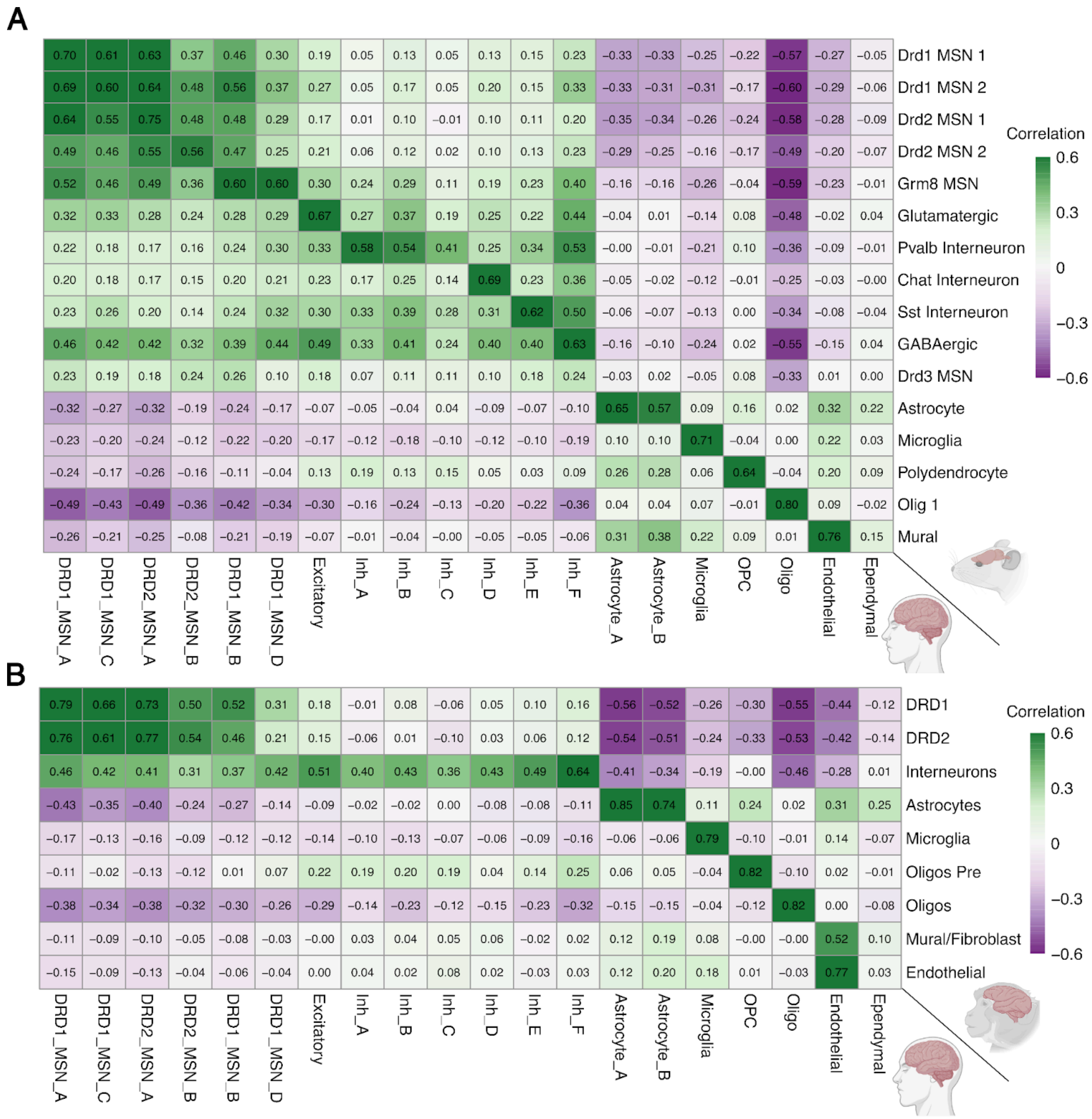


**Supplementary Figure. 9. Cross-species registration of neuronal and glial cell types in the human NAc. (A)** Heatmap of Pearson correlations between per-gene *t*-statistics from one-vs-all tests computed for each reported human cell type (rows) and rodent reference [26] cell type (columns). Correlation was evaluated over the union of the top 250 DEGs for each human cell type which uniquely mapped to rodent orthologs. **(B)** Same as **(A)** but comparing human NAc cell types (rows) to a non-human primate (NHP) reference [24] cell types (columns), evaluated over the union of the top 250 DEGs for each human cell type which uniquely mapped to macaque orthologs. Tiles show correlation coefficients (green = positive; purple = negative).

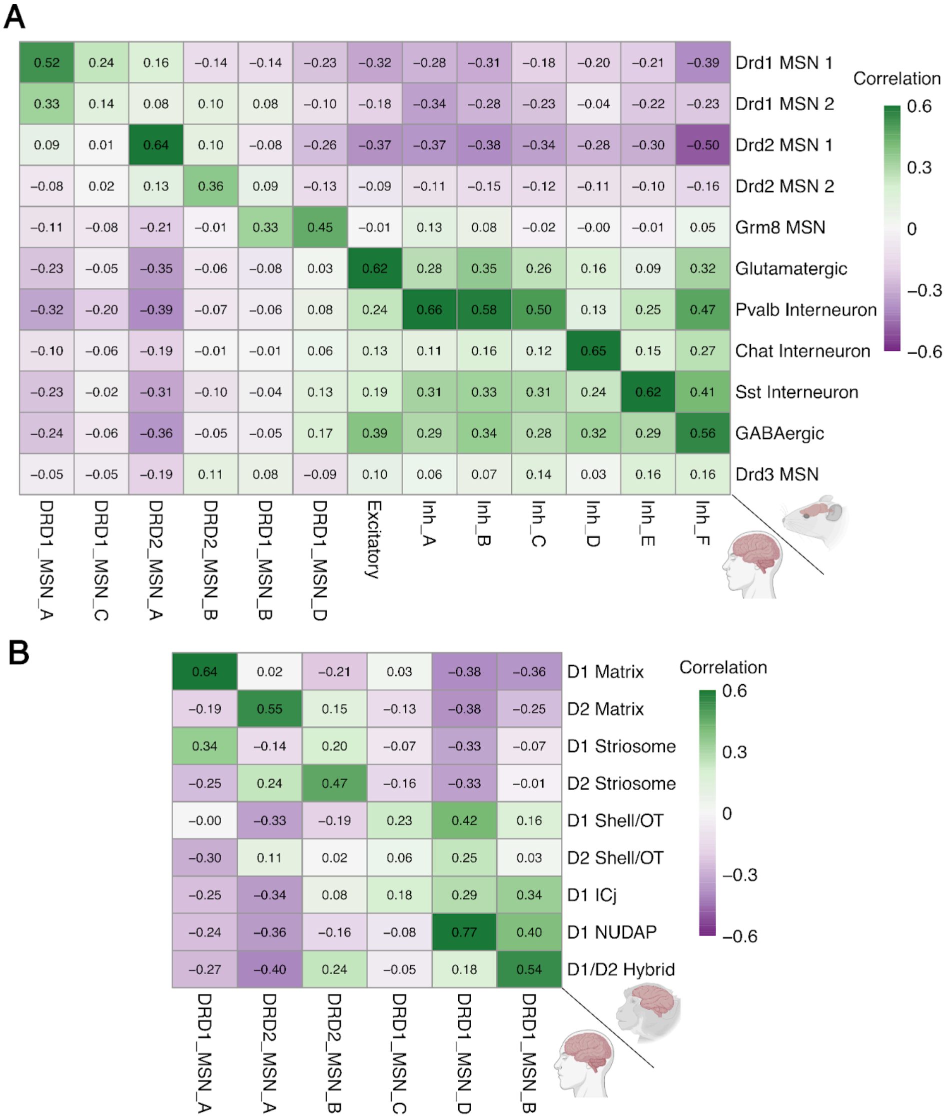


**Supplementary Figure. 10. Cross-species registration of neuronal cell types. (A)** Heatmap of Pearson correlations between per-gene *t*-statistics from one-vs-all tests computed for each neuronal human cell type (rows) and rodent reference [26] cell type (columns). Correlation was evaluated over the union of the top 250 DEGs per human neuronal cell type mapped to 1:1 human-rodent orthologs.**(B)** As in **(A)**, but comparing human MSN subtypes (rows) to the non-human primate reference (MSN-only, columns) [24], using 1:1 human-macaque orthologs.Tiles show correlation coefficients (green = positive; purple = negative).

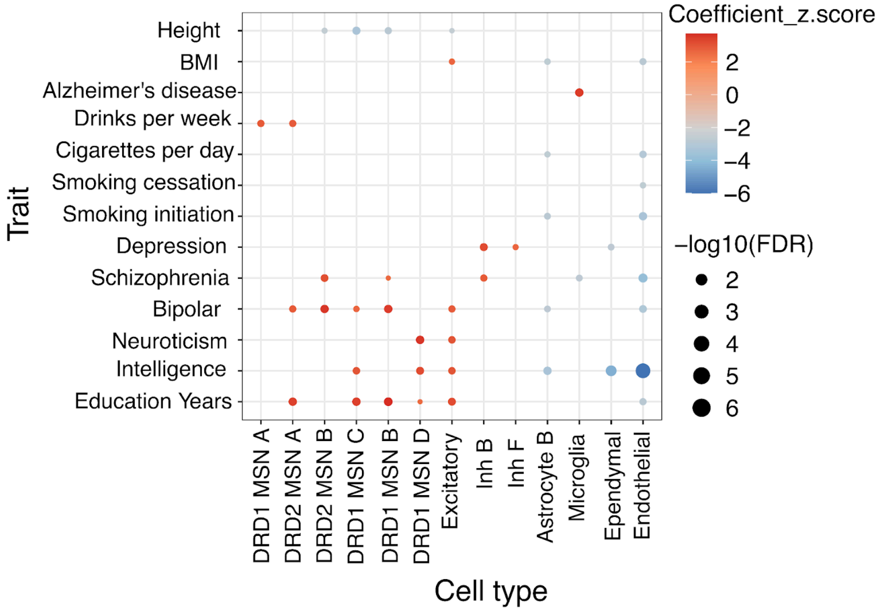


**Supplementary Figure. 11. Heritability enrichment of genes corresponding to NAc cell types.** Dot plot summarizing significant (FDR < 0.1) stratified LD score regression (s-LDSC) results across neuropsychiatric and behavioral traits for gene sets derived from human NAc snRNA-seq cell types. Dot color encodes the regression coefficient z-score (red = enrichment; blue = depletion) and dot size encodes $-\log_{10}$(FDR). Notable enrichments include Alzheimer's disease in microglia, drinks per week in DRD1_MSN_A and DRD2_MSN_A, and depression in inhibitory neuron populations Inh_F and Inh_B.

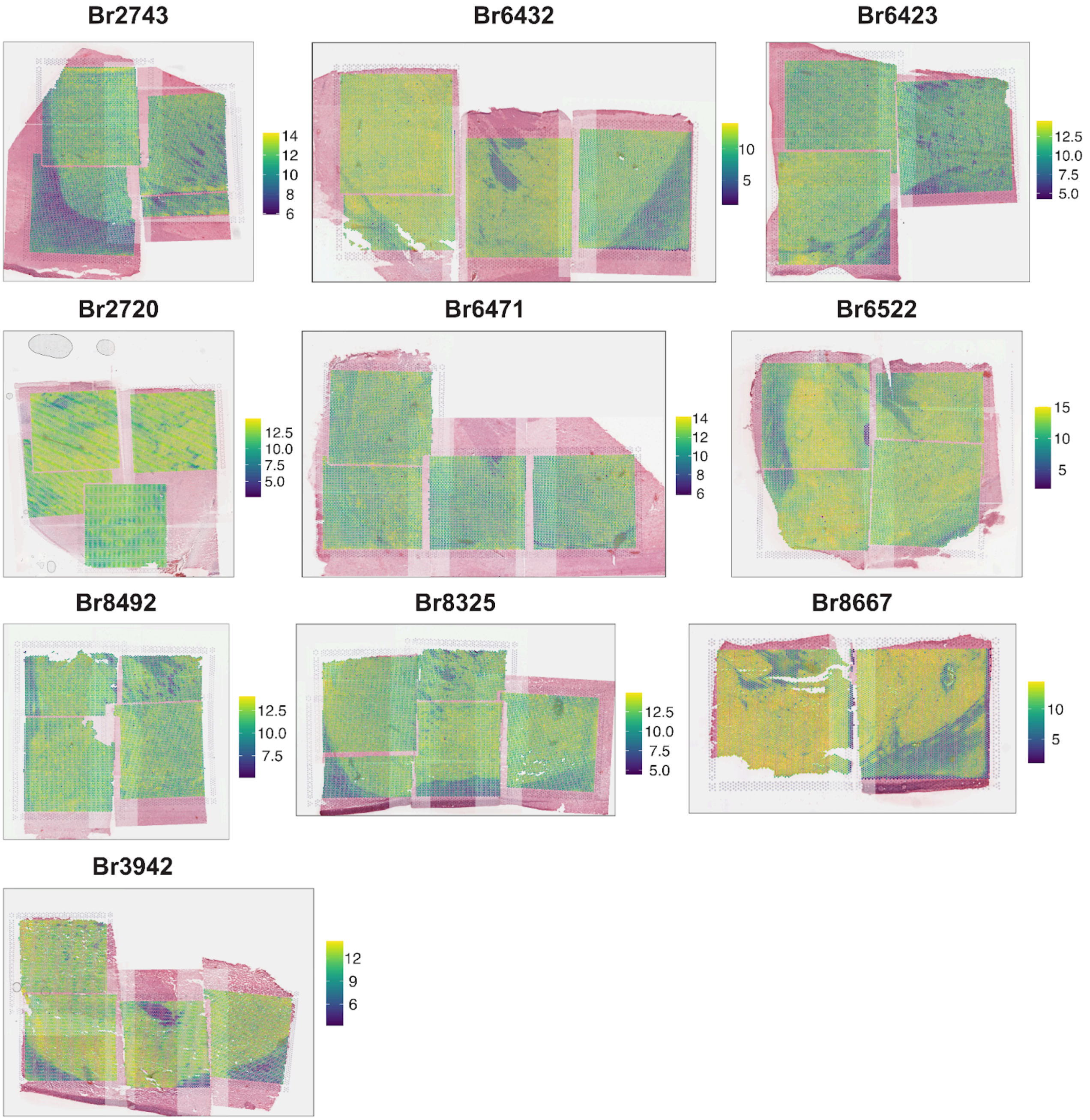


**Supplementary Figure. 12. Spatial distribution of library size per spot across donors.** H&E images of each of the 10 donors overlaid with Visium spots that are colored by the $\log_2$(sum_umi per spot). Donors are ordered based on their positions along the anterior-posterior (A-P) axis, (top-left → bottom-right). Color bars show per-donor scales. Images are oriented with dorsal at the top and lateral to the right (medial left, ventral bottom). Regions more likely to be white matter (WM) based on histology exhibited lower $\log_2$(`sum_umi` per spot) compared to gray matter spots.

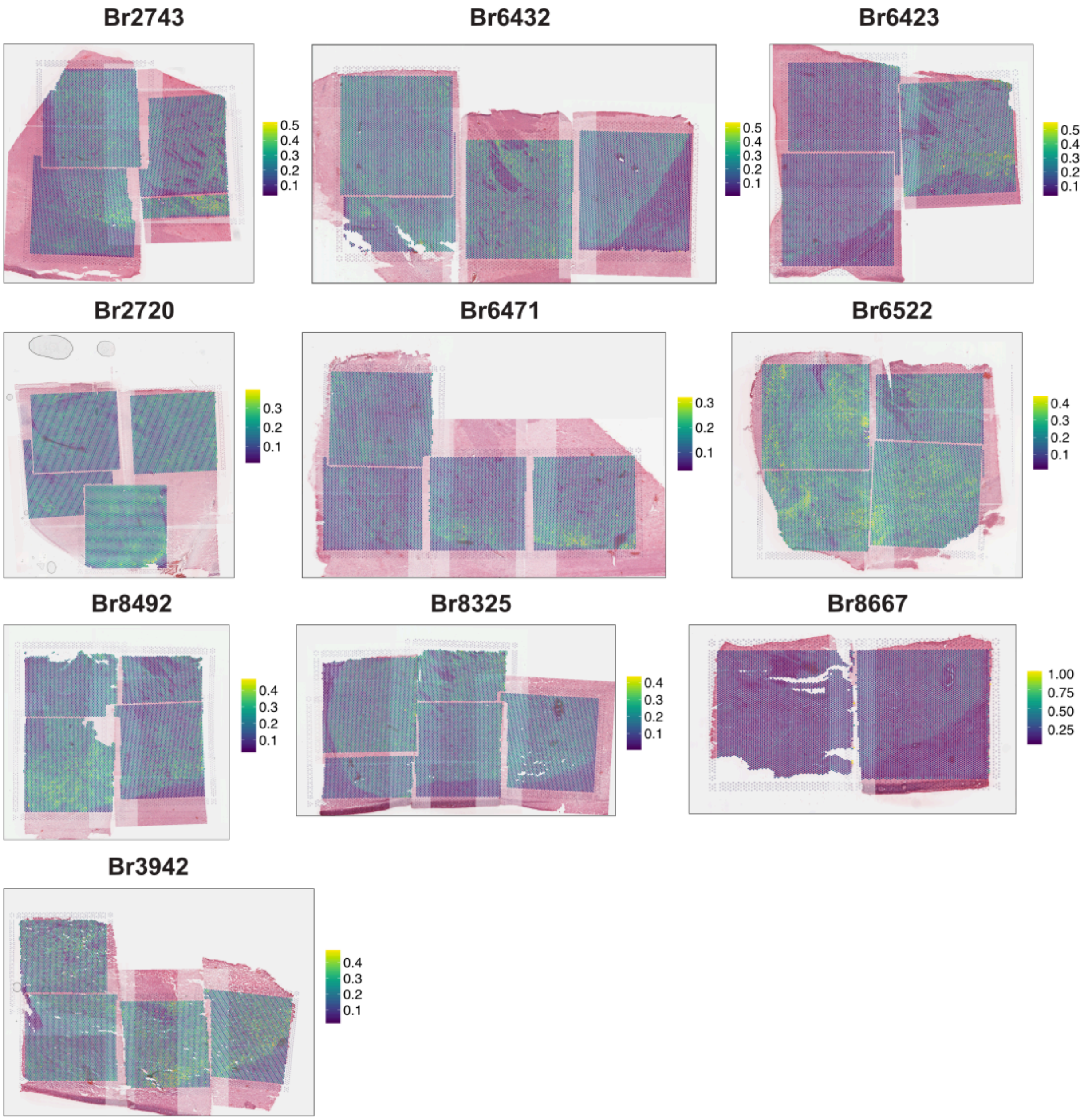


**Supplementary Figure. 13. Spatial distribution of ratio of mitochondrial gene expression (expr_chrM_ratio) across donors.** H&E images for each of the 10 donors are overlaid with Visium spots colored by expr_chrM_ratio. Donors are ordered along the A-P axis (top-left → bottom-right). Color bars show per-donor scales. Images are oriented with dorsal at the top and lateral to the right (medial left, ventral bottom). Regions more likely to be white matter (WM) based on histology exhibited lower `expr_chrM_ratio` when compared to gray matter.

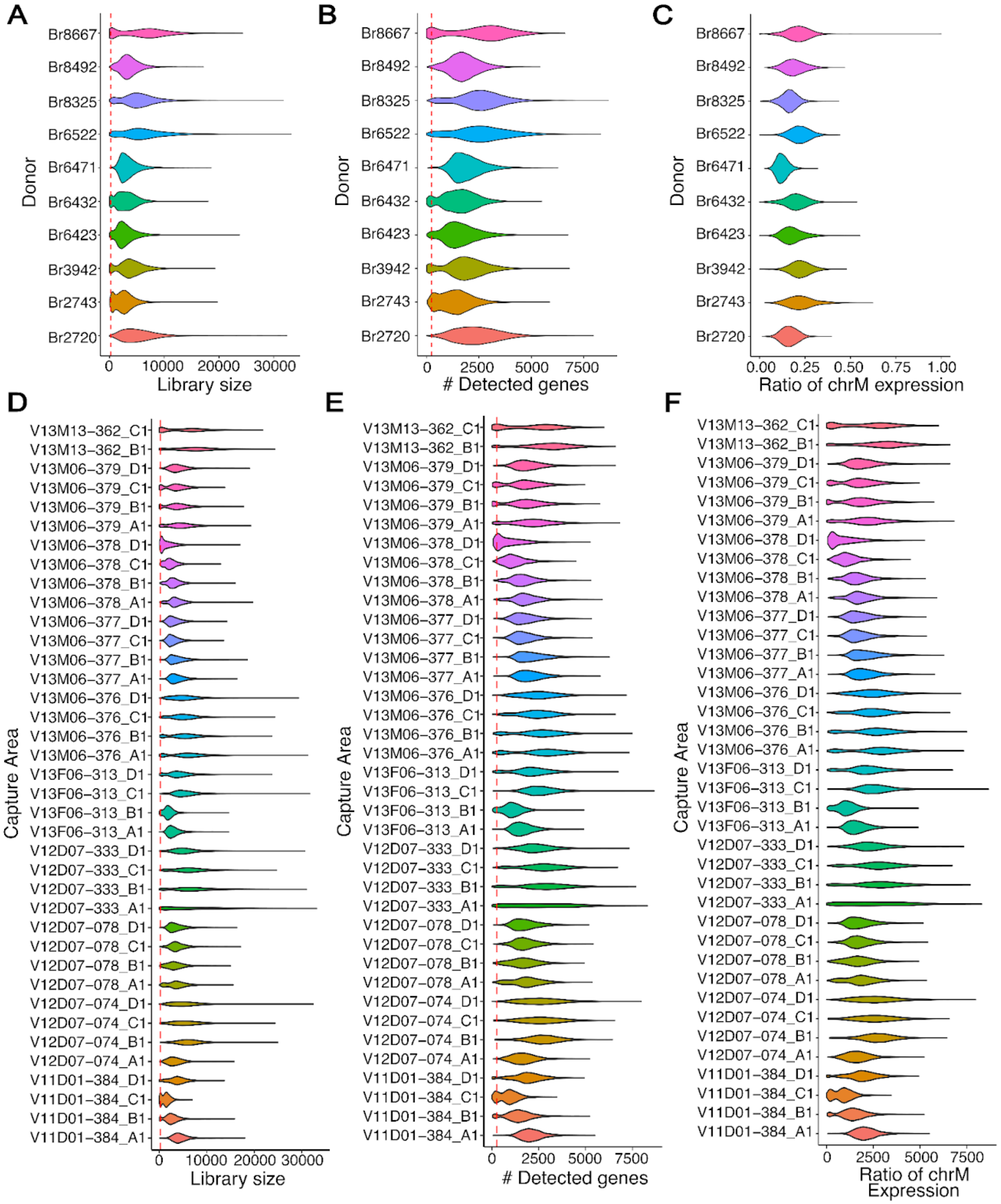


**Supplementary Figure. 14. Per-spot QC metrics across donors and capture areas. (A-C)** Violin plots by donor showing **(A)** Library size (sum UMIs per spots), **(B)** Number of detected genes per spot, and **(C)** Ratio of mitochondrial (chrM) gene expression **(D-F)** The same metrics stratified by capture area. Red dashed lines mark QC cutoffs used for spot filtering: 250 UMIs for library size (**A**, **D**), and 250 genes for detected genes (**B**, **E**).

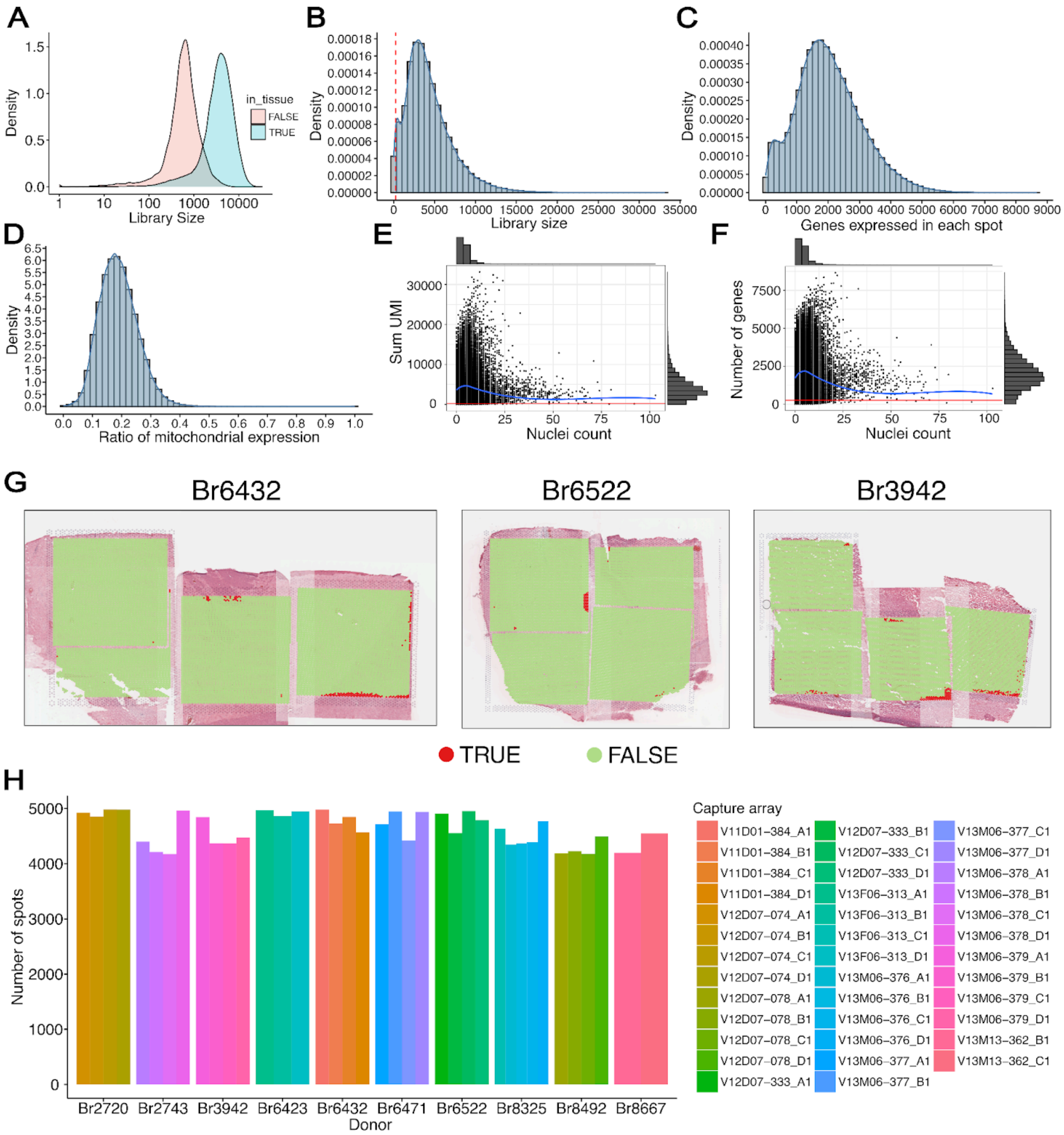


**Supplementary Figure. 15. Spot-level QC and filtering of SRT data.**
**(A)** Density of library size (sum UMIs/spot) for spots annotated to be in-tissue compared to spots annotated to be not in-tissue. Only in-tissue spots were retained for downstream analyses.
**(B)** Global distribution of library size with the QC cutoff indicated (red dashed line, 250 UMIs).
**(C)** Global distribution of detected genes per spot with the QC cutoff indicated (red dashed line, 250 genes).
**(D)** Global distribution of the mitochondrial expression fraction.
**(E)** Relationship between the library size and number of nuclei detected in each spot by *Vistoseg*.
**(F)** Same as **(E)**, but for the relationship between number of detected genes and number of nuclei per spot.
**(G)** Examples from three donors (Br6432, Br6522, Br3942) illustrating low-quality spots (red; `sum_umi` < 250 and `edge_distance` < 6) that were excluded from downstream analyses and retained spots shown in green.
**(H)** Number of spots retained after QC, stratified by donor and capture area.

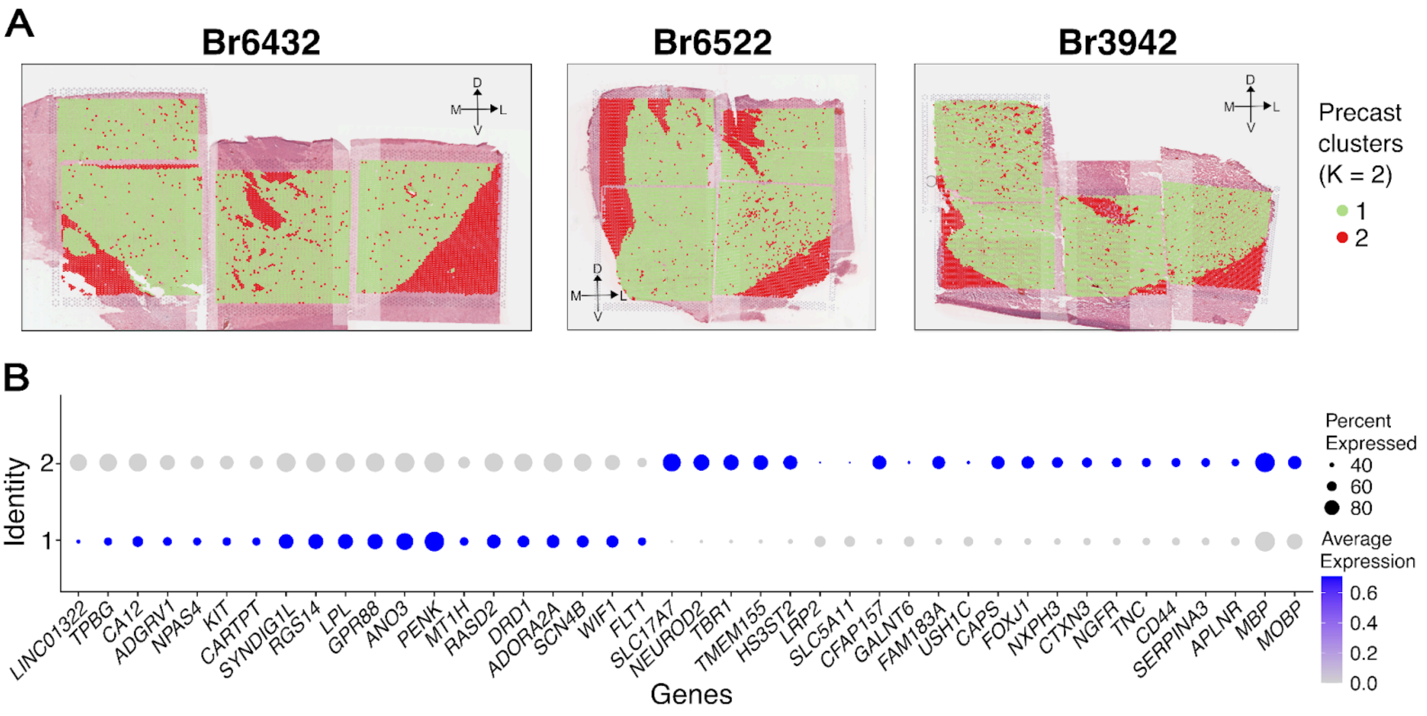


**Supplementary Figure. 16. *PRECAST* (k=2) clustering separates white and gray matter.** **(A)** Spot plots showing *PRECAST* cluster assignments (k = 2) using default spatially variable genes (SVGs) without controlling for broad differences between white matter (WM) and gray matter, across three representative donors (Br6432, Br6522, Br3942). Spots are colored by the corresponding cluster assignment (1 = Green, 2 = Red). Images are oriented with dorsal at the top and lateral to the right. Cluster 2 spots (red) localize to regions predicted to be WM based on histology.
**(B)** Dot plot of top 20 DEGs identified between *PRECAST* clusters 1 and 2, along with *MBP* and *MOBP*. Dot size indicates the percent of spots expressing the gene and the color corresponds to the average gene expression across all spots belonging to a particular cluster. Cluster 1 shows MSN/inhibitory neuron markers (e.g., *PENK*, *ADORA2A*, *DRD1*, *KIT*), whereas Cluster 2 is enriched for myelin genes (*MBP*, *MOBP*), confirming WM identity.

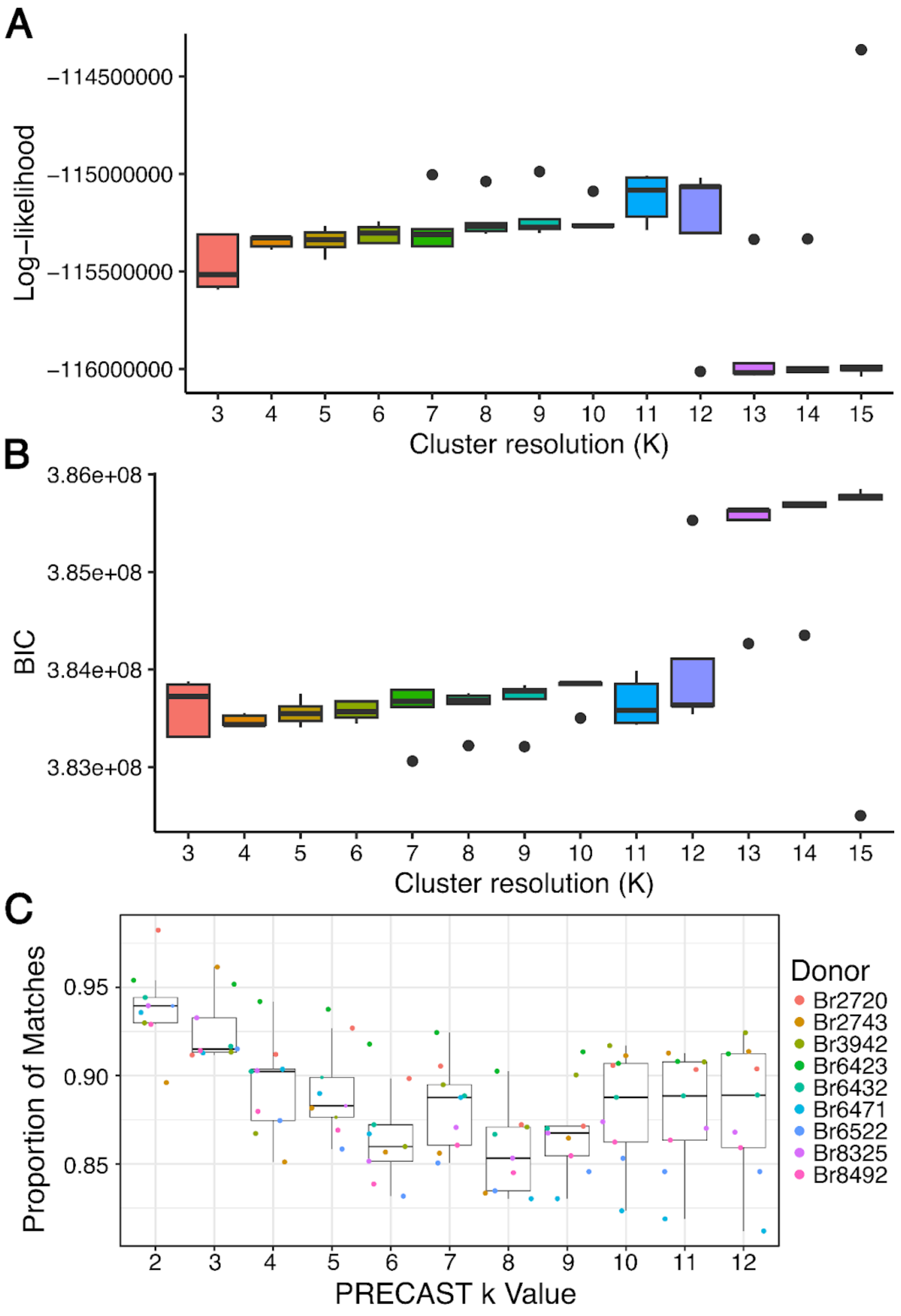


**Supplementary Figure. 17. *PRECAST* model selection and clustering concordance.**
**(A)** Log-likelihood of *PRECAST* model fit (y-axis) across clustering resolutions (k=3-15, x-axis). Boxplots summarize model fit across five independent random initializations.
**(B)** Bayesian information criterion (BIC, y-axis) across clustering resolutions (k=3-15, x-axis). Boxplots summarize BIC across five independent random initializations. Lower values indicate an optimal trade-off between model fit and model complexity.
**(C)** Concordance of cluster labels for overlapping spots which act as technical replicates (y-axis) as a function of resolution (k, x-axis), shown per donor (points) with boxplots summarizing across donors for random start 3. Higher values indicate better agreement between cluster assignments of overlapping spots.

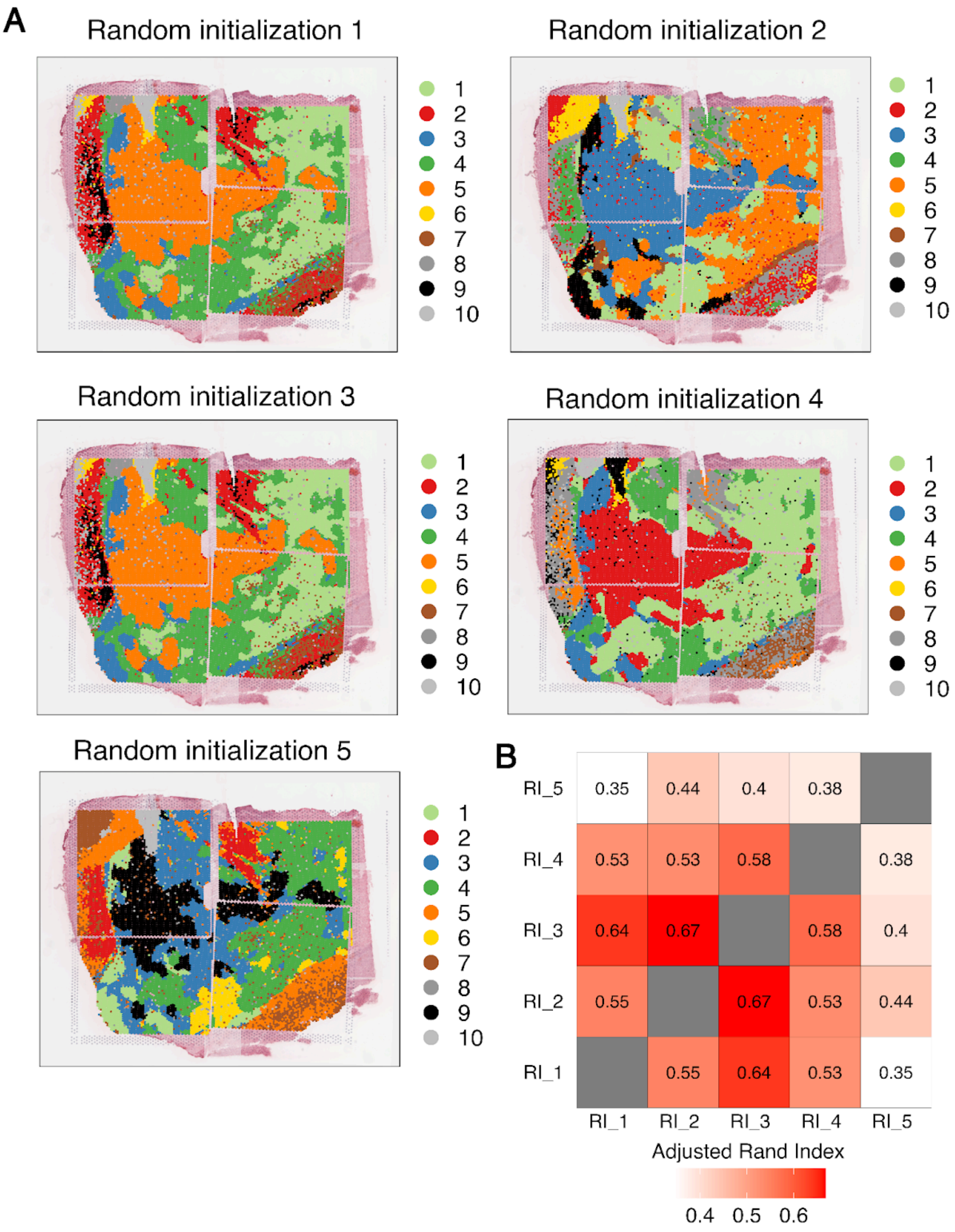


**Supplementary Figure. 18.** ***PRECAST*** **clustering stability across random initializations (k = 10).**
**(A)** Spot plots showing *PRECAST* cluster assignments for the same tissue section (Br6522) across five random starts (RI_1–RI_5); colors denote cluster labels 1–10. All images are oriented with dorsal at the top and lateral to the right.
**(B)** Pairwise adjusted Rand index (ARI) between the five independent random initializations. Higher values indicate greater agreement. The highest concordance is observed for RI_3 with RI_1 (ARI = 0.64) and RI_2 (ARI = 0.67), supporting the choice of RI_3 for downstream analyses.

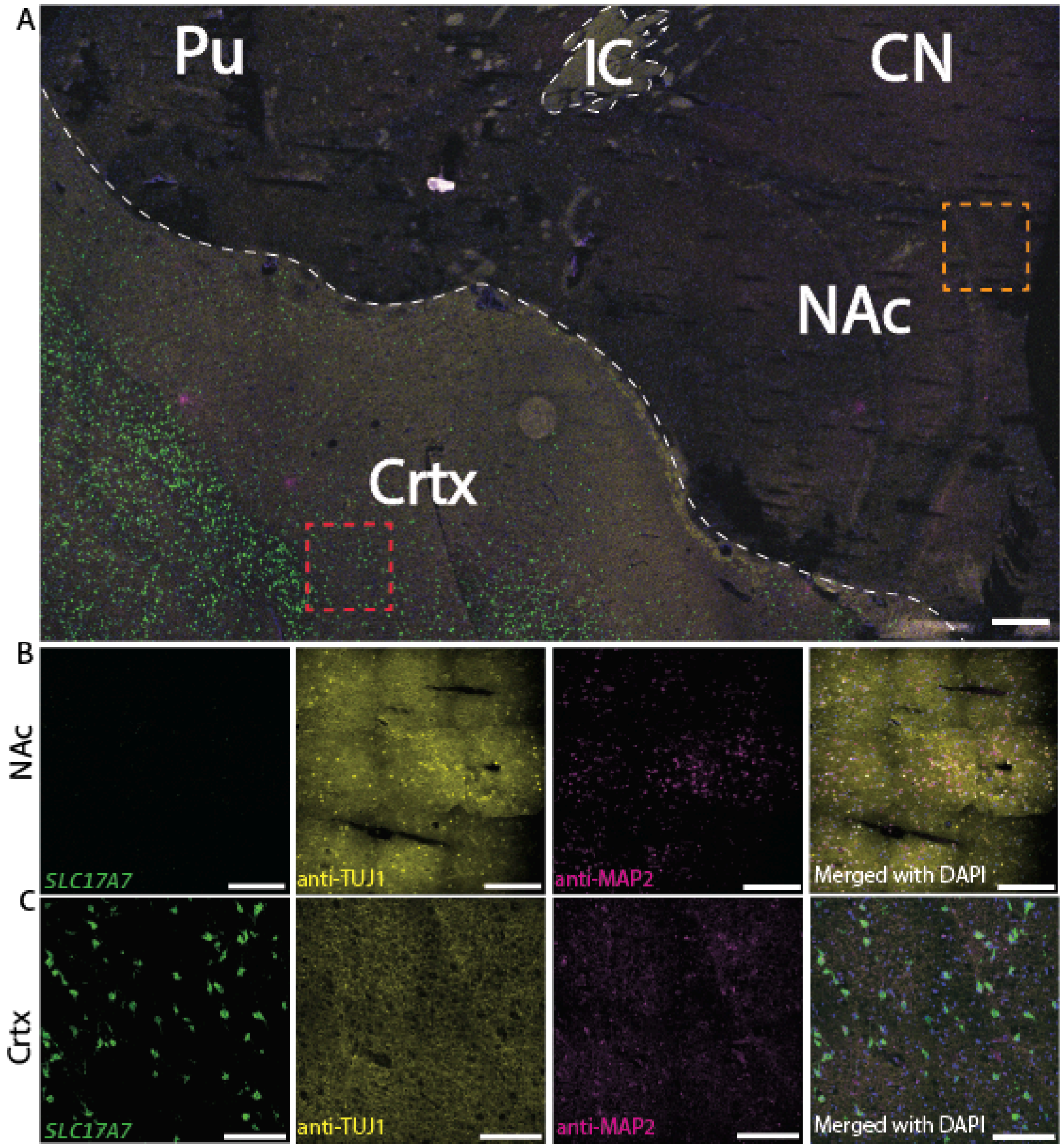


**Supplementary Figure. 19. *SLC17A7* expression is restricted to the cortex and absent in the NAc.** **(A)** 2x RNAScope-IF image from donor Br6423 illustrating anatomical locations of the NAc, including cortex (Crtx), adjacent caudate nucleus (CN), putamen (Pu), and internal capsule (IC). RNAScope probe was used to mark *SLC17A7* gene expression (green), while antibodies against TUJ1 (yellow) and MAP2 (magenta) were used to denote axons/dendrites or dendrites, respectively. DAPI (blue) was used to denote nuclei. Scale bar 1000 μm. **(B)** Zoomed in 40x images of the NAc (orange square as shown in **(A)**) reveals strong signal for TUJ1 (yellow) and MAP2 (magenta) but no *SLC17A7* staining. Scale bar 100 μm. **(C)** Zoomed in 40x images of the Crtx (red square as shown in **(A)**) reveals strong signal for TUJ1 (yellow), MAP2 (magenta), and *SLC17A7* (green) staining. Scale bar 100 μm.

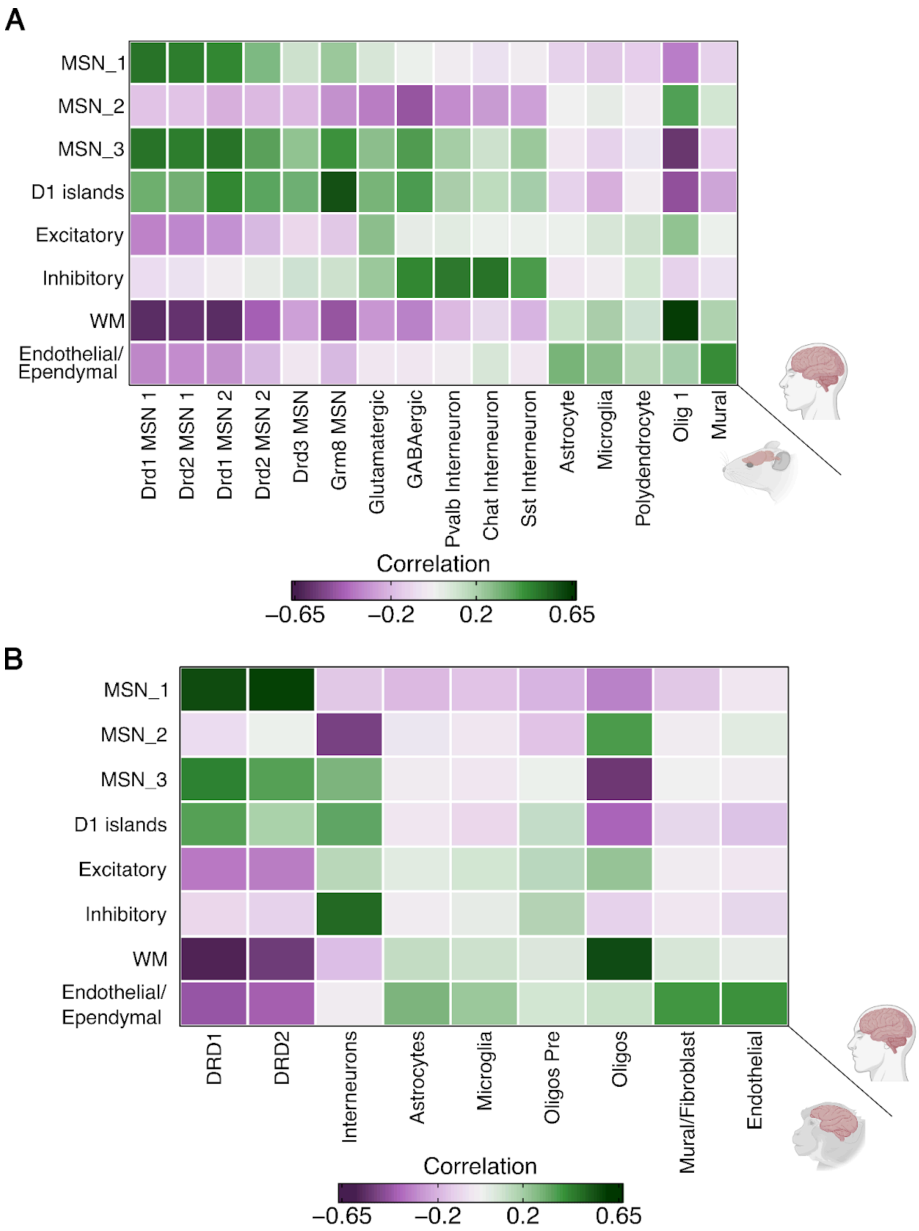


**Supplementary Figure. 20. Cross-species registration of human SRT-derived spatial domains (SpDs) to neuronal and glia rodent and NHP cell types.(A)** Heatmap of Pearson correlations between per-gene t-statistics from one-vs-all contrasts computed for each human SpD (rows) and each rodent reference [26] cell type (columns), evaluated over the union of the top 250 DEGs per SpD mapped to 1:1 human–rodent orthologs present in both datasets. **(B)** Same as **(A),** but comparing SpDs (rows) to the non-human primate (NHP) reference [24] across all annotated cell classes (columns), using 1:1 human–macaque orthologs. Tiles show correlation coefficients (green = positive; purple = negative).

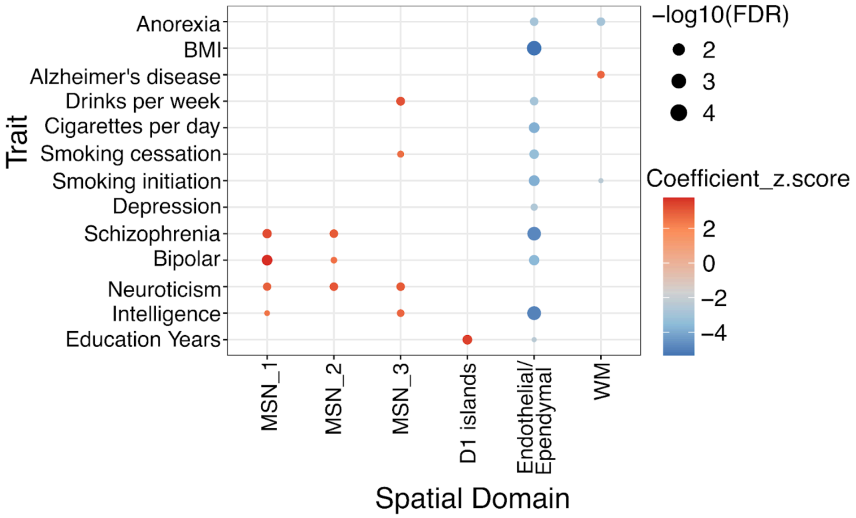


**Supplementary Figure. 21. Heritability enrichment of genes corresponding to spatial domains (SpDs) in the NAc.** Dot plot summarizing significant (FDR < 0.1) stratified LD score regression (s-LDSC) results across neuropsychiatric and behavioral traits for gene sets derived from SRT-derived SpDs. Dot color encodes the regression coefficient z-score (red = enrichment; blue = depletion) and dot size encodes $-\log_{10}$(FDR). Notable enrichments include schizophrenia, bipolar disorder, and neuroticism in MSN_1 and MSN_2, and drinks per week and smoking cessation in MSN_3.

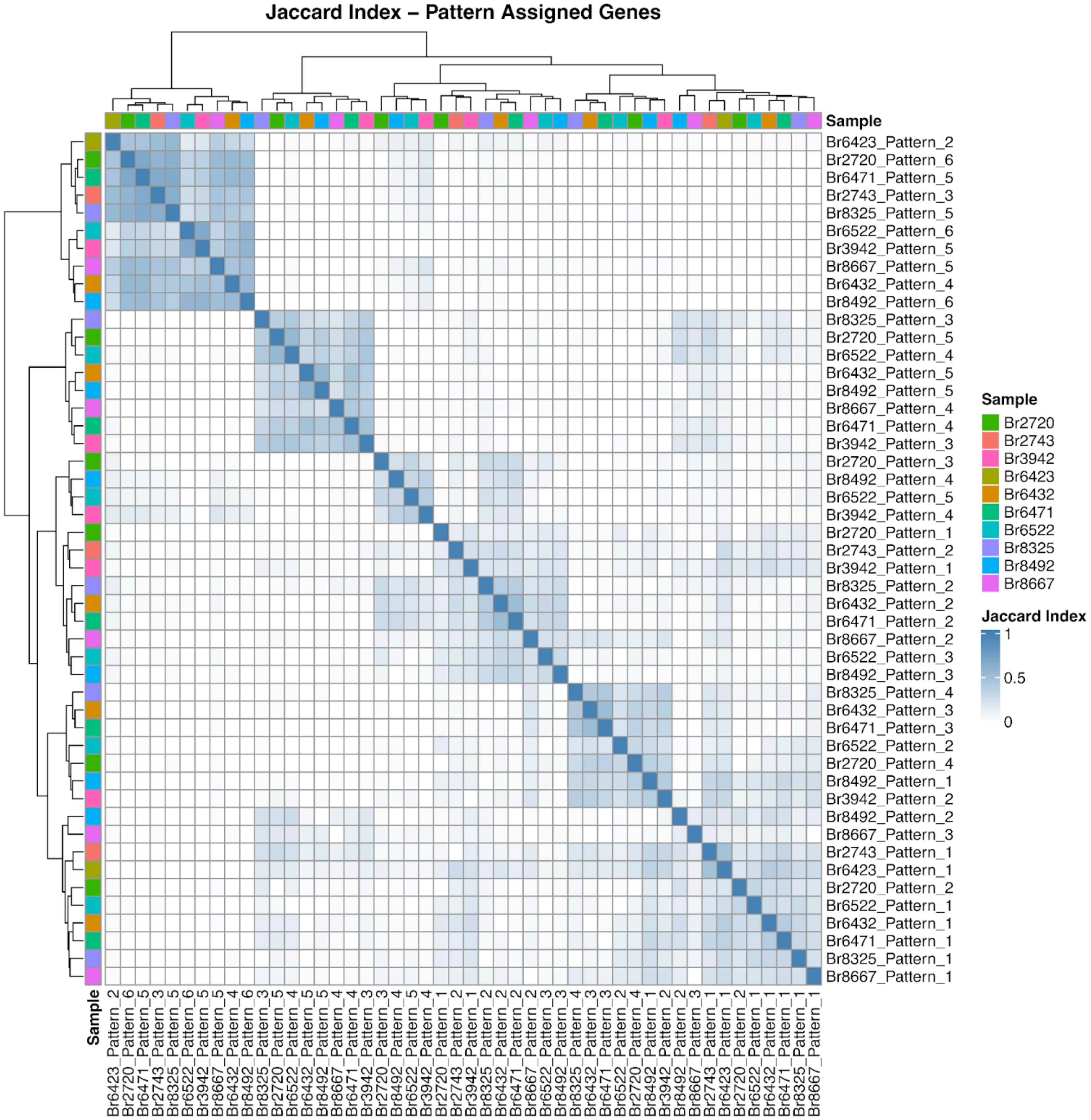


**Supplementary Figure. 22. Gene-level similarity of *MERINGUE* spatial patterns across donors.** Heatmap of pairwise Jaccard indices (0–1) between the sets of genes assigned to each donor-specific *MERINGUE* pattern (labels: donor_pattern_i). A value of 1 indicates identical gene sets while 0 indicates no overlap. Rows and columns are ordered by hierarchical clustering (Ward's method) and colored side bars denote donors.

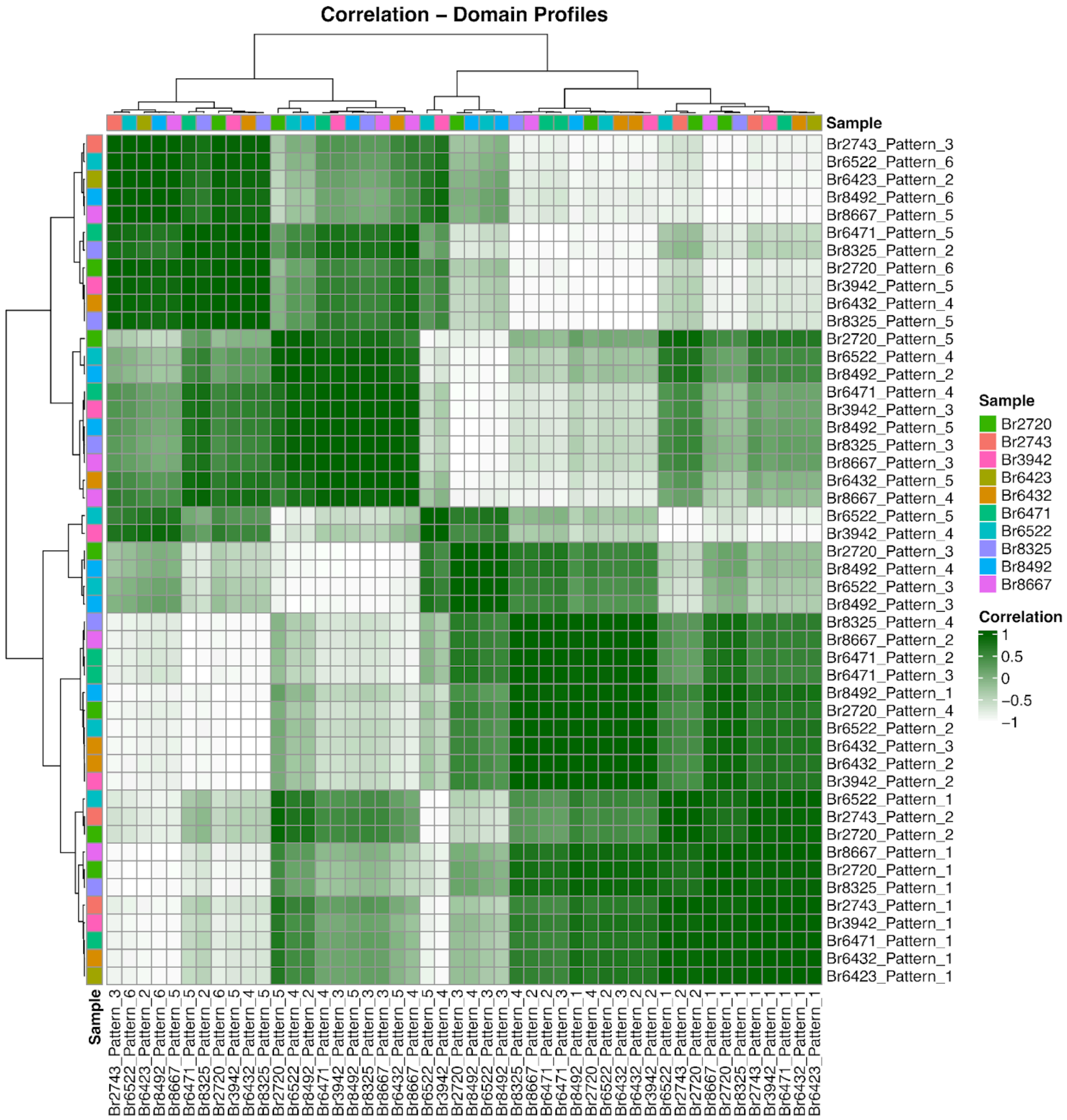


**Supplementary Figure. 23. Domain-level similarity of *MERINGUE* spatial patterns across donors.** Heatmap of Pearson correlations between domain averaged profiles for each donor-specific *MERINGUE* pattern (labels: donor_pattern_i). For each pattern, factor scores were averaged within the *PRECAST*-defined MSN domains (MSN_1-3) to form a domain averaged profile, and profiles were correlated across patterns. Rows and columns are ordered by hierarchical clustering (Ward's method) and side bar colors denote donors.

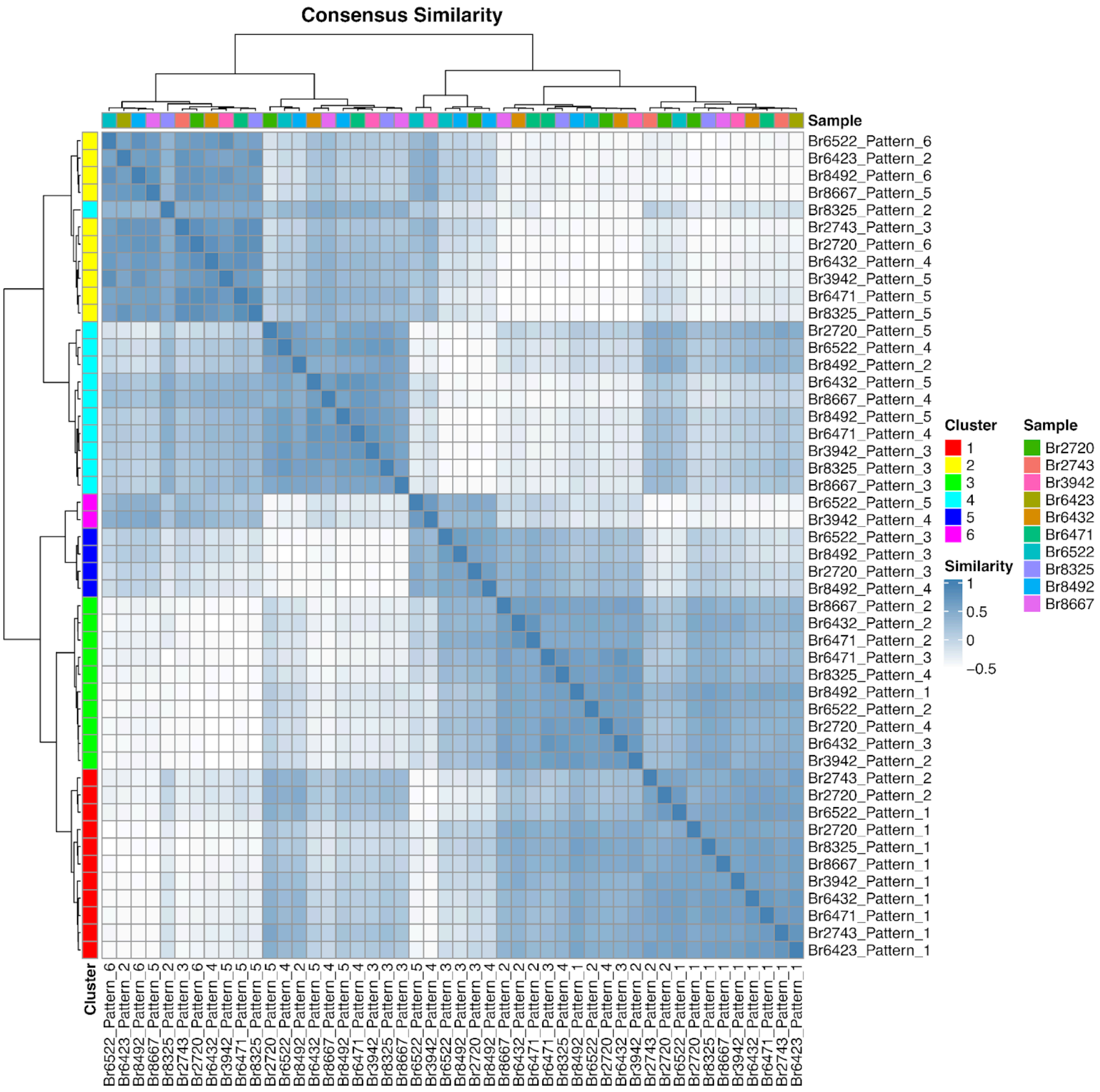


**Supplementary Figure. 24. Consensus similarity of *MERINGUE* patterns and *MERINGUE* Consensus Pattern (MCP) grouping.**

Heatmap showing the consensus similarity between all donor-specific *MERINGUE* patterns, computed as the mean of (i) the gene-level Jaccard overlap of pattern-assigned genes and (ii) the correlation of domain activation profiles (MSN_1–3). Rows/columns are ordered by Ward hierarchical clustering. The left color bar marks clusters obtained by cutting the dendrogram at k = 6 (used to define *MERINGUE* consensus patterns, MCPs); top color bar indicates donors. These clusters were consolidated into four MCPs detected across multiple donors for downstream analyses.

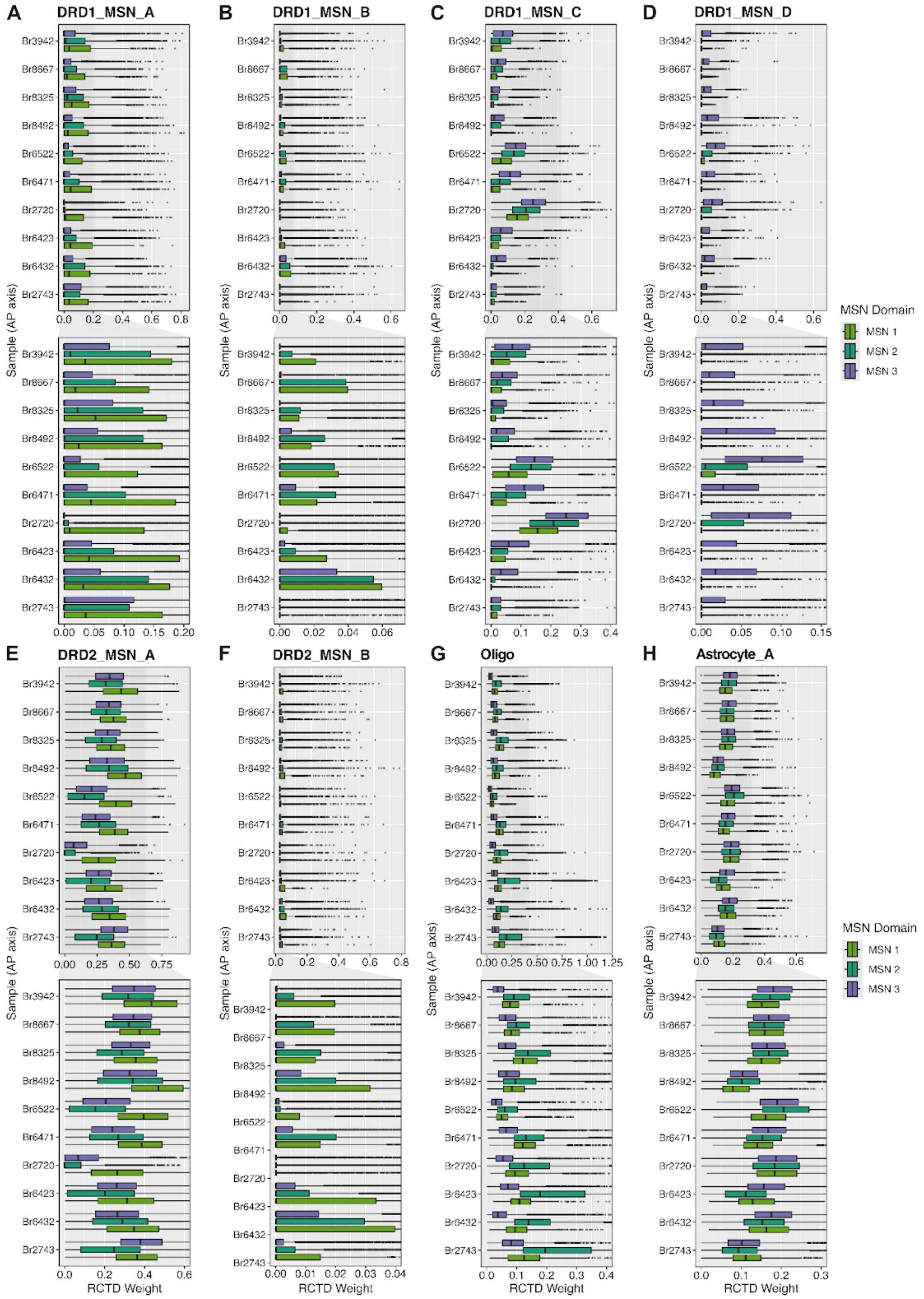


**Supplementary Figure. 25. *RCTD*-derived cell-type composition across MSN spatial domains.** Boxplots show the distribution of spot-level *RCTD* weights for eight cell types with non-zero contributions across MSN domains, grouped by donor and stratified by domain (colors: MSN_1-3 SpDs). Donors are ordered along the A–P axis (bottom = most anterior, top = most posterior). Panels: **(A)** DRD1_MSN_A, **(B)** DRD1_MSN_B, **(C)** DRD1_MSN_C, **(D)** DRD1_MSN_D, **(E)** DRD2_MSN_A, **(F)** DRD2_MSN_B, **(G)** Oligodendrocytes, **(H)** Astrocytes. Consistent biases are evident: DRD1_MSN_A and DRD2_MSN_A are highest in MSN_1 SpD, DRD1_MSN_C is enriched in MSN_3 SPD, oligodendrocytes increase in MSN_2 SpD, DRD1_MSN_D appears within MSN_3 SpD in several donors.

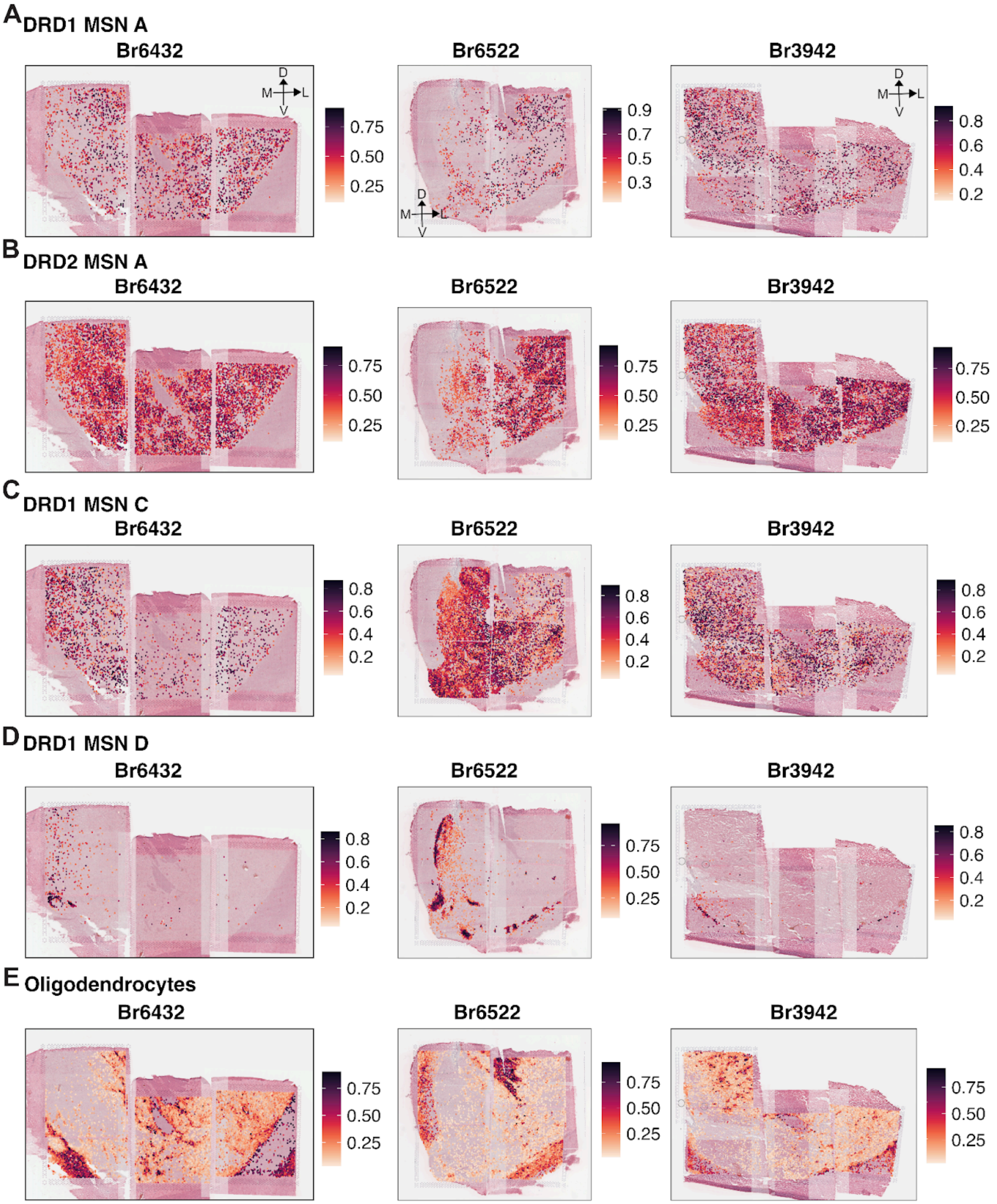


**Supplementary Figure. 26. Spatial maps of *RCTD* weights across the A-P axis.**
Spot plots of *RCTD*-derived cell-type weights overlaid on H&E for three representative donors ordered along the A–P axis: Br6432 (anterior), Br6522 (intermediate), Br3942 (posterior). All images are oriented with dorsal at the top and lateral to the right. Color indicates the *RCTD* weight per spot. **(A)** DRD1_MSN_A. **(B)** DRD2_MSN_A. **(C)** DRD1_MSN_C. **(D)** DRD1_MSN_D. **(E)** Oligodendrocytes. Consistent trends are visible across donors: DRD2_MSN_A shows a lateral bias and DRD1_MSN_C is enriched medially

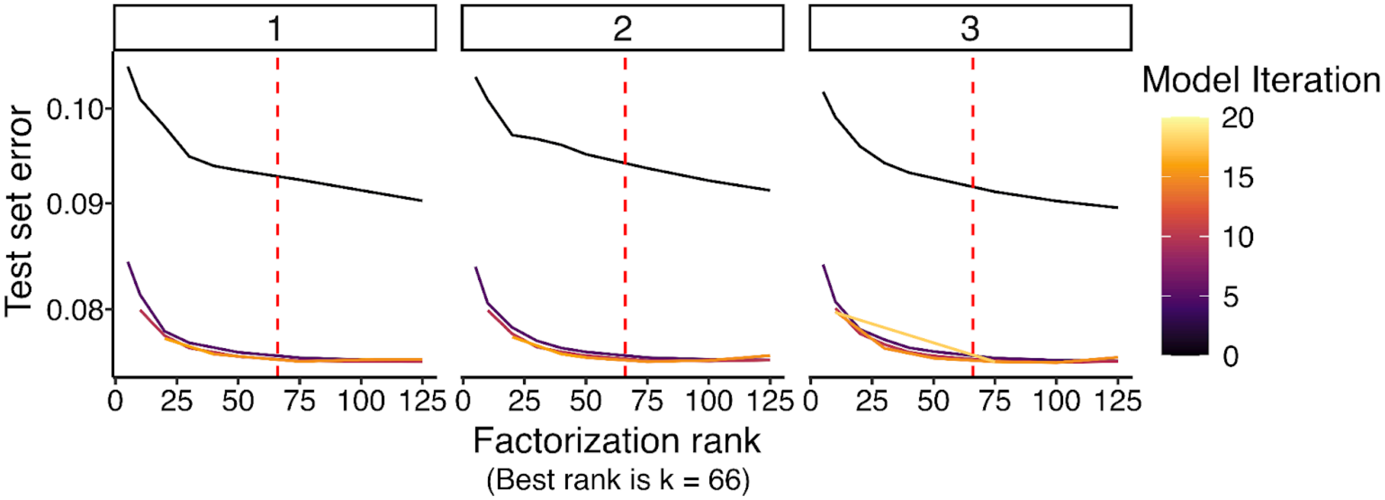


**Supplementary Figure. 27. NMF rank selection through cross-validation.**
Held-out test error (y-axis) versus factorization rank (x-axis) for three independent cross-validation replicates (panels 1–3). For each replicate, 20% of entries were masked and used to compute test error; colored curves show the error over solver iterations. Error decreases and then plateaus with increasing rank and the red dashed line marks the selected rank k = 66, the smallest value beyond which additional factors yield minimal improvement and results are stable across replicates.

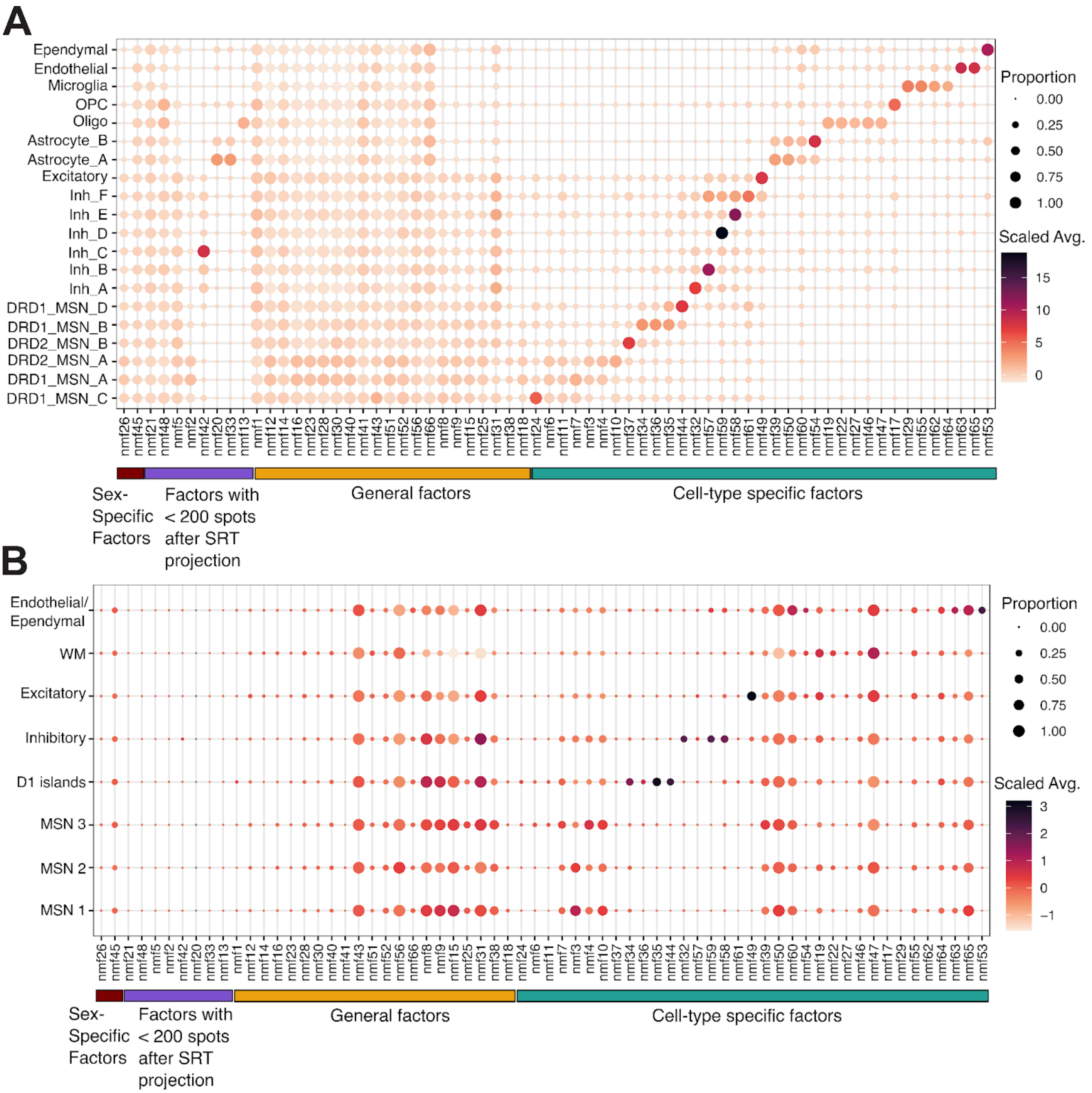


**Supplementary Figure. 28. NMF factor activity summarized across cell types and spatial domains.** **(A)** Dot plot of NMF factors across snRNA-seq cell types. Dot size shows the proportion of nuclei with non-zero factor loading; color shows the z-scored mean loading (scaled average).
**(B)** The same summary after projecting factors to SRT and aggregating within spatial domains (SpDs). Dot size shows the proportion of spots with non-zero projection; color shows the z-scored mean projection weight. Factors correlated with donor sex (nmf26, nmf45) and factors detected in <200 SRT spots were removed. SpD-associated factors were defined as those with >20% prevalence and scaled average >0.2. Bottom annotations indicate NMF factor categories: sex-associated, <200 spots after SRT projection, general (non-cell type-specific), and cell type-specific.

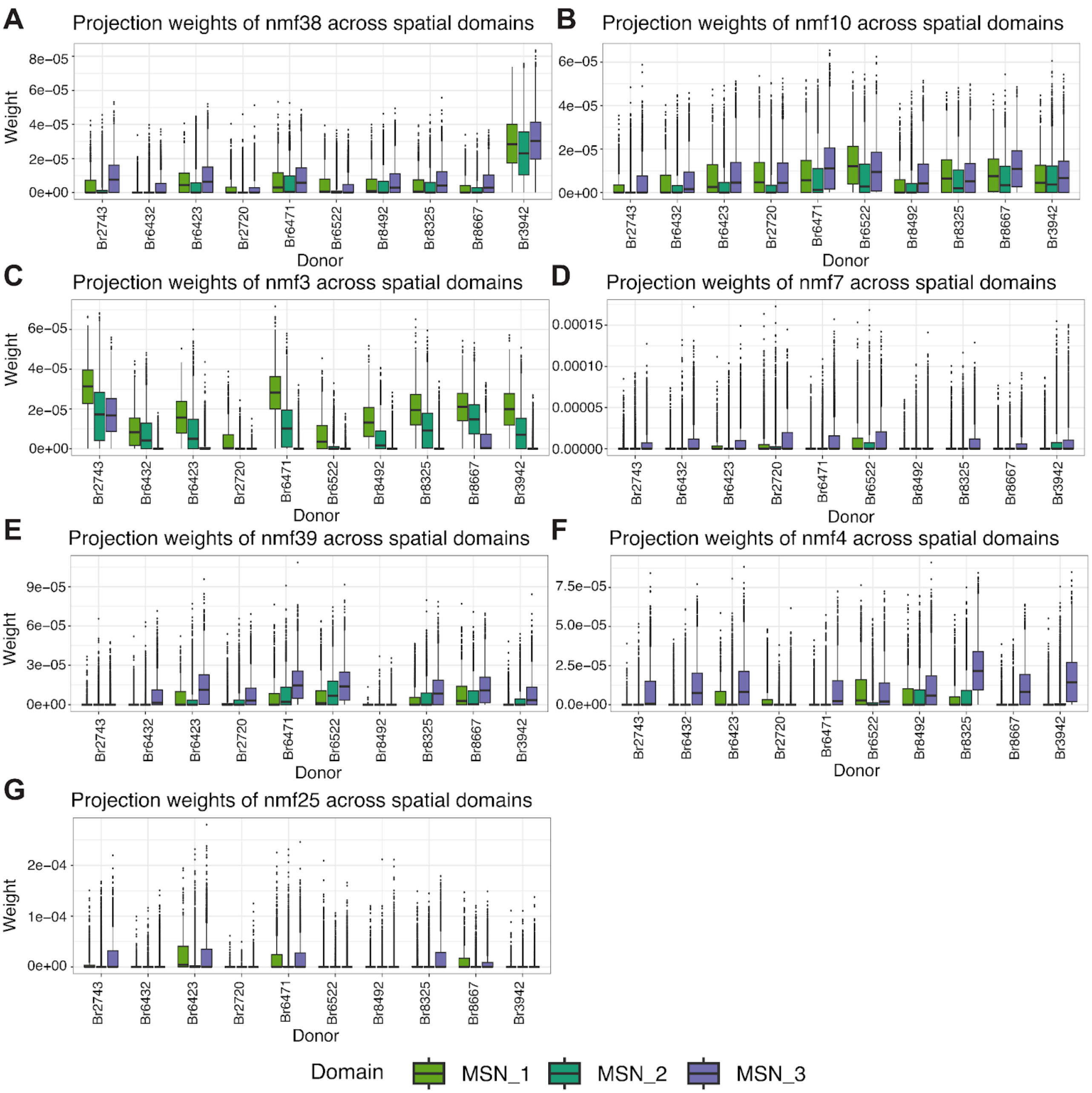


**Supplementary Figure. 29. Projection weights of MSN-specific NMF factors across SpDs.** Boxplots showing spot-level projection weights (y-axis) for MSN-associated factors across 10 donors (x-axis) stratified by MSN SpD (MSN_1-3) for **(A)** nmf38, **(B)** nmf10, **(C)** nmf3, **(D)** nmf7, **(E)** nmf39, **(F)** nmf4, and **(G)** nmf25. Donors are ordered along the A-P axis (left = anterior, right = posterior). Domain biases are evident: nmf3 is higher in MSN_1-2 SpDs (laterally-biased SpDs), nmf39, nmf4, and nmf7 are enriched in MSN_3 (medially-biased SpD), nmf10 is broadly distributed with lowest projection scores in MSN_2 SpD, while nmf38 and nmf25 show donor-biased projections.

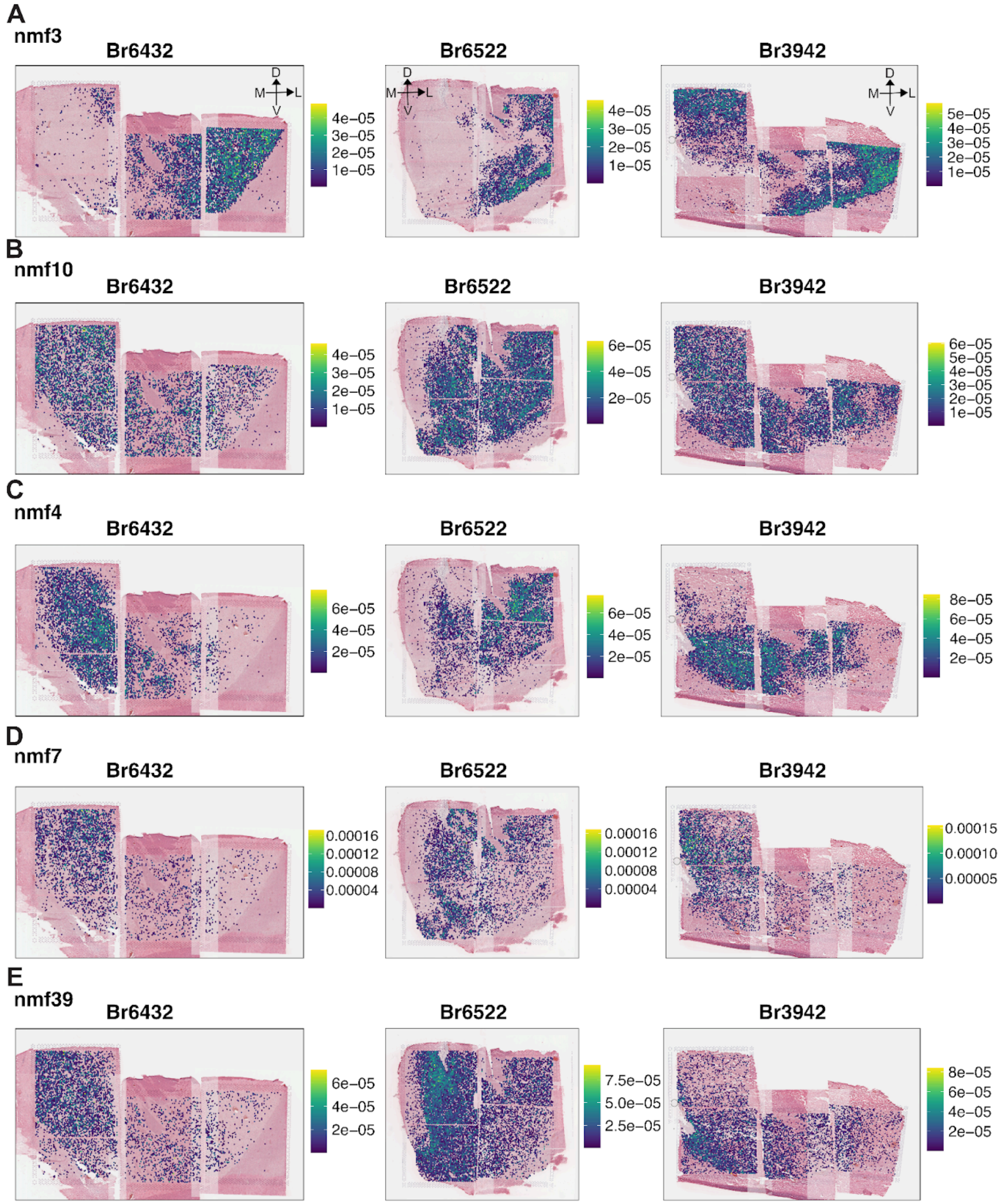


1

2 **Supplementary Figure. 30. Spatial projection of MSN-associated NMF factors.** Spot plots showing
3 projected factor weights across three representative donors: Br6432 (anterior), Br6522 (intermediate), and
4 Br3942 (posterior) for MSN-associated NMF factors **(A)** nmf3, **(B)** nmf10, **(C)** nmf4, **(D)** nmf7, and **(E)** nmf39.
5 All images are oriented with dorsal at the top and lateral to the right. Color bars indicate panel-specific scales
6 for projection weights. Consistent trends are observed, with nmf3 exhibiting a lateral bias, while nmf4 and nmf7
7 exhibit a medial bias. nmf10 is broadly distributed.

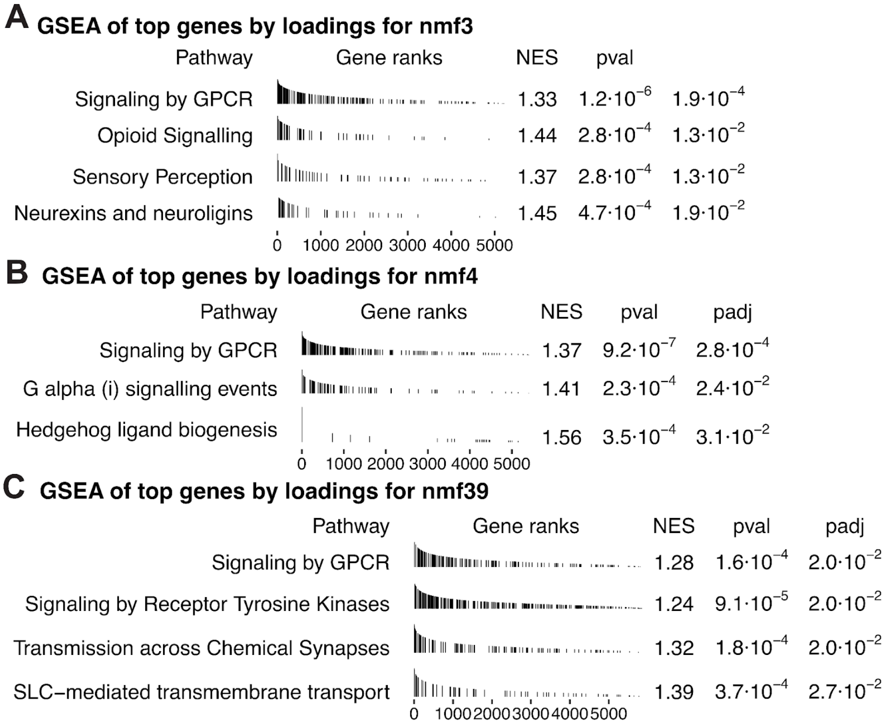


**Supplementary Figure. 31. Gene set enrichment analysis (GSEA) of MSN-associated NMF factors.** For each factor, pathways enriched among genes ranked by factor loadings are shown. Tick marks show gene ranks of pathway associated genes, and tables report normalized enrichment score (NES), nominal *p*-values, and Benjamini-Hochberg adjusted *p*-values (FDR). Significance was defined as FDR $< 0.05$. **(A)** Genes with high loadings in factor nmf3 are significantly enriched for “Signaling by GPCR”, “Opioid Signaling”, “Sensory perception”, and “Neurexins and neuroligins” Reactome pathways. **(B)** Genes with high loadings in factor nmf4 are significantly enriched for “Signaling by GPCR”, “G alpha (i) signalling events”, and “Hedgehog ligand biogenesis” Reactome pathways. **(C)** Genes with high loadings in factor nmf39 are significantly enriched for “Signaling by GPCR”, “Signaling by Recpetor Tyrosine Kinases”, “Transmission across Chemical Synapses”, and “SLC-mediated transmembrane transport” Reactome pathways.

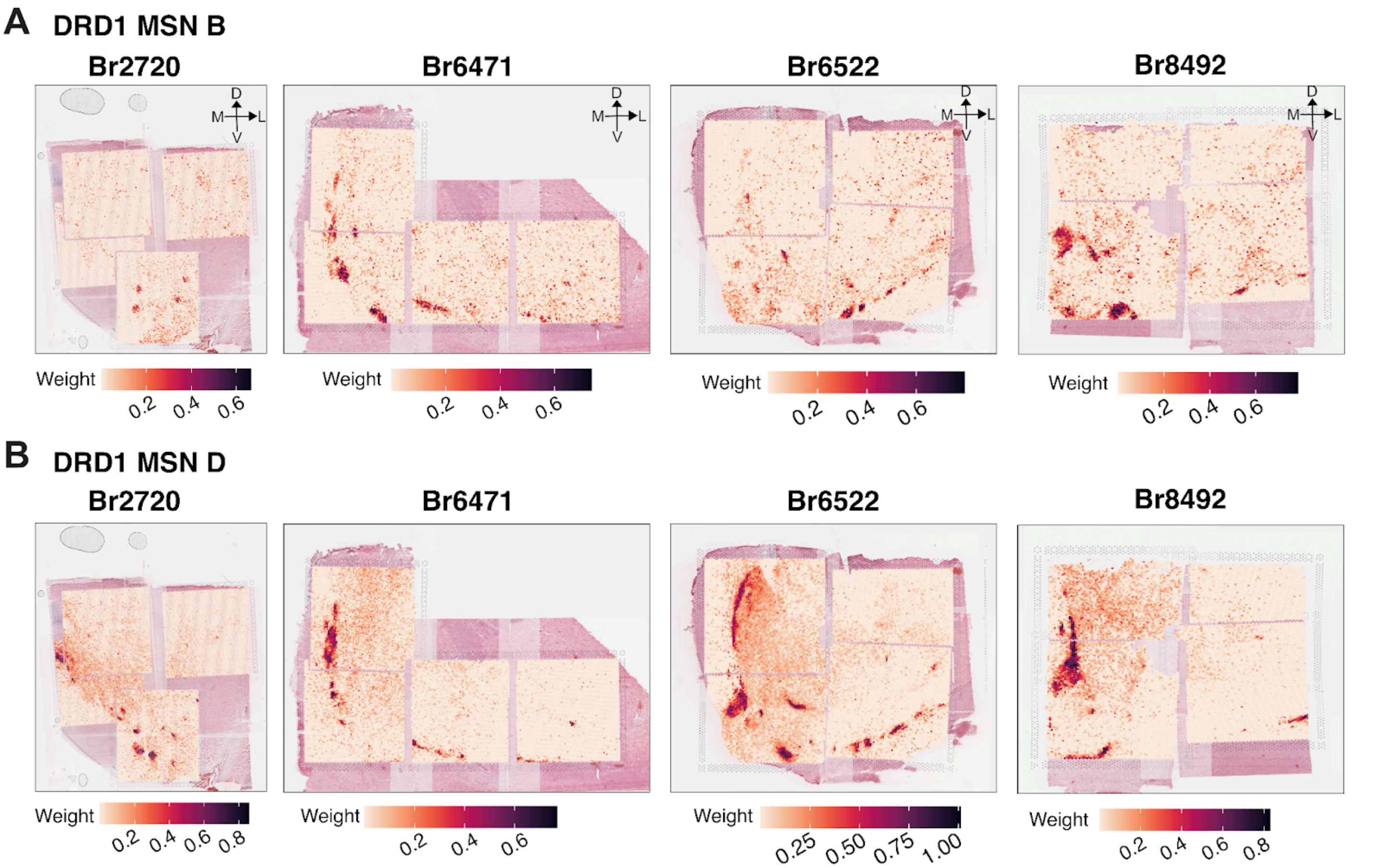


**Supplementary Figure. 32. Spatial maps of *RCTD*-derived weights for MSN subtypes found in the D1 islands.** Spot plots showing *RCTD*-derived weights corresponding to **(A)** DRD1_MSN_B and **(B)** DRD1_MSN_D for four intermediate-position donors (Br2720, Br6471, Br6522, Br8492) with prominent D1 islands. Images are oriented dorsal at the top and lateral to the right. Darker colors indicate higher *RCTD* weight (greater inferred contribution). Both subtypes localize to medial D1 islands, with DRD1_MSN_D showing broader coverage and DRD1_MSN_B enriched in ventro-medial pockets.

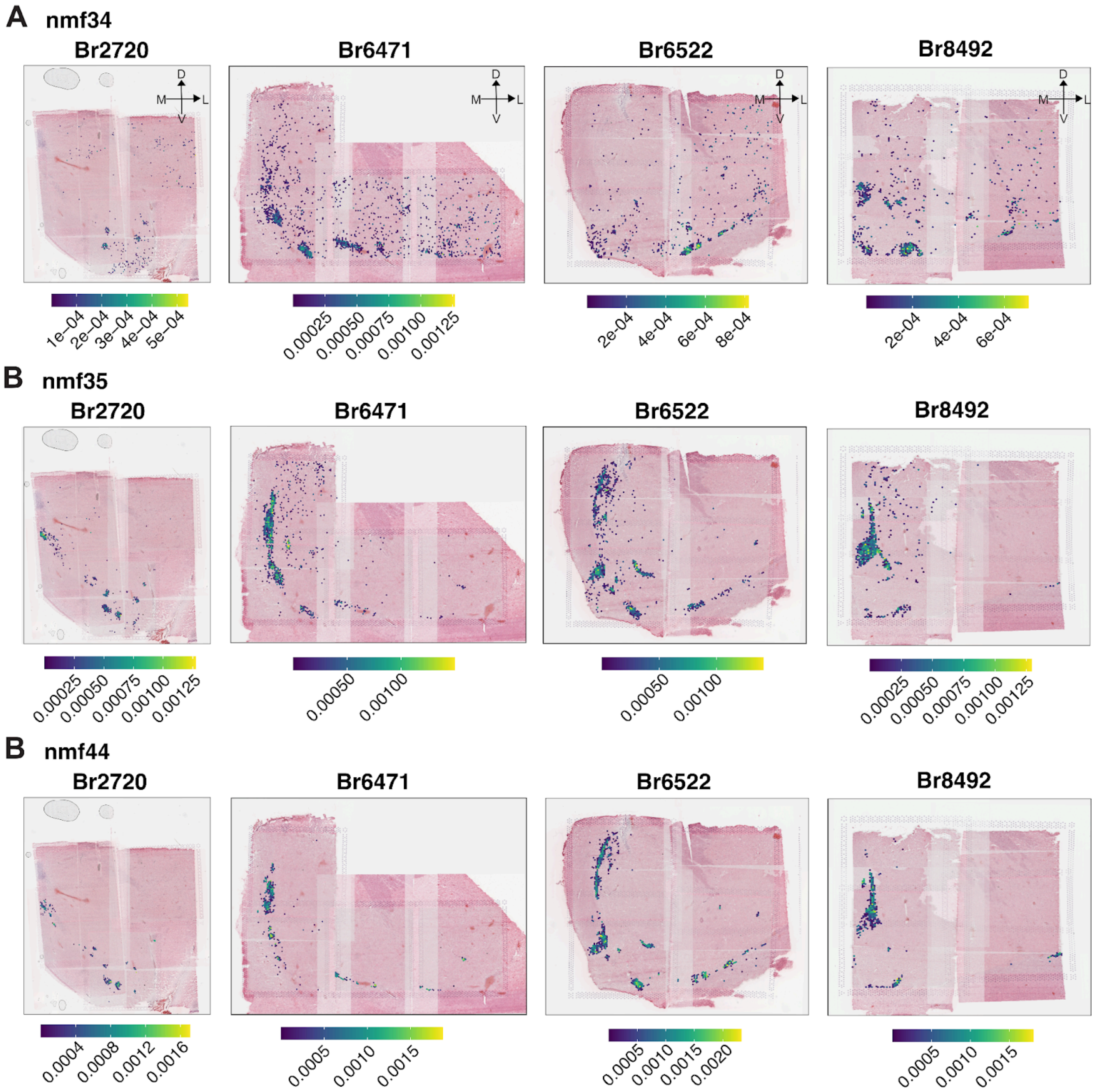


**Supplementary Figure. 33. Spatial projection of D1-islands associated NMF factors.** Spot plots showing projected factor weights for four intermediate-position donors (Br2720, Br6471, Br6522, Br8492). Images are oriented dorsal at the top and lateral to the right right, with panel-specific color scales indicating per-spot projection weight. **(A)** nmf34 marks spatially restricted ventromedial subregions within D1 islands. **(B)** nmf35 exhibits broader coverage across D1 islands. **(C)** nmf44 highlights distinct patches within D1 islands, partially overlapping with nmf34 and nmf35.

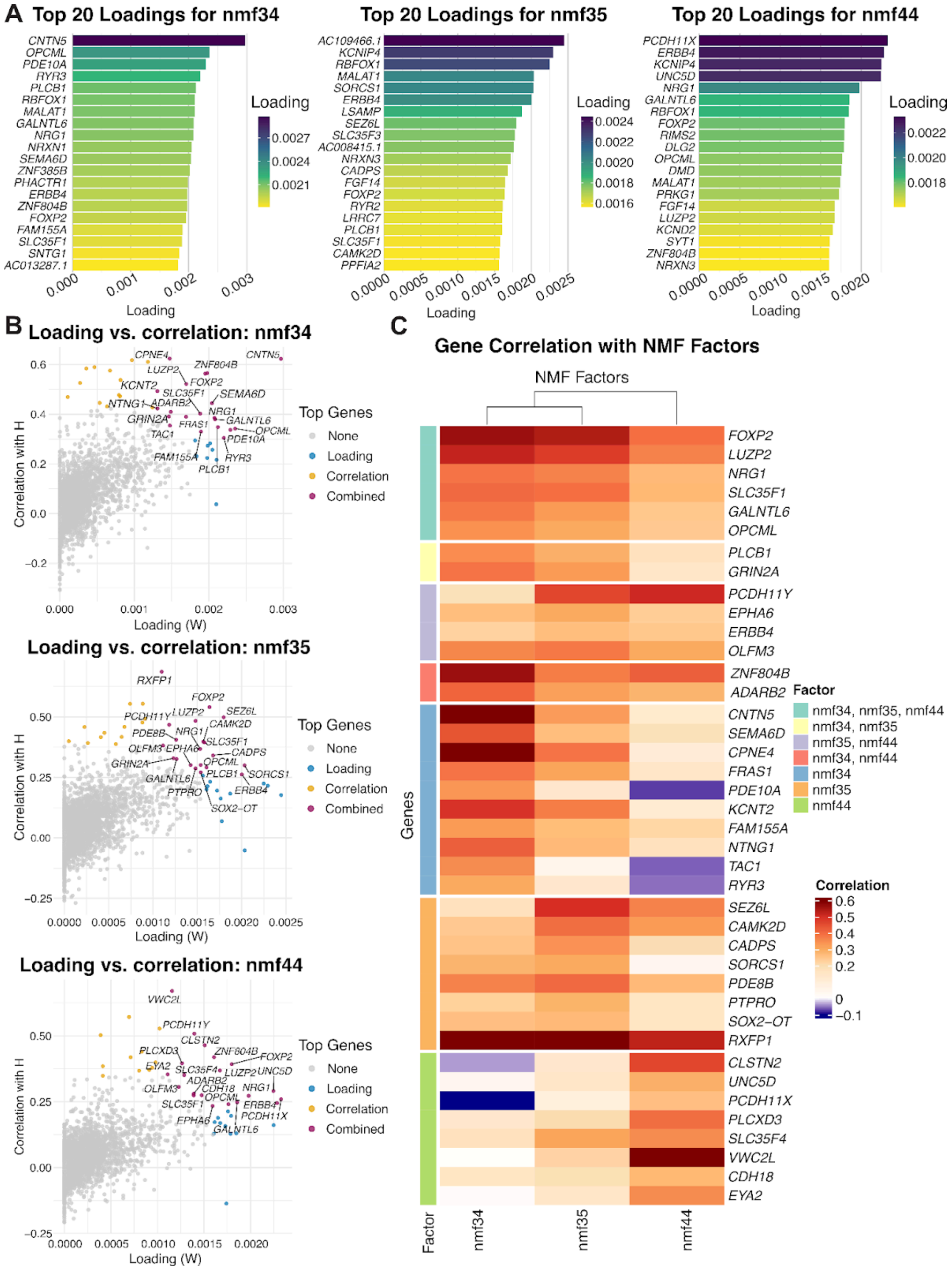


**Supplementary Figure. 34. Genes underlying D1-islands associated NMF factors in snRNA-seq.** Non-negative matrix factorization (NMF) decomposes the non-negative expression matrix $A$ (genes × nuclei) into $W$ (genes × factors; gene contributions to each latent program) and $H$ (factors × nuclei; per-nucleus program activation), such that $A \approx W \times H$. **(A)** For nmf34, nmf35, and nmf44, bar plots show the top 20 genes ranked by their loadings in $W$. **(B)** For each factor, scatter plots relate each gene's loading in $W$ (x-axis) to the correlation between that factor's $H$ scores and the gene's log-normalized expression (y-axis). Genes high on both axes are interpreted as factor-defining features. **(C)** Heatmap of correlations between $H$ and expression for the top correlation-ranked genes, highlighting shared (*FOXP2*, *NRG1*, *OPCML*) and factor-specific sets (*CPNE4/TAC1* for nmf34, *CAMK2D/SEZ6L/PDE8B* for nmf35, *VWC2L/CLSTN2* for nmf44).

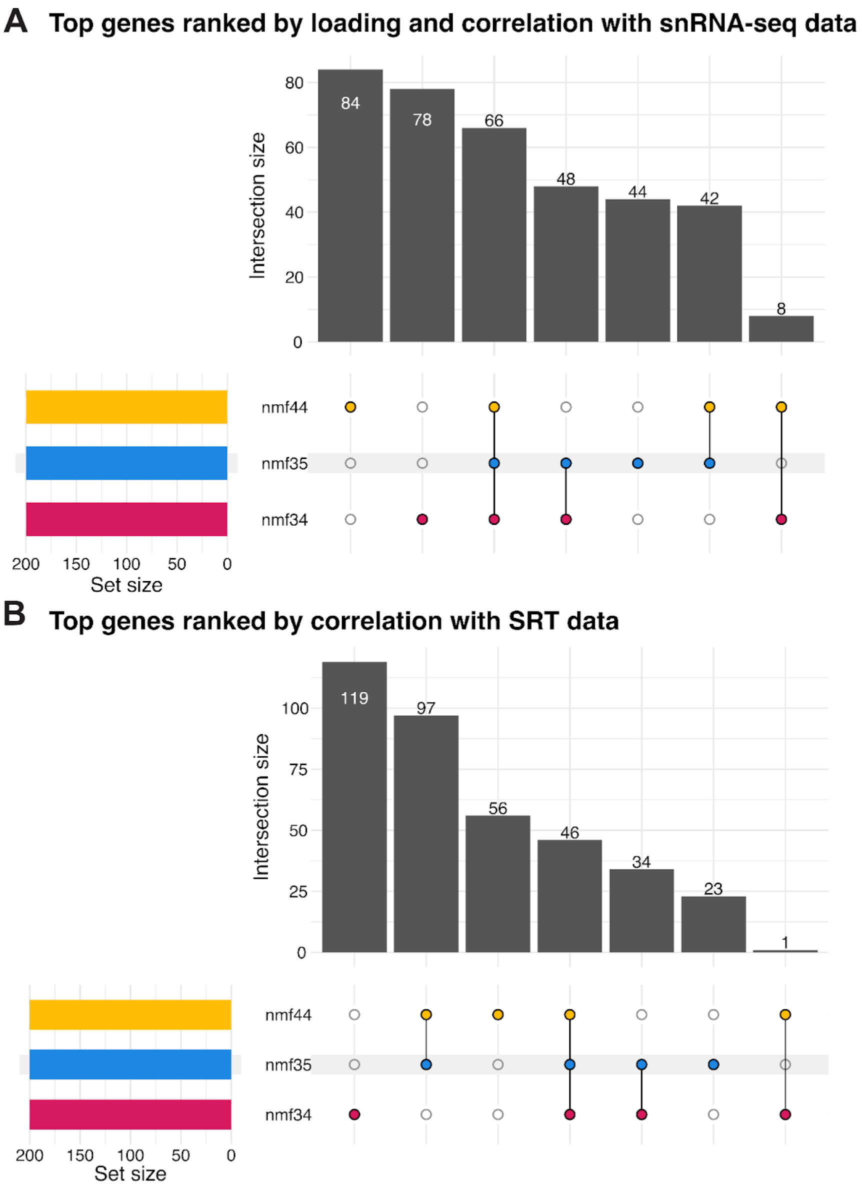


**Supplementary Figure. 35**. **Overlap among top-ranked genes for D1 island–associated NMF factors.** UpSet plots show intersections among the top 200 genes per factor (nmf34, nmf35, nmf44). Filled dots indicate which factors contribute to each intersection; bar height gives the intersection size; horizontal bars (left) give each factor's set size (all = 200). **(A)** Genes ranked in snRNA-seq by a joint criterion (high loading in $W$ and strong correlation between $H$ scores and gene expression). **(B)** Genes ranked in SRT by correlation between projected factor scores $H'$ and log-normalized spatial expression.

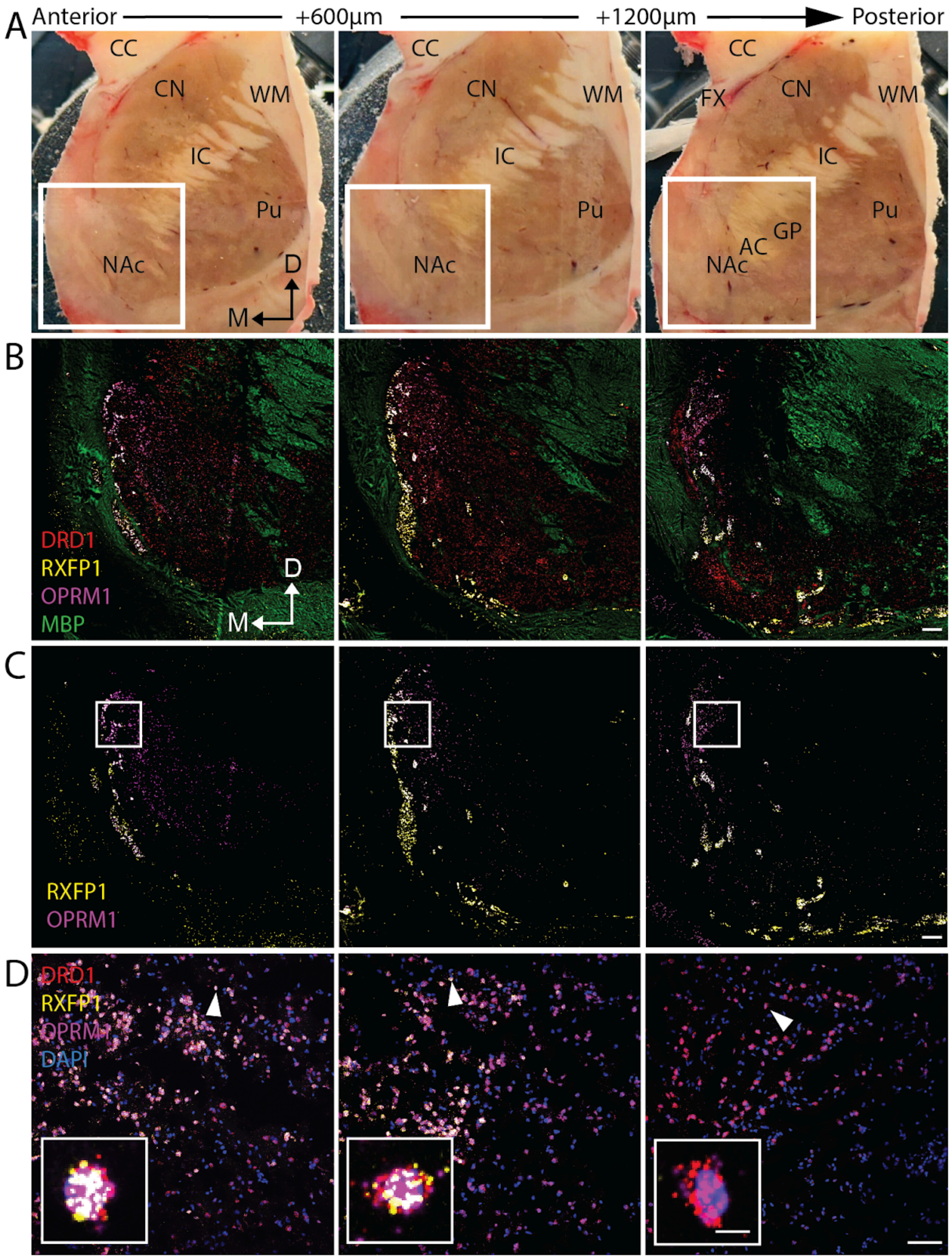


**Supplementary Figure. 36. smFISH validation of D1 island marker genes.**
**(A)** Tissue block from an independent donor (Br4032) sectioned at three depths on the A-P axis with 600 µm and 1200 µm intervals. AC - anterior commissure; CC - corpus callosum; CN - caudate nucleus; D - dorsal; FX - fornix; GP - globus pallidus; IC - internal capsule; M - medial; NAc - nucleus accumbens; ON - optic nerve; Pu - putamen; WM - white matter. **(B-C)** 2X smFISH images of tissue at anterior, intermediate, and posterior aspects of the NAc with D1 island marker genes *DRD1* (red), *RXFP1* (yellow), *OPRM1* (magenta), and white matter marker gene *MBP* (green). Individual staining for *RFXP1* and *OPRM1* is presented in **(C)** for clarity. Scale bar 1,000 µm. **(D)** 20X zoomed images of locations indicated by white boxes in **(C)**. Scale bar 100 µm. Insets depict a single neuron indicated by the white arrow. Scale bar of the inset is 5 µm.

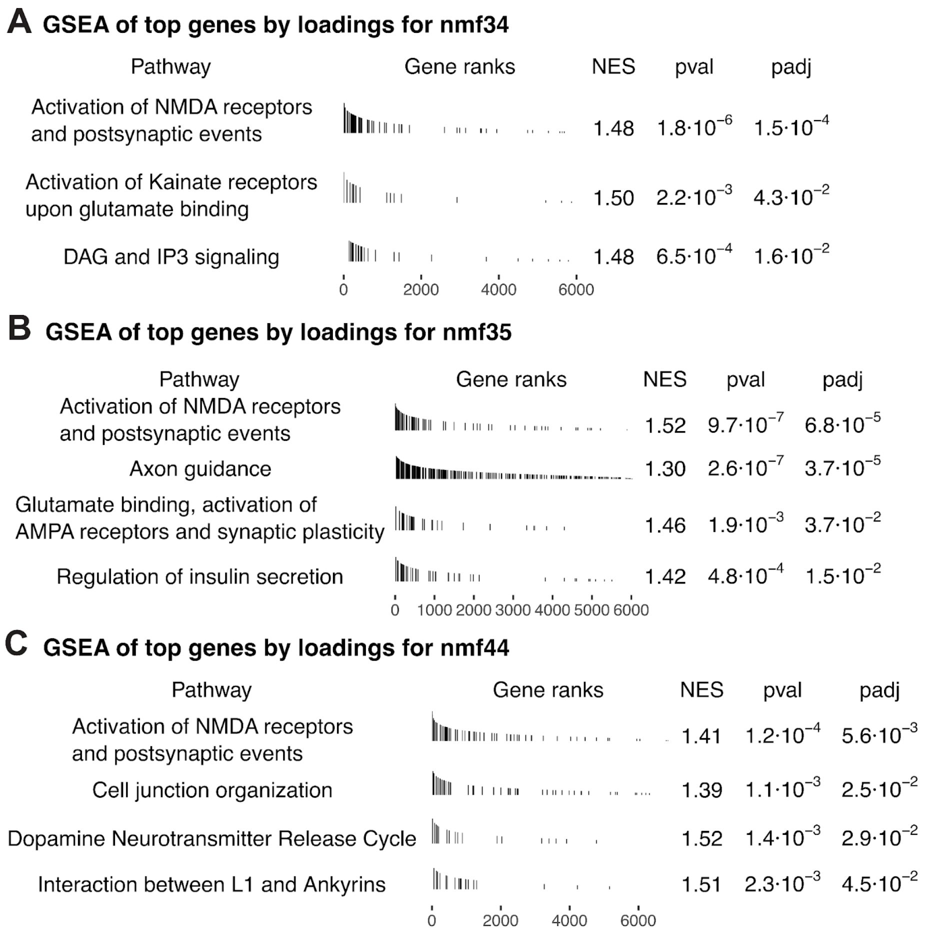


**Supplementary Figure. 37. GSEA of D1 island-associated NMF factors.** For each factor, pathways enriched among genes ranked by factor loadings are shown. Tick marks show gene ranks of pathway associated genes, and tables report normalized enrichment score (NES), nominal *p*-values, and Benjamini-Hochberg adjusted *p*-values. Significance was defined as adj. $p$-value < 0.05.

**(A)** Genes with high loadings in factor nmf34 are significantly enriched for “Activation of NMDA receptors and post-synaptic events”, “Activation of Kainate receptors upon glutamate binding”, “DAG and IP3 signaling” Reactome pathways.

**(B)** Genes with high loadings in factor nmf35 are significantly enriched for “Activation of NMDA receptors and postsynaptic events”, “Axon guidance”, and “Glutamate binding, activation of AMPA receptors and synaptic plasticity” Reactome pathways.

**(C)** Genes with high loadings in factor nmf44 are significantly enriched for “Activation of NMDA receptors and postsynaptic events”, “Cell junction organization”, “Dopamine neurotransmitter release cycle”, “Interaction between L1 and Ankyrins” Reactome pathways.

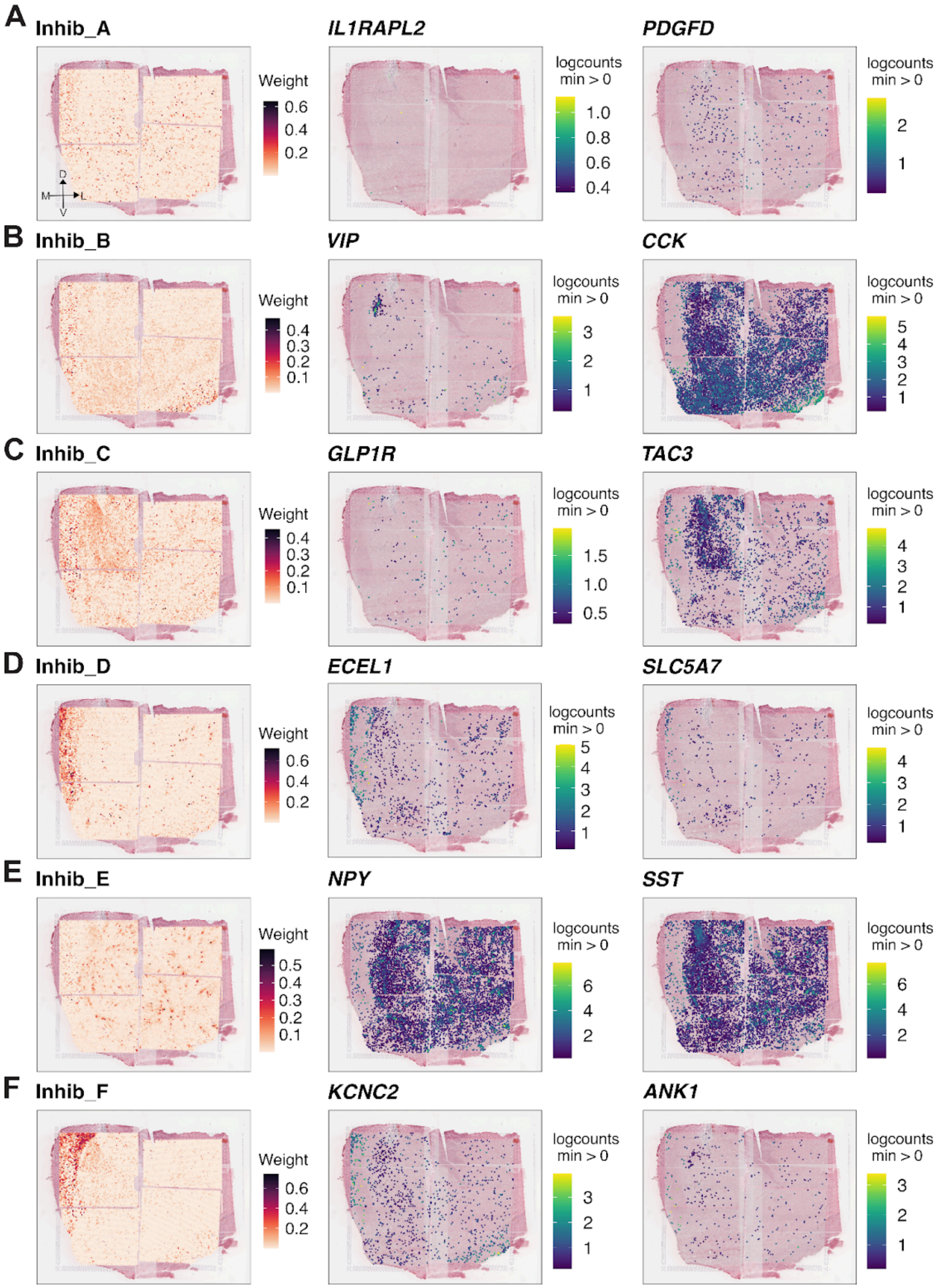


**Supplementary Figure. 38. Spatial localization of inhibitory neuron subtypes and their corresponding marker genes.** Spot plots showing *RCTD*-derived weights for six inhibitory neuronal populations overlaid on H&E in a representative donor (Br6522). All images are oriented with dorsal at the top and lateral to the right. For each subtype, adjacent panels display the spatial expression of two marker genes (log-normalized counts). **(A)** Inh_A; *IL1RAPL2*, *PDGFD*. **(B)** Inh_B; *VIP*, *CCK*. **(C)** Inh_C; *GLP1R*, *TAC3*. **(D)** Inh_D (cholinergic): *ECEL1*, *SLC5A7*. **(E)** Inh_E; *NPY*, *SST*. **(F)** Inh_F; *KCNC2*, *ANK1*. Patterns illustrate subtype-specific distributions across the NAc, with color bars indicating *RCTD* weight (left) and gene expression (right).

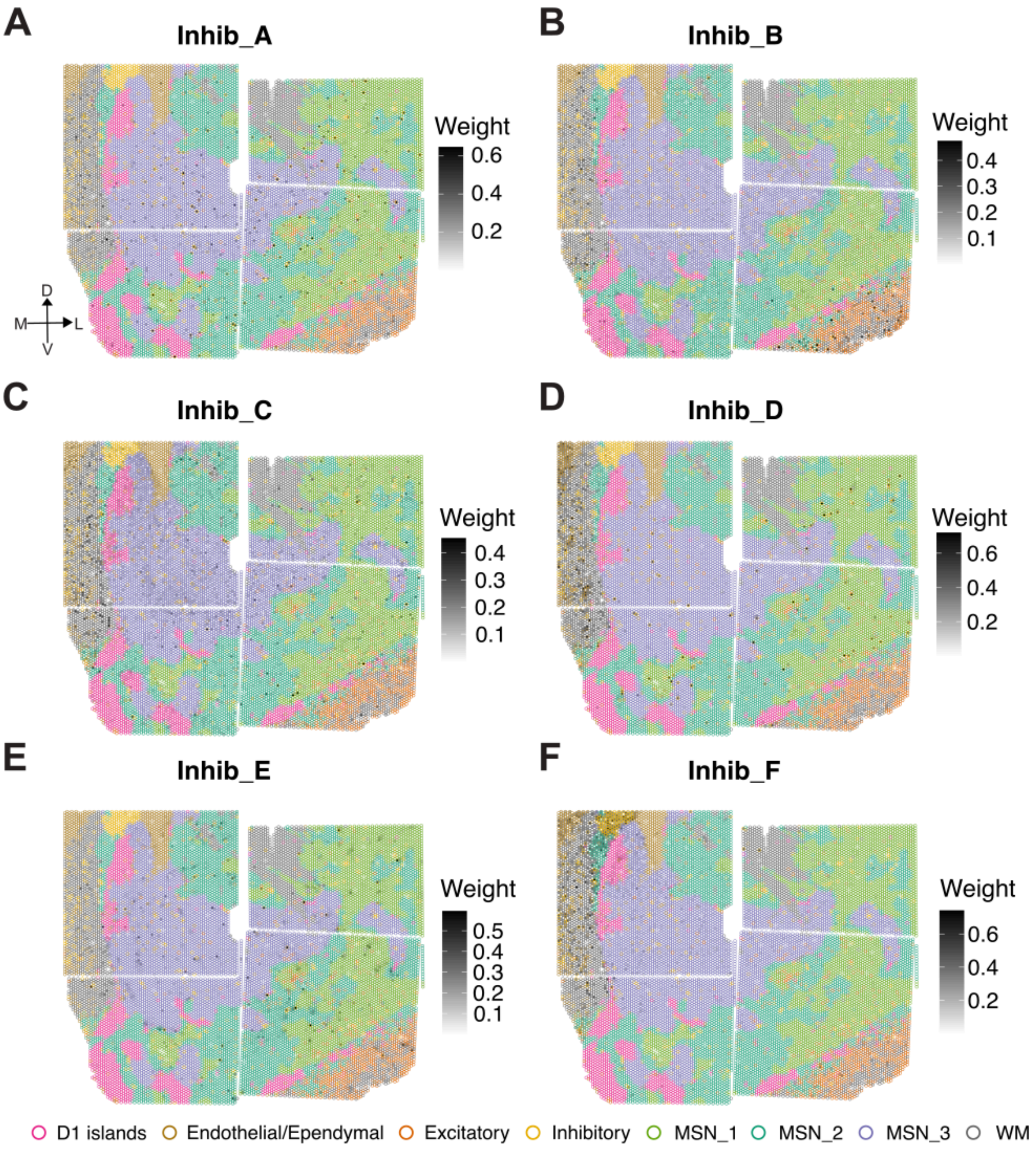


**Supplementary Figure. 39. Spatial mapping of inhibitory cell types across SpDs.** *escheR* [173] spot plots visualizing *RCTD*-derived weights corresponding to six inhibitory neuron subtypes in a representative donor (Br6522): **(A)** Inhib_A (*IL1RAPL2*+/*PDGFD*+), **(B)** Inhib_B (*VIP*+/*CCK*+) **(C)** Inhib_C (*GLP1R*+/*TAC3*+) **(D)** Inhib_D (*SLC5A7*+/*CHAT*+) **(E)** Inhib_E (*NPY*+/*SST*+/*CORT*+) **(F)** Inhib_F (*KCNC2*+/*ANK1*+) . Spot outlines are colored by SRT-derived SpDs and spot fills (grayscale) encode the corresponding *RCTD*-derived weights. Color bars indicate per-panel weight scales (darker = higher). All images are oriented with dorsal at the top and lateral to the right.

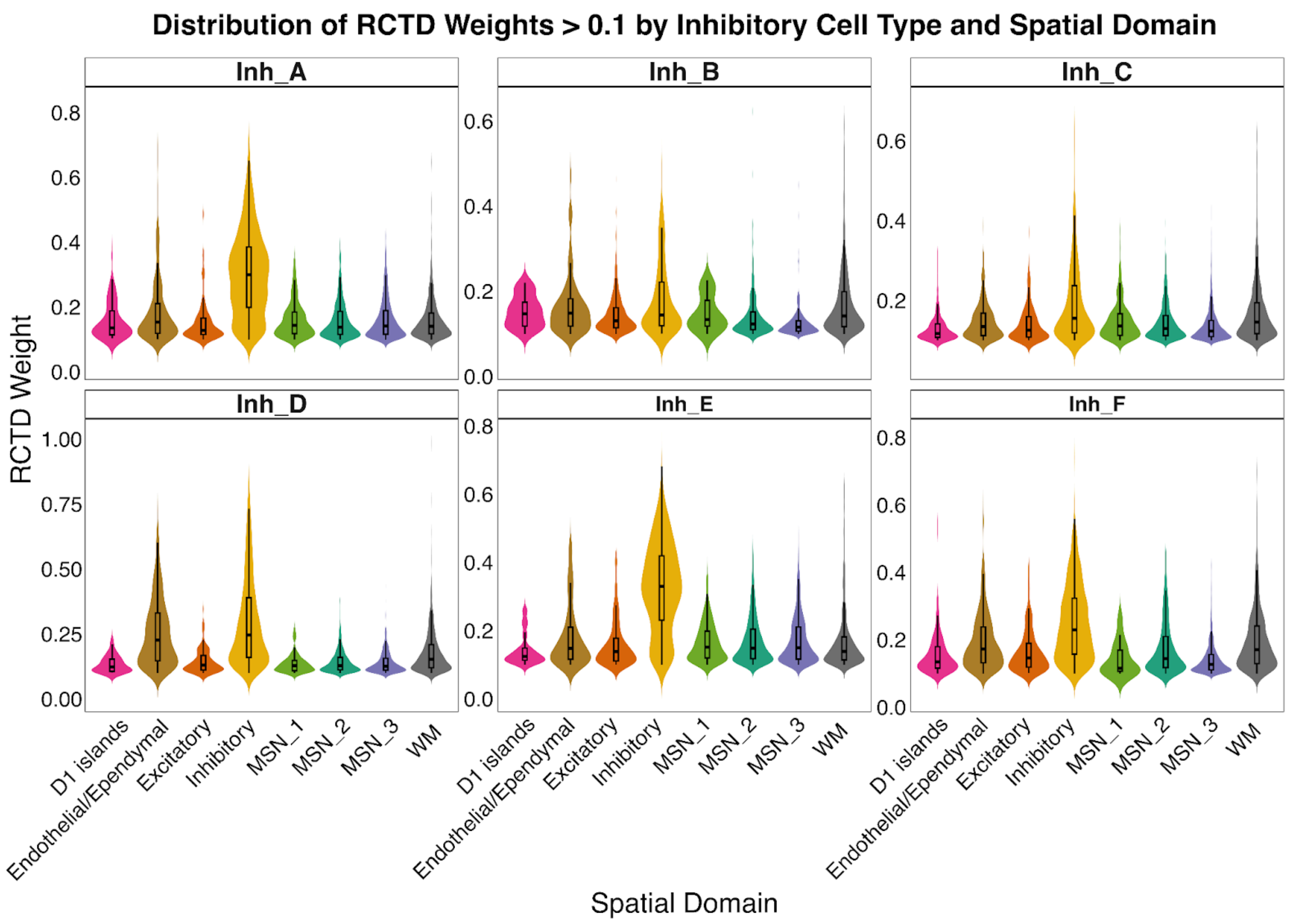


**Supplementary Figure. 40. Distribution of *RCTD*-derived weights for inhibitory cell types by SpD.** Violin plots showing *RCTD*-derived weights (y-axis) across SRT-derived SpDs (x-axis) after restricting to SRT spots in each SpD where the weight for the indicated inhibitory subtype exceeds 0.1. The inhibitory cell types visualized include **(A)** Inhib_A (*IL1RAPL2*+/*PDGFD*+), **(B)** Inhib_B (*VIP*+/*CCK*+), **(C)** Inhib_C (*GLP1R*+/*TAC3*+), **(D)** Inhib_D (*SLC5A7*+/*CHAT*+), **(E)** Inhib_E (*NPY*+/*SST*+/*CORT*+), **(F)** Inhib_F (*KCNC2*+/*ANK1*+).

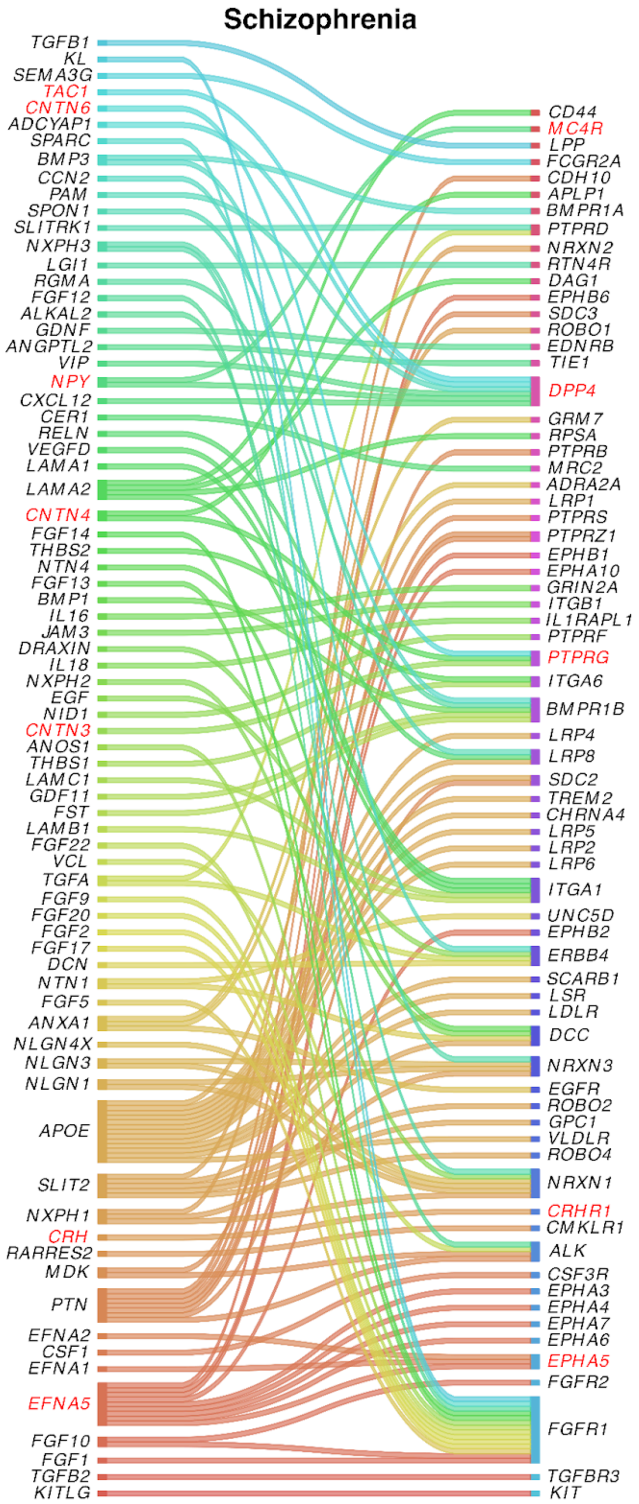


**Supplementary Figure. 41. Schizophrenia (SCZ)-relevant ligand-receptor (LR) signaling identified in the human NAc.** Sankey plots summarizing LR interactions in which at least one interacting partner is a SCZ risk gene prioritized by *OpenTargets*. Interactions were independently identified by applying *LIANA* to snRNA-seq data, and retained only if the same directed interaction appears in *OmniPath*, and at least one partner is a SCZ risk gene. Red labels indicate LR pairs that occur across multiple neuropsychiatric traits including *TAC1→MC4R*, *NPY→DPP4*, *CRH→CRHR1*, *CNTN*3/4/6→*PTPRG*, and a previously identified LR interaction with known SCZ relevance *EFNA5→EPHA5*.

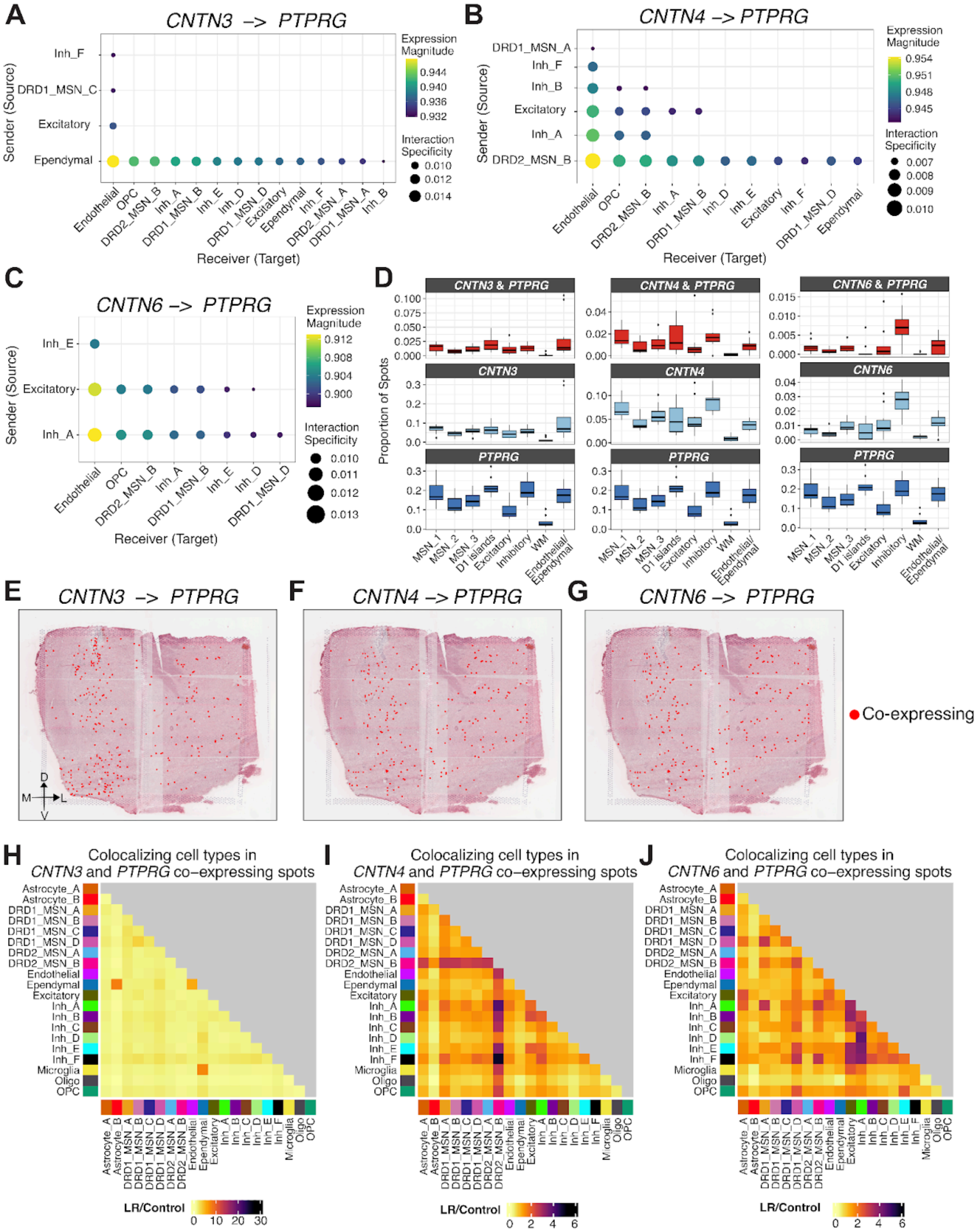


**Supplementary Figure. 42. Cell type and spatial context of CNTN3/4/6-PTPRG signaling.** Dotplots summarizing *LIANA* predictions of interacting cell types for high-confidence LR interactions (consensus rank ≤ 0.01) involving (**A**) *CNTN3→PTPRG*, (**B**) *CNTN4→PTPRG*, and (**C**) CNTN6→PTPRG. Dot size encodes the interaction specificity (natmi.edge_specificity) and dot color represents expression magnitude (`sca.LRscore`) for corresponding sender cell type (y-axis) and receiver cell type (x-axis) pair. (**D**) Boxplots summarizing the proportion of SRT spots (y-axis) in each SpD (x-axis) that co-express ligand and receptor (top), express only the ligand (middle), and express only the receptor (bottom). (**E-G**) Spot plots marking SRT spots that co-express the ligand and receptor (red) overlaid on H&E of a representative donor (Br6522). All images are oriented with dorsal at the top and lateral to the right. (**H-J**) Heatmaps showing the cell type co-localization in co-expressing spots relative to control spots, computed from *RCTD*-derived weights. Values depict the enrichment ratio $R$, where $R > 1$ indicates cell-type pairs that co-occur more often in co-expressing spots.

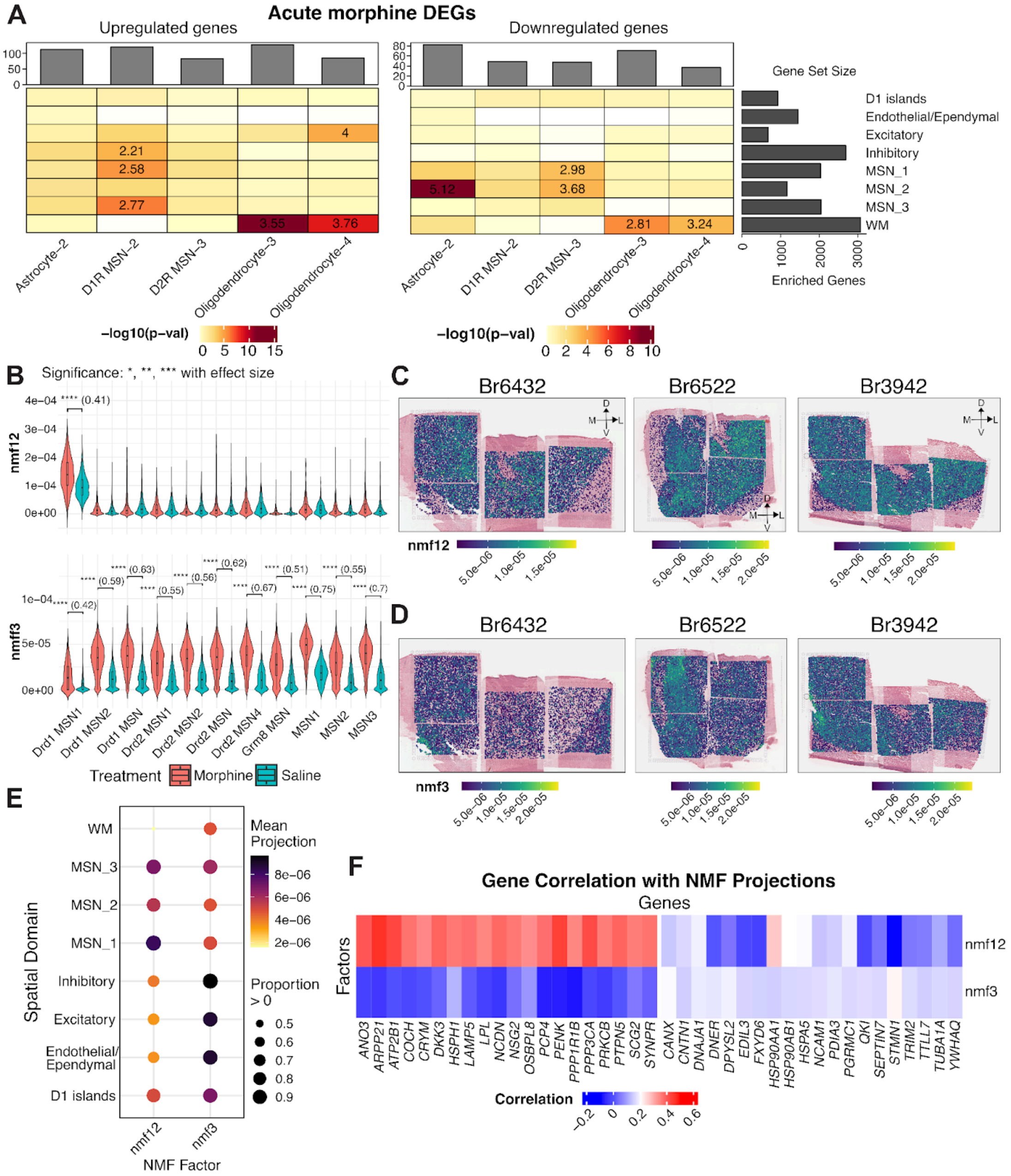


**Supplementary Figure. 43. Spatial mapping of response to acute morphine. (A)** Enrichment of differentially expressed genes (DEGs) from rodent acute morphine vs. saline controls [27] across human NAc spatial domains (SpDs). Up- and down-regulated gene sets from individual rodent cell types were tested separately. Heatmap visualizes the -$\log_{10}$(*p*-value) and select odds ratio (OR) for significant tests are included. (**B**) Violin plots comparing the factor weights for two acute-morphine associated factors inferred from MSN-restricted rodent snRNA-seq data. Weights (y-axis) are shown across cell types (x-axis) stratified by morphine and saline exposure. nmf12 is additionally specific to a single cell type Drd1 MSN 1, while nmf3 exhibits high weights across multiple MSN subtypes. Spot plots visualizing (**C**) nmf12 and (**D**) nmf3 projections across three representative donors: Br6432 (anterior), Br6522 (intermediate), and Br3942 (posterior). All images are oriented with dorsal at the top and lateral to the right. (**E**) Dotplot summarizing factor projections by SpD. Dot size encodes the proportion of SRT spots within a particular SpD which has non-zero projection weights, and dot color represents the mean projection weight. (**F**) Heatmap showing the correlation between SRT projection weights and log-normalized gene expression for the top 20 most correlated genes for morphine-associated factors nmf3 and nmf12. nmf12 is associated with modulators of dopaminergic signaling (*PENK*, *PRKCB*, *PTPN5*) while nmf3 is associated with genes corresponding to stress and metabolic response (*HSP90AA1*, *DNAJA1*).

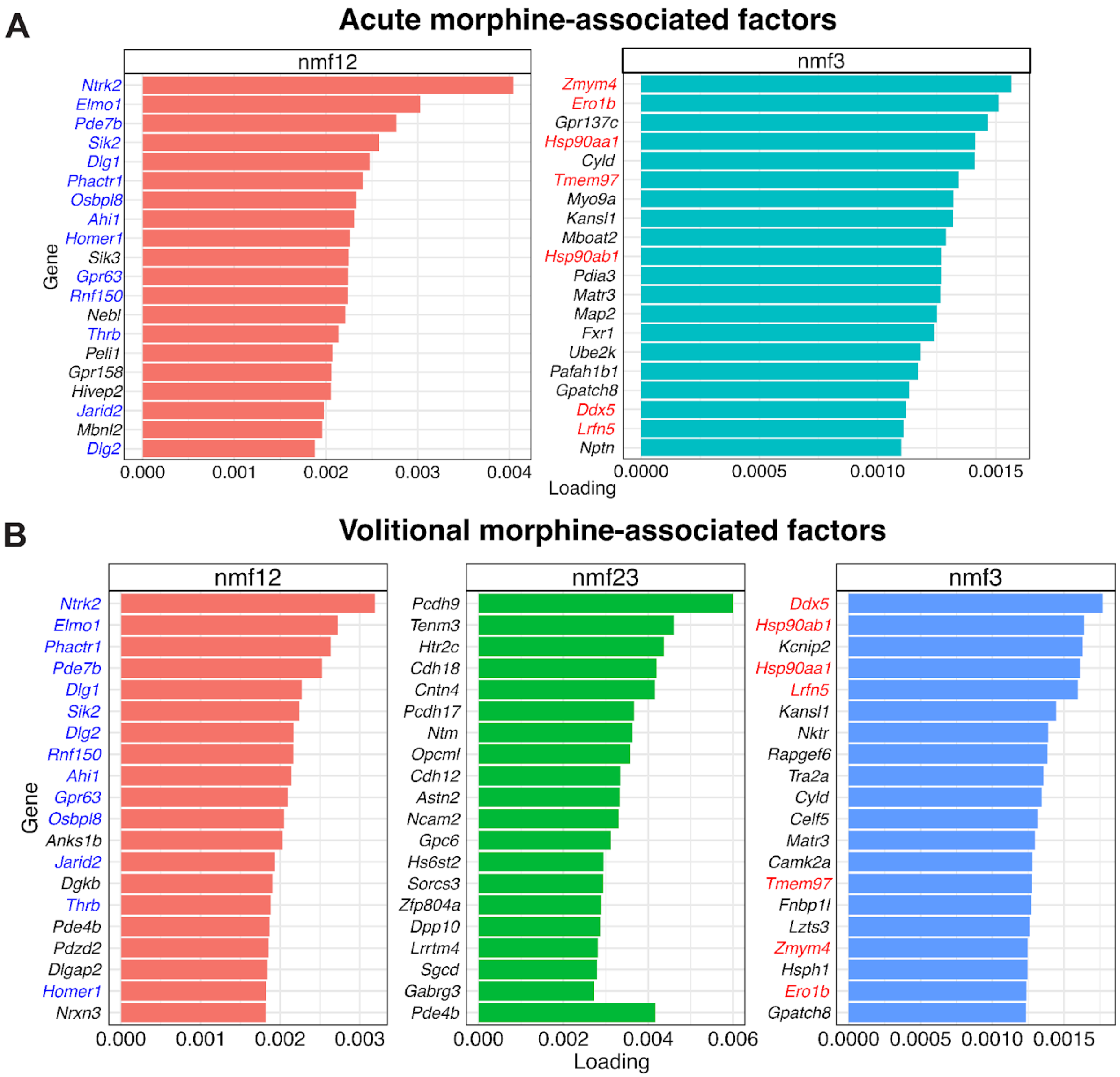


**Supplementary Figure. 44. Top rodent genes defining morphine-associated NMF factors. (A)** Bar plot showing gene loadings (x-axis) of top 20 rodent genes ranked by loading for factors nmf12 and nmf3 derived from MSN-restricted snRNA-seq data corresponding to acute morphine and matched saline controls [27]. (**B**) Bar plot showing gene loadings (x-axis) of top 20 rodent genes ranked by loading for factors nmf12, nmf23 and nmf3 derived from MSN-restricted snRNA-seq data corresponding to chronic morphine self-administration (volitional morphine) and matched saline controls.

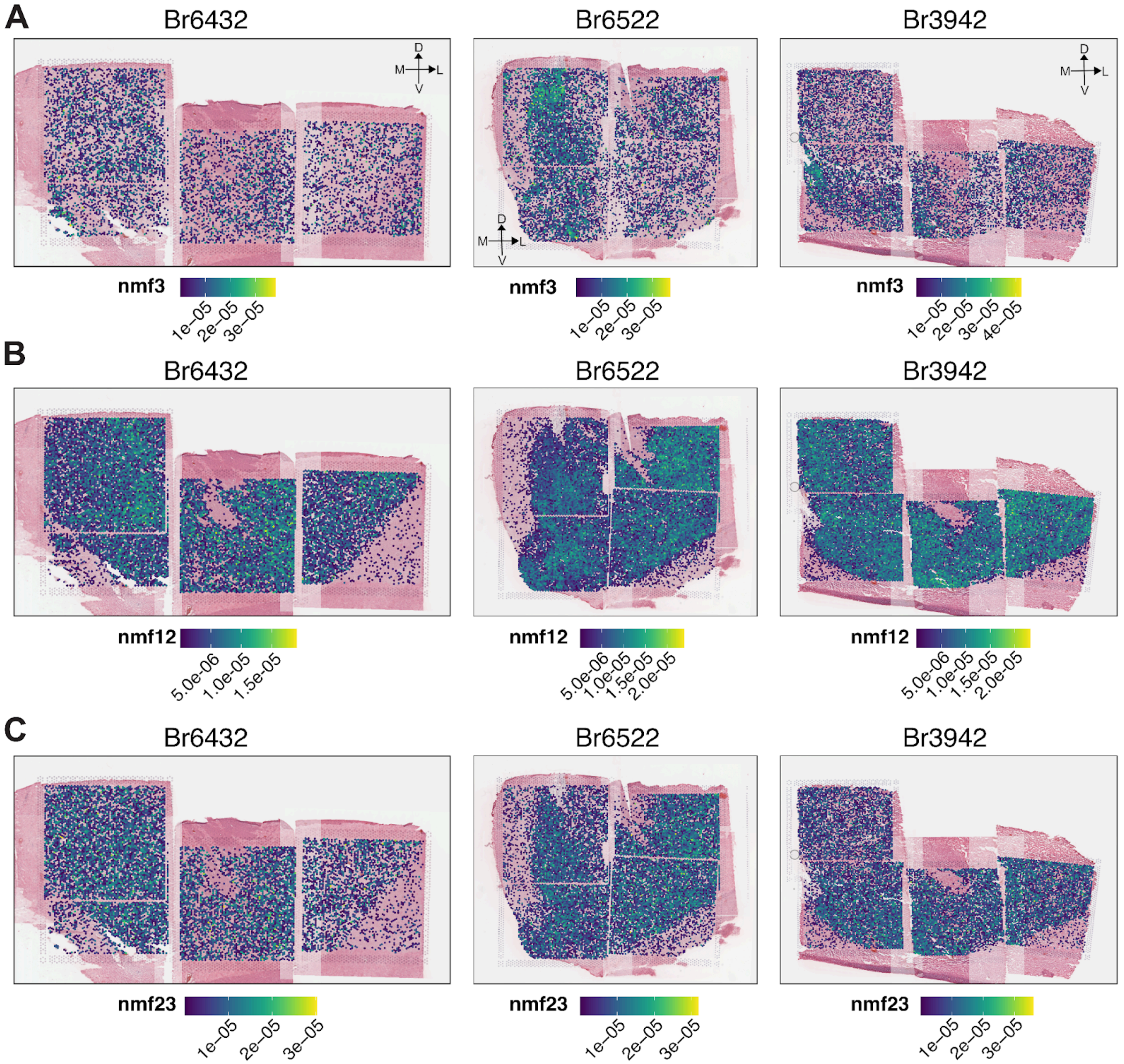


**Supplementary Figure. 45. Spatial projections of volitional morphine-associated factors in the human NAc.** Spot plots depicting the projections of NMF factors derived from MSN-restricted rodent snRNA-seq data for animals self-administering morphine and saline controls [27], shown for three representative donors: Br6432 (anterior), Br6522 (intermediate), and Br3942 (posterior). Panels correspond to distinct NMF factors including: (**A**) nmf3, (**B**) nmf12, and (**C**) nmf23. Per-panel color bars indicate the scale of the per-spot projection weight. All images are oriented with dorsal at the top and lateral to the right.

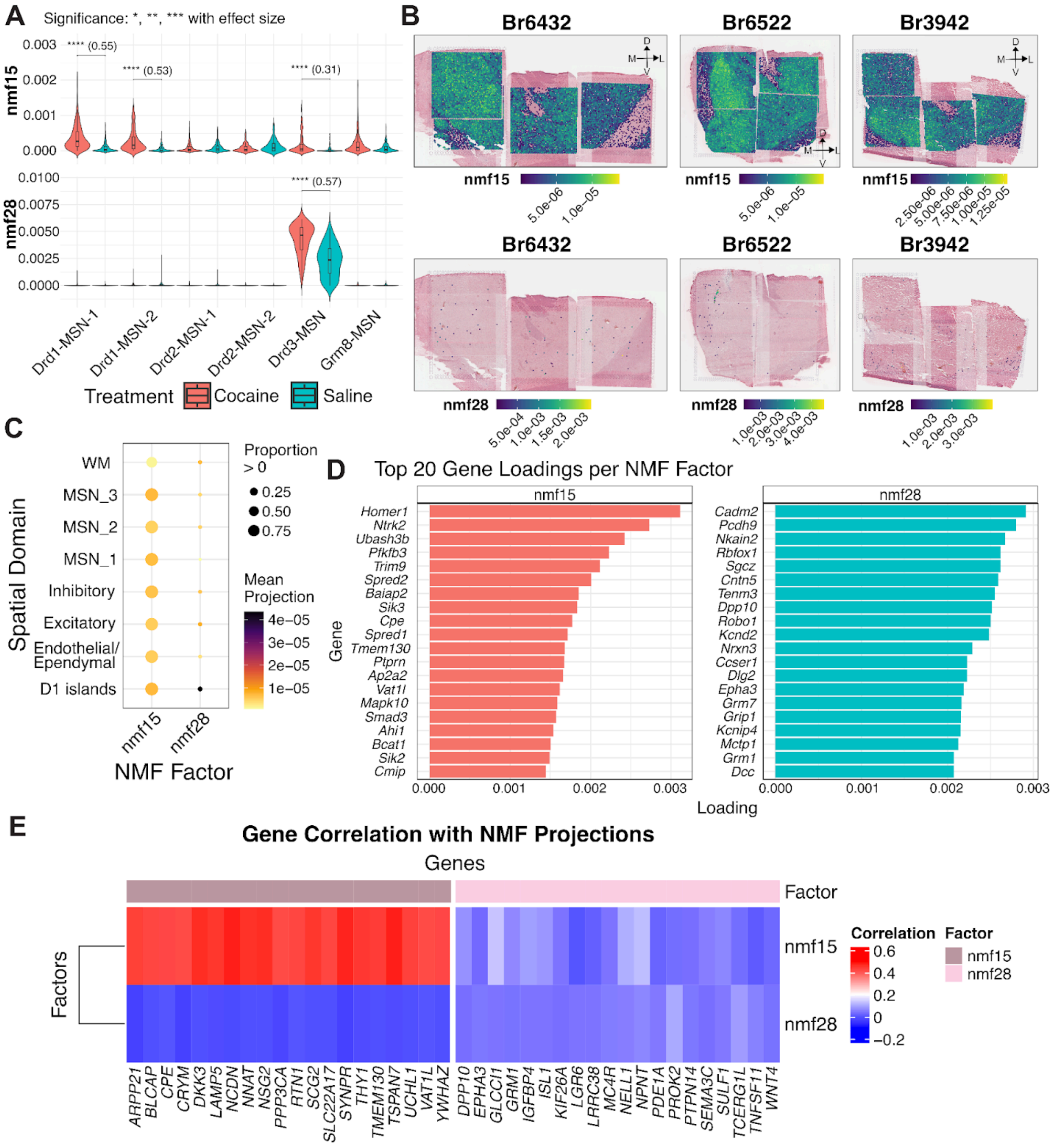


**Supplementary Figure. 46. Spatial mapping of response to acute cocaine. (A)** Violin plots comparing weights (y-axis) for acute cocaine-associated NMF factors (nmf15, nmf28) inferred from MSN-restricted rodent snRNA-seq data across MSN cell types (x-axis) stratified by cocaine vs. saline exposed cells. **(B)** Spatial projections of nmf15 (top) and nmf28 (bottom) across three representative donors: Br6432 (anterior), Br6522 (intermediate), Br3942 (posterior). All images are oriented with dorsal at the top and lateral to the right. **(C)** Dotplot summarizing factor projection weights by SRT-derived SpDs. Dot size encodes the proportion of SRT spots within a particular SpD which has non-zero projection weights, and dot color represents the mean projection weight. **(D)** Barplot showing the factor loadings (x-axis) corresponding to the top 20 rodent genes (y-axis) ranked by loadings for acute cocaine-associated factors nmf15 (left) and nmf28 (right). **(E)** Heatmap showing the correlation between SRT projection weights and log-normalized gene expression for the top 20 most correlated genes for acute cocaine-associated factors nmf15 and nmf28. nmf15 is associated with genes including *ARPP21*, *CRYM*, *RTN1*, and *SYNPR*, while nmf28 is associated with *GRM1* (metabotropic glutamate receptor), *MC4R*, and *PDE1A* which play a role in reward processing.

### Supplementary Tables

**Supplementary Table. 1**. **Donor demographic information.** Demographic information on brain donors used for transcriptomic profiling, including Donor ID, age at time of death, sex, psychiatric diagnosis, post-mortem interval (PMI), body mass index (BMI, calculated postmortem), screening RNA Integrity Number (RIN) from prefrontal cortex (PFC) calculated at time of brain collection.

**Supplementary Table. 2**. **snRNA-seq DEGs by cluster**. Differential expression testing was performed using 2 separate methods. DEGs were calculated using a 1-versus-all-others method via `findMarkers_1vALL()` from *DeconvoBuddies*[129] package or using a pairwise method testing with the `findMarkers()` function from *scran*. For 1vALL testing, the logFC column represents the standardized log-fold change. For pairwise testing, the logFC column represents the $\log_2$ fold change of the comparison with the largest *p*-value.

**Supplementary Table. 3 Stratified LD score regression (sLDSC) results for cell type-specific annotations.** For each of the 20 annotated cell types, the top 10% of genes by relative expression were selected to derive variant annotations, which were used to assess the enrichment of trait heritability in each cell type using sLDSC. Columns correspond to the cell type, trait, the proportion of SNPs included in the annotation (*Prop._SNPs*), the proportion of total SNP heritability explained (*Prop._h2*) with its standard error, the enrichment statistic with its standard error and nominal *p*-value, and the regression coefficient with its standard error and *z*-score. The *p_zscore* column provides the two-sided *p*-value corresponding to the coefficient *z*-score, and the FDR column gives Benjamini–Hochberg adjusted *p*-values across tests.

**Supplementary Table. 4. Summary of *nnSVG* results across donors with and without controlling for *PRECAST* (k=2) clusters**. SVGs were identified with *nnSVG*[51] in individual samples and the following metrics were estimated for each gene detected as a spatially variable gene (SVG) in at least one sample, including (i) the proportion of samples in which the gene was significantly identified to be a SVG (FDR < 0.05), (ii) the proportion of samples in which the gene was among the top 100 SVGs. (iii) mean *nnSVG* rank across all samples, and (iv) rank of the mean *nnSVG* rank among all genes. The control covariate indicates whether *PRECAST* k = 2 cluster assignments were used to account for broad transcriptional differences between white matter and gray matter when nnSVGs were inferred. The top 2,000 SVGs based on the mean *nnSVG* rank were used to infer *PRECAST* clusters. No covariates were adjusted to infer *PRECAST* k = 2 cluster assignments, while *PRECAST* (k ≥ 3) clusters were inferred using nnSVGs while controlling for *PRECAST* k = 2 cluster assignments.

**Supplementary Table. 5. Significant DEGs corresponding to unmerged *PRECAST* k= 10 clusters.** Differential expression testing was performed on pseudobulk samples aggregated by *PRECAST* (k= 10) cluster assignments and capture area. Cluster assignments from random start 3 were used. Results are provided for three tests: one-vs-all enrichment, pairwise clustering contrasts, and ANOVA for significantly differentially expressed genes (FDR < 0.05). All tests were performed adjusting for donor sex and slide number.

**Supplementary Table. 6. Significant DEGs corresponding to SRT-derived spatial domains (SpDs).** Differential expression testing was performed on pseudobulk samples aggregated by SpD and capture area. SpDs were obtained by merging *PRECAST* k =10 clusters 2 with 9 and 6 with 10. Results are provided for three tests: one-vs-all enrichment, pairwise clustering contrasts, and ANOVA for significantly differentially expressed genes (FDR < 0.05). All tests were performed adjusting for donor sex and slide number.

**Supplementary Table. 7. Stratified LD score regression (sLDSC) results for SpD-specific annotations.** For each spatial domain (SpD), the top 15% of genes by relative expression were selected to derive variant annotations, which were used to assess the enrichment of trait heritability in each domain using sLDSC. Columns correspond to SpD, trait, the proportion of SNPs included in the annotation (*Prop._SNPs*), the proportion of total SNP heritability explained (*Prop._h2*) with its standard error, the enrichment statistic with its standard error and nominal *p*-value, and the regression coefficient with its standard error and *z*-score. The *p_zscore* column provides the two-sided *p*-value corresponding to the coefficient *z*-score, and the FDR column gives Benjamini–Hochberg adjusted *p*-values across tests.

**Supplementary Table. 8. Genes correlated with MERINGUE consensus patterns (MCPs).** For each MCP, Pearson correlations were computed between pattern scores and spatial gene expression. Table columns correspond to the top 100 genes ranked by correlation strength, the associated MCP, the correlation coefficient, and the within-MCP rank.

**Supplementary Table. 9. Genes correlated with MSN-associated NMF factors.** For each MSN associated NMF factor, including nmf3, nmf4, nmf7, nmf10, and nmf39, Pearson correlations were computed between factor projection scores and spatial gene expression. Table columns correspond to the top 200 genes ranked by correlation strength, the associated NMF factor, the correlation coefficient, and the within-factor rank.

**Supplementary Table. 10. Gene set enrichment analysis (GSEA) of MSN-associated NMF factors.** GSEA was performed using the *fgsea* package with Reactome pathway annotations on genes ranked by NMF weights (W) for factors nmf3, nmf4, nmf7, nmf10, and nmf39. Table columns correspond to the pathway, nominal enrichment *p*-value (*pval*), BH-adjusted *p*-value (*padj*), expected error of the *p*-value logarithm (*log2err*), enrichment score (*ES*), normalized enrichment score (*NES*), and pathway size after removing genes not present in the ranked list.

**Supplementary Table. 11 Stratified LD score regression (sLDSC) results for SpD-specific annotations.** For each NMF factor, factor-specific gene sets were used to construct SNP annotations, which were then tested for enrichment of trait heritability using sLDSC. Columns correspond to NMF factor, trait, the proportion of SNPs included in the annotation (*Prop._SNPs*), the proportion of total SNP heritability explained (*Prop._h2*) with its standard error, the enrichment statistic with its standard error and nominal *p*-value, and the regression coefficient with its standard error and *z*-score. The *p_zscore* column provides the two-sided *p*-value corresponding to the coefficient *z*-score, and the FDR column gives Benjamini–Hochberg adjusted *p*-values across tests.

**Supplementary Table. 12 Genes correlated with D1 islands-associated NMF factors.** For each D1 islands associated NMF factor, including nmf34, nmf35, nmf44, Pearson correlations were computed between factor projection scores and spatial gene expression. Table columns correspond to the top 200 genes ranked by correlation strength, the associated NMF factor, the correlation coefficient, and the within-factor rank.

**Supplementary Table. 13 Gene set enrichment analysis (GSEA) of D1 islands-associated NMF factors.** GSEA was performed using the *fgsea* package with Reactome pathway annotations on genes ranked by NMF weights (W) for factors nmf34, nmf35, nmf44. Table columns correspond to the pathway, nominal enrichment *p*-value (*pval*), BH-adjusted *p*-value (*padj*), expected error of the *p*-value logarithm (*log2err*), enrichment score (*ES*), normalized enrichment score (*NES*), and pathway size after removing genes not present in the ranked list.

**Supplementary Table. 14. Trait-relevant ligand-receptor (LR) interactions.** LR interactions identified in OmniPath and predicted by LIANA in which either the ligand or receptor is a risk gene for anxiety, depression, schizophrenia, or substance dependence.

**Supplementary Table. 15. Top rodent genes for drug-associated NMF factors.** For each NMF factor associated with chronic volitional morphine (nmf3, nmf12, nmf23), acute morphine (nmf3, nmf12), and acute cocaine (nmf15, nmf28), the top 100 rodent genes are ranked by their factor loadings, with higher values indicating stronger association with the corresponding factor.

**Supplementary Table. 16. Genes correlated with drug-associated NMF factors.** For each NMF factor, associated with chronic volitional morphine (nmf3, nmf12, nmf23), acute morphine (nmf3, nmf12), and acute cocaine (nmf15, nmf28), Pearson correlations were computed between factor projection scores and spatial gene expression. Table columns correspond to the top 200 genes ranked by correlation strength, the associated NMF factor, the correlation coefficient, and the within-factor rank.
